# Upregulation of Inhibitor of DNA Binding 1 and 3 is Important for Efficient Thermogenic Response in Human Adipocytes

**DOI:** 10.1101/2024.06.28.601234

**Authors:** Rini Arianti, Boglárka Ágnes Vinnai, Rahaf Alrifai, Gyath Karadsheh, Yousif Qais Al-Khafaji, Szilárd Póliska, Ferenc Győry, László Fésüs, Endre Kristóf

## Abstract

Brown and beige adipocytes can be activated by β-adrenergic agonist via cAMP-dependent signaling. Performing RNA-sequencing analysis in human cervical area-derived adipocytes, we found that dibutyryl-cAMP, which can mimic *in vivo* stimulation of browning and thermogenesis, enhanced the expression of browning and batokine genes and upregulated several signaling pathway genes linked to thermogenesis. We observed that the expression of ‘Inhibitor of DNA binding and cell differentiation’ (ID) 1 and particularly ID3 was strongly induced by the adrenergic stimulation. The degradation of ID1 and ID3 elicited by the ID antagonist AGX51 during thermogenic activation prevented the induction of proton leak respiration that reflects thermogenesis and abrogated cAMP analogue-stimulated upregulation of thermogenic genes and mitochondrial complex I, II, and IV subunits. The presented data suggests that ID proteins contribute to efficient thermogenic response of adipocytes during adrenergic stimulation.

**Graphical abstract:** 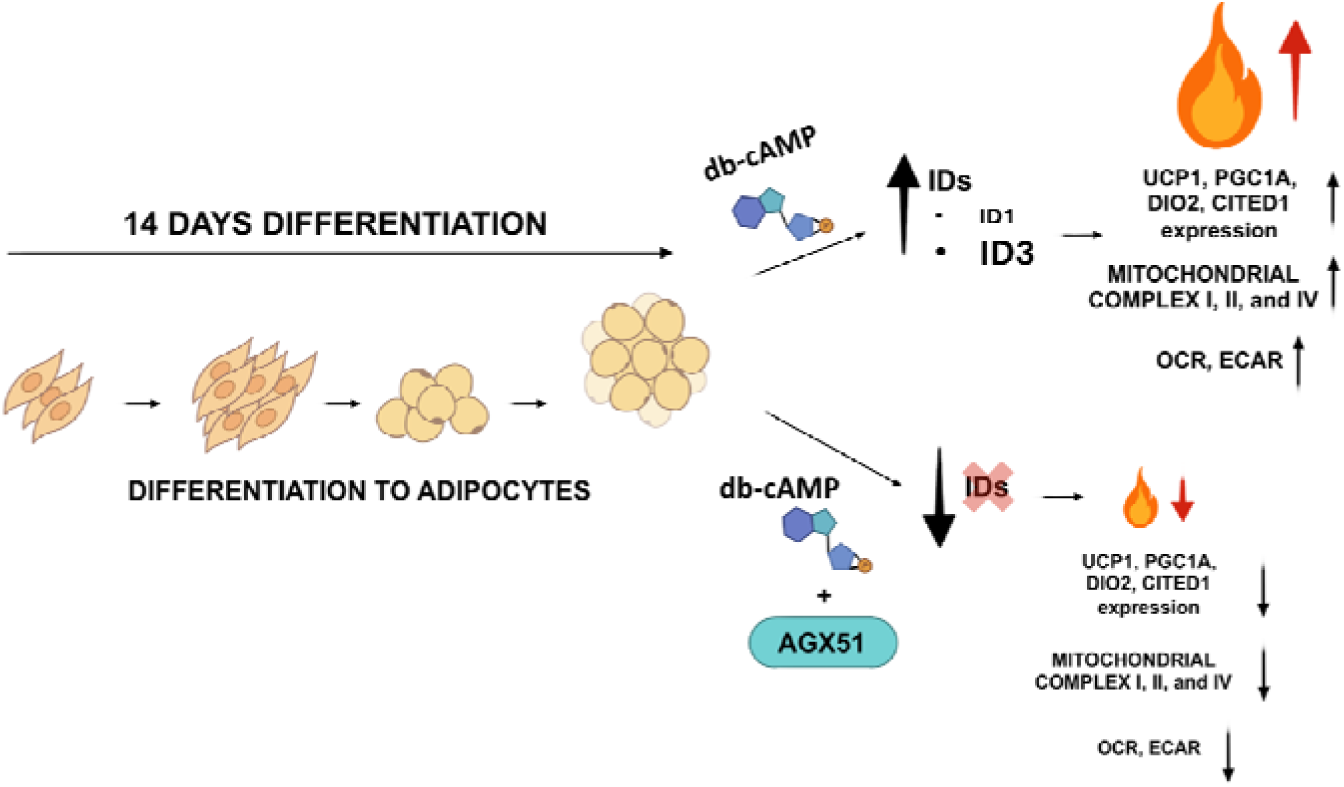

## Introduction

In humans, there are two main types of adipose tissue, white and brown (WAT and BAT). WAT is a critical regulator of systemic energy homeostasis as it acts as a calorie storage. In nutrient surplus conditions, WAT stores the excess in the form of neutral lipids, while in the case of nutrient deficit, it supplies fatty acids and glycerol to other tissues via lipolysis [1]. Anatomically, WAT consists of two main depots, subcutaneous (SC) and visceral WAT (VAT) which is located around the internal organs. In obesity, the pathologic accumulation of excess VAT is strongly associated with metabolic complications, such as insulin resistance and type 2 diabetes [2, 3]. White adipocytes, resident in WAT, are characterized by the presence of a single large (unilocular) lipid droplet and small number of mitochondria [4].

Mammals also possess BAT, which dissipates energy via non-shivering thermogenesis by the action of the two thermogenic types of adipocytes, classical brown and beige (also known as brite). Brown adipocytes possess multilocular lipid droplets and abundant mitochondria with dense cristae, expressing uncoupling protein 1 (UCP1) which uncouples the mitochondrial proton gradient from adenosine triphosphate (ATP) synthesis resulting in heat release [5]. Beige adipocytes are referred as an inducible form of thermogenic fat cells which are sparsely distributed within several WAT depots. Due to the decay of BAT amount shortly after birth, it has been considered functionally insignificant in adults for a long period of time. However, numerous studies utilizing positron emission tomography (PET) have provided evidence to the contrary, showing that human adults indeed possess significant amounts of BAT depots which can be stimulated to generate heat by cold exposure [6, 7]. A more recent study has refined the PET-computed tomography (CT) method to precisely identify brown/beige adipose depots in humans. Using this improved technique, Leitner *et al.* successfully mapped the distribution of BAT throughout the entire body and estimated its thermogenic capacity. BAT and brownable depots were found dispersed in various areas, including the cervical, supraclavicular, axillary, mediastinal, paraspinal, and abdominal regions [8].

Inhibitors of DNA binding and cell differentiation (ID) proteins are classified within the helix-loop-helix (HLH) transcription factor family; however, unlike other members of this family, they do not possess a DNA binding motif. They act as inhibitors of basic HLH transcription factors, thereby negatively controlling cell type-specific gene expression. Their significance lies in their involvement in development processes, where they play a crucial role in regulating cell-cycle progression, cell proliferation, cell differentiation, cell fate determination, hematopoiesis, angiogenesis, and metabolic adaptation of cancer cells [9, 10]. This is achieved through the modulation of various cell-cycle regulators using both direct and indirect interactions [11]. Although ID proteins contain a highly conserved HLH domain, they display significant sequence divergence. Targets of ID proteins include E, Ets, and Paired Box proteins, Retinoblastoma, p107, p130, MyoD, and Myf-5 [12, 13].

Browning of WAT and the activation of brown and beige adipocytes can be triggered by external cues, such cold exposure, physical exercise, or β-adrenergic agonists [5]. Subsequently, cyclic adenosine monophosphate (cAMP) activates protein kinase A (PKA) that phosphorylates a variety of downstream targets, including transcription (co)factors to upregulate thermogenic gene expression. In our experiments, we treated human cervical area-derived adipocytes with dibutyryl (db)-cAMP, which mimics adrenergic induction of *in vivo* browning and thermogenesis [14], and analyzed the global transcriptomic response by RNA-sequencing. In accordance with the elevation of thermogenic genes [15–17], such as *UCP1*, *PGC1a*, *DIO2*, and *CITED1*, we found that brown adipocyte content and browning capacity were significantly increased by adrenergic stimulation in both SC and deep neck (DN)-area derived cervical adipocytes. We observed that genes of several signaling pathways related to thermogenesis and the expression of ID proteins, especially ID3, were induced in these adipocytes. In order to reveal the functional significance of the upregulation of IDs, we applied the ID antagonist, AGX51 to inhibit ID family proteins during adrenergic stimulation [18]. It was found that cAMP analog-stimulated elevation of proton leak respiration, which reflects induced thermogenesis, of mitochondrial complex I and of thermogenic gene expression were hampered by AGX51, which promoted the degradation of ID1 and ID3. Our results suggest that induction of IDs, especially ID3, are necessary for the efficient thermogenic response during adrenergic stimulation in human adipocytes.

## Results

### Adrenergic stimulation increased thermogenic capacity and batokine secretion in human cervical area-derived adipocytes

To explore the gene expression changes in adipocytes derived from human SC or DN progenitors upon adrenergic stimulation by db-cAMP, global RNA-sequencing analysis was carried out. The expression of general adipocyte markers, such as *GLUT4*, *FABP4*, *LPL*, *AGPAT*, *ADIPOQ*, *PLIN1*, *LEPR*, and *PPARG* [19] was similar between untreated and db-cAMP-treated adipocytes derived from the two depots (Figure 1a). We analyzed the influence of db-cAMP on the browning capacity of adipocytes by utilizing the publicly available webtools based on the pattern of gene expression: BATLAS to quantify brown adipocyte content [20] and ProFAT to measure browning capacity [21]. In accordance with the high browning potential of cervical area-derived adipocytes [22] and our previous results [23], it was found that DN-derived adipocytes had higher BATLAS and ProFAT scores as compared to SC-derived ones (Figure 1b, c). Db-cAMP elevated BATLAS score significantly only in DN-derived adipocytes (Figure 1b), however, it increased ProFAT score in both cell types (Figure 1c). Db-cAMP-treated DN-derived adipocytes had the highest expression level of BATLAS brown marker genes as shown in the heatmap displayed in Supplementary Figure 1. The expression of ProFAT marker genes was elevated by cAMP-driven stimulation in both types of adipocytes (Supplementary Figure 2). The expression of the most characteristic brown adipocyte markers was increased in db-cAMP-treated adipocytes derived from both depots (Figure 1d), and the upregulation of *UCP1*, *PGC1a*, *DIO2*, and *CITED1* was validated by quantitative real time PCR (RT-qPCR) (Figure 1e).

**Figure 1.**
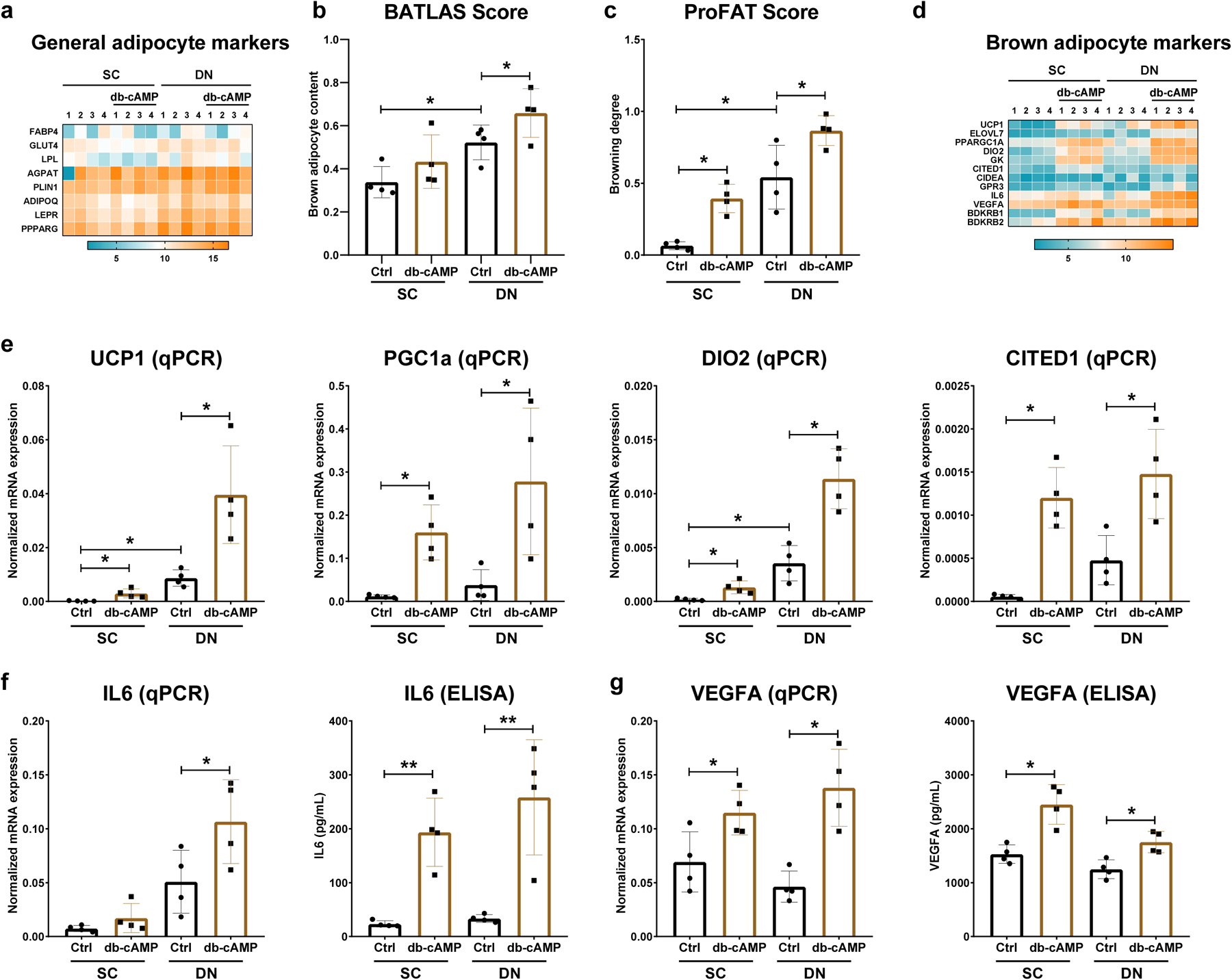
The expression of brown/beige markers in subcutaneous (SC) and deep neck (DN)-derived human adipocytes after 10 h of thermogenic activation driven by dibutyryl-cAMP (db-cAMP). (a) Heatmap displaying the gene expression pattern of adipocyte markers. (b) Brown adipocyte content quantified by BATLAS. (c) Browning capacity quantified by ProFAT. (d) Heatmap displaying the expression of brown/beige adipocyte markers. (e) mRNA expression of *UCP1*, *PGC1a*, *DIO2*, and *CITED1* analyzed by RT-qPCR. (f) mRNA expression (left panel) and secretion (right panel) of IL6 by *ex vivo* differentiated and activated adipocytes detected by RT-qPCR and ELISA, respectively. (g) mRNA expression (left panel) and secretion (right panel) of VEGFA by *ex vivo* differentiated and activated adipocytes detected by RT-qPCR and ELISA, respectively. VST scores from DESeq2 analysis were used to generate the heatmaps. n=4, statistical analysis was performed by one-way ANOVA followed by Tukey’s post-hoc test, *p<0.05.

RNA-sequencing data confirmed by RT-qPCR showed that the expression of interleukin-6 (*IL6*) and vascular endothelial growth factor A (*VEGFA*), which are known as brown adipocyte adipokines (batokines) [24, 25], were also elevated by db-cAMP in human cervical area-derived adipocytes (Figure 1d,f,g), leading to increased secretion of these batokines (Figure 6f, g, right panels) into the conditioned media. It could be concluded that the cell permeable cAMP analogue strongly increased the expression of thermogenic genes, browning potential, and batokine secretion in both SC and DN-derived adipocytes.

### Adrenergic stimulation promotes upregulation of several signaling pathways

As a next step, we identified 1527 (758 induced and 769 suppressed genes) and 1718 (927 induced and 791 suppressed genes) differentially expressed genes (DEGs) upon thermogenic activation in SC or DN-derived adipocytes, respectively (Figure 2a, Supplementary Tables 3-6). A total of 588 genes were commonly induced (Figure 2b, left panel) whereas 476 genes were commonly suppressed (Figure 2b, right panel) in both types of adipocytes upon cAMP analogue-driven stimulation in comparison to untreated ones. In addition to well characterized thermogenic genes (Figure 1), the expression of nuclear receptor subfamily 4 group A member 1 (*NR4A1*), G protein-coupled receptor class C group 5 member A (*GPRC5A*), interleukin 11 (*IL11*), long intergenic non-protein coding RNA 473 (*LINC00473*), and *ID3* were also strongly upregulated by db-cAMP in both SC and DN-derived adipocytes (Figure 2c). *NR4A1*, which encodes a nuclear receptor, was reported to be upregulated in brown adipocytes in response to β-adrenergic stimulation and in BAT of cold-exposed mice [26]. Regarding *ID* genes, *ID1* expression was induced only in SC, whereas *ID4* was upregulated only in DN-derived adipocytes (Figure 2c). In contrast to DN, in SC-derived adipocytes *ID4* expression was suppressed by db-cAMP (Figure 2c).

**Figure 2.**
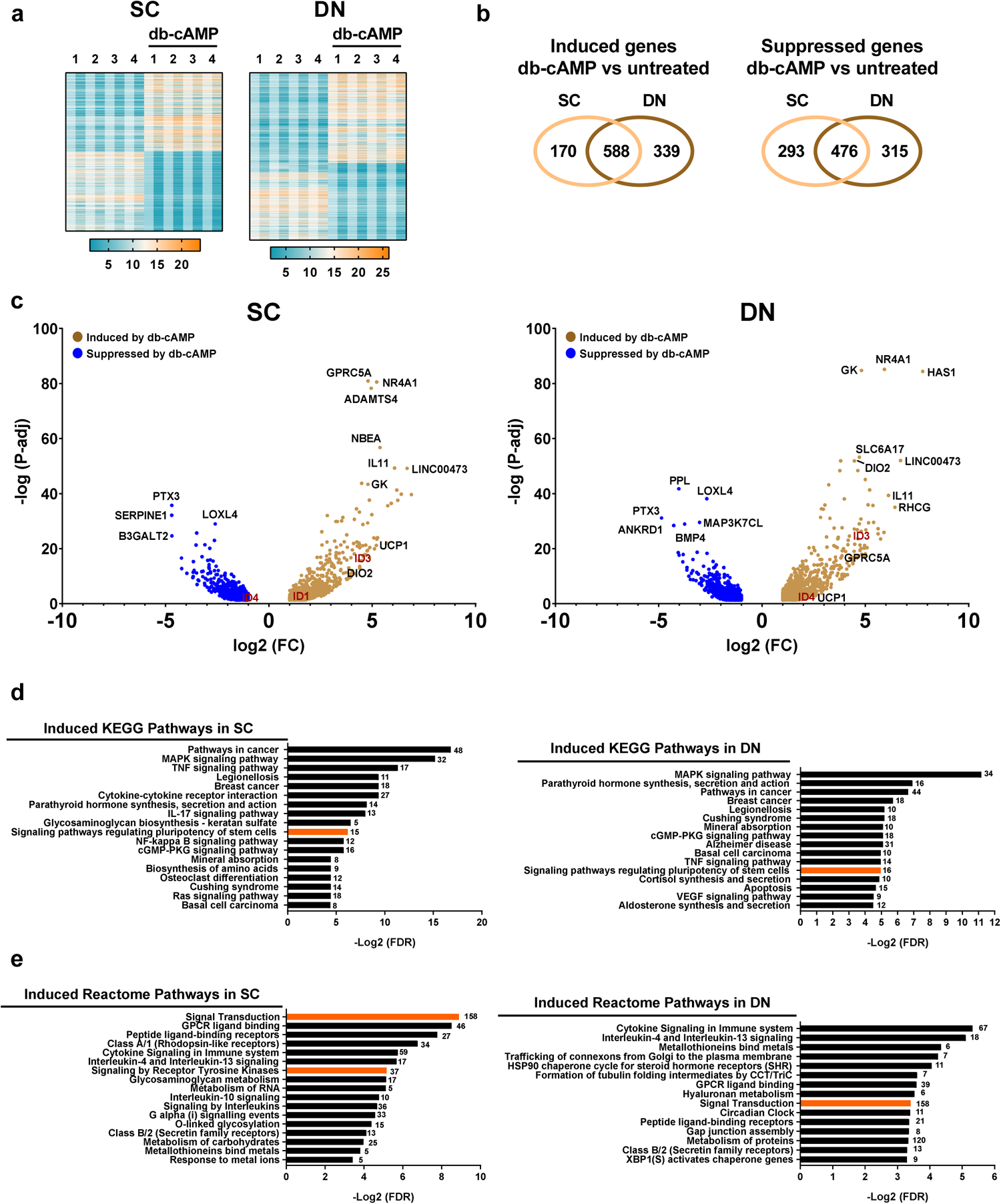
Differentially expressed genes (DEGs) and enriched pathways in subcutaneous (SC) and deep neck (DN)-derived adipocytes after 10 h of thermogenic activation driven by dibutyryl-cAMP (db-cAMP). (a) Heatmap displaying the expression values of DEGs in SC (left panel) or DN (right panel)-derived adipocytes in response to db-cAMP-driven stimulation. VST scores from DESeq2 analysis were used to generate the heatmaps. (b) Venn diagrams displaying the numbers of more (left panel) or less expressed (right panel) genes in comparison of db-cAMP vs untreated adipocytes of SC or DN origins. (c) Volcano plot illustrating the induced and suppressed genes by db-cAMP in SC (left panel) or DN (right panel)-derived adipocytes. (d) Overrepresented KEGG pathways which are upregulated in db-cAMP-treated SC (left panel) or DN (right panel)-derived adipocytes. (e) Overrepresented Reactome pathways which are upregulated in db-cAMP-treated SC (left panel) or DN (right panel)-derived adipocytes. Orange bars indicate the pathways in which genes encoding IDs are involved. Numbers indicate the number of DEGs in each pathway. FC: fold change, P-adj: adjusted p-value, FDR: false discovery rate.

To investigate the gene expression pathways affected by adrenergic-driven browning and thermogenic activation among the DEGs, we performed enrichment pathway analysis by KEGG and Reactome Pathway Database. We found that genes, whose levels were increased in db-cAMP-treated SC and DN-derived adipocytes, were significantly overrepresented in several KEGG pathways, such as the MAPK, TNF, and cGMP-PKG signaling, parathyroid hormone (PTH) synthesis, secretion, and action, and signaling pathways regulating pluripotency of stem cells (Figure 2d). TGF-β signaling was the only overrepresented pathway which was inhibited by db-cAMP in SC-derived adipocytes. MicroRNAs in cancer, axon guidance, Rap1 signaling, breast cancer, NOD-like receptor signaling, and AGE-RAGE signaling pathway in diabetic complication were downregulated in db-cAMP-treated DN-derived adipocytes. Our Reactome analysis showed that genes induced by db-cAMP in both adipocyte types were overrepresented in pathways of signal transduction, GPCR ligand binding, peptide ligand-binding receptors, cytokine signaling in immune system, IL4 and IL13 signaling, metallothionein bind metals, and class B/2 secretin family receptors (Figure 2e). We did not find any Reactome pathway when we reviewed genes suppressed by db-cAMP.

### Elevated level of ID1 and ID3 in adrenergic stimulated adipocytes

As we observed the upregulation of *ID1* and *ID3* by adrenergic stimulation in cervical area-derived adipocytes (Figure 2c and 3a, Supplementary Table 3 and 5), we were tempted to further investigate the importance of ID family in adipocyte browning and thermogenesis. These gene regulatory proteins were involved in signaling pathway regulating pluripotency of stem cells (Figure 2d, orange bar), signal transduction, and signaling by receptor tyrosine kinases (Figure 2e, orange bar), which were overrepresented during adrenergic stimulation. We could validate the elevation of ID1 and ID3 expressions by cAMP analogue-driven stimulation both at mRNA and protein level in both types of adipocytes (Figure 3b, c) registering particularly high induction of ID3. The expression of ID2 was not responsive to db-cAMP (Figure 3a-c) in either SC or DN-derived adipocytes. Although *ID4* was among the DEGs, we could not validate its upregulation by RT-qPCR and immunoblotting (Figure 3a-c). These results suggested that db-cAMP-stimulated upregulation of ID1 and especially ID3 may play a role in the induction of browning and thermogenic activation by adrenergic stimulation of human cervical-area derived adipocytes.

**Figure 3.**
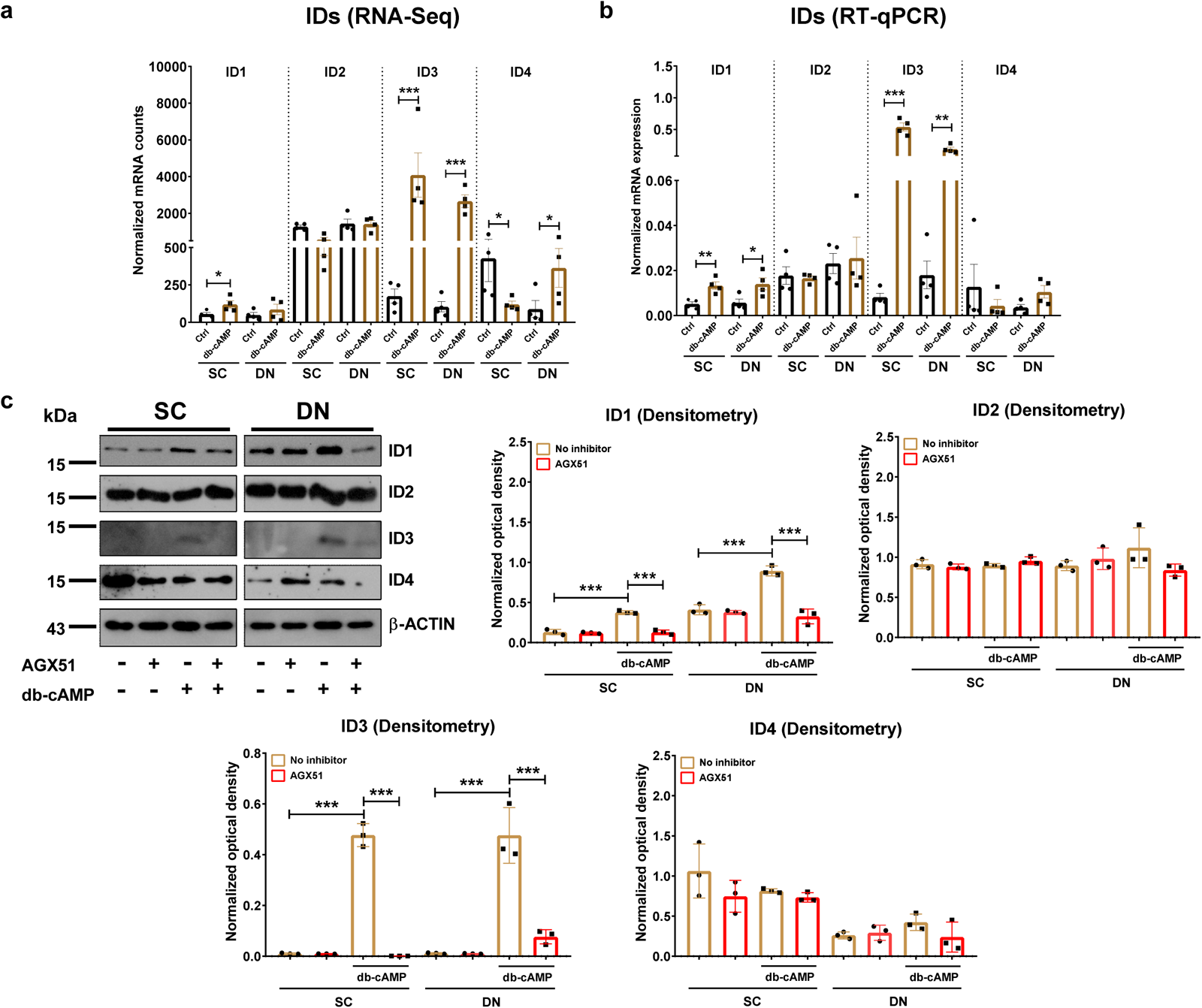
Expression levels of Inhibitor of DNA Binding (ID) 1-4 in subcutaneous (SC) and deep neck (DN)-derived adipocytes in the presence or absence of dibutyryl-cAMP (db-cAMP)-driven thermogenic activation. (a, b) The normalized mRNA expression based on RNA-seq analysis (a) or RT-qPCR (b). n=4, RNA-seq data was analyzed by DESeq2. Statistical analysis was performed by unpaired t-test. (c) The effect of the Inhibitor of DNA Binding (ID) antagonist, AGX51 on the expression of ID proteins detected by immunoblotting, n=3. The original pictures of the full-length blots are displayed in Supplementary Fig. 3. Statistical analysis was performed by one-way ANOVA. *p<0.05, **p<0.01, and ***p<0.001.

### ID antagonist abrogated cAMP analogue-stimulated elevation of oxygen consumption, extracellular acidification, and expression of mitochondrial complexes in adipocytes

Having observed the upregulation of IDs during adrenergic stimulation of human cervical area-derived adipocytes, we aimed to know whether inhibition of IDs would affect thermogenesis. We treated adipocytes in the presence or absence of thermogenic stimulation with the ID antagonist AGX51 which disrupts the interaction between ID and E proteins, resulting in unbound ID proteins that are rapidly degraded [18]. First, we investigated how AGX51 administration affected the expression of ID family members in human cervical area-derived adipocytes during db-cAMP-driven thermogenic activation. We found that in the presence of AGX51, the upregulation of ID1 and ID3 was completely prevented in both SC and DN-derived adipocytes (Figure 3c) leading to low level of ID1 and particularly of ID3. AGX51 treatment did not significantly affect the expression of ID2 and ID4 irrespective to the presence of thermogenic activation (Figure 3c).

Next, we measured cellular respiration of SC and DN-derived adipocytes to investigate the effect of AGX51 on oxygen consumption rate (OCR) and glycolysis related extracellular acidification rate (ECAR) during adrenergic stimulation. We found that db-cAMP increased maximal OCR and proton leak respiration that indirectly reflects heat generation, and ECAR in both types of adipocytes (Figure 4a, b). AGX51 hampered the db-cAMP-stimulated elevation of maximal and proton leak respiration and of ECAR in both SC and DN-derived adipocytes without affecting the OCR in unstimulated cells (Figure 4a, b). We also analyzed the effect of AGX51 on the expression of mitochondrial complex subunits and found that cAMP analogue-stimulated elevation of the subunit of mitochondrial complex I, which catalyzes the transfer of electrons from NADH to ubiquinone [27], was abrogated by AGX51 in both SC and DN-derived adipocytes (Figure 4c). ID antagonist hampered the db-cAMP-stimulated upregulation of complex II and IV only in SC-derived adipocytes (Figure 4c). AGX51 did not affect the expression of mitochondrial complex subunits III and V in either unstimulated or stimulated adipocytes (Figure 4c).

**Figure 4.**
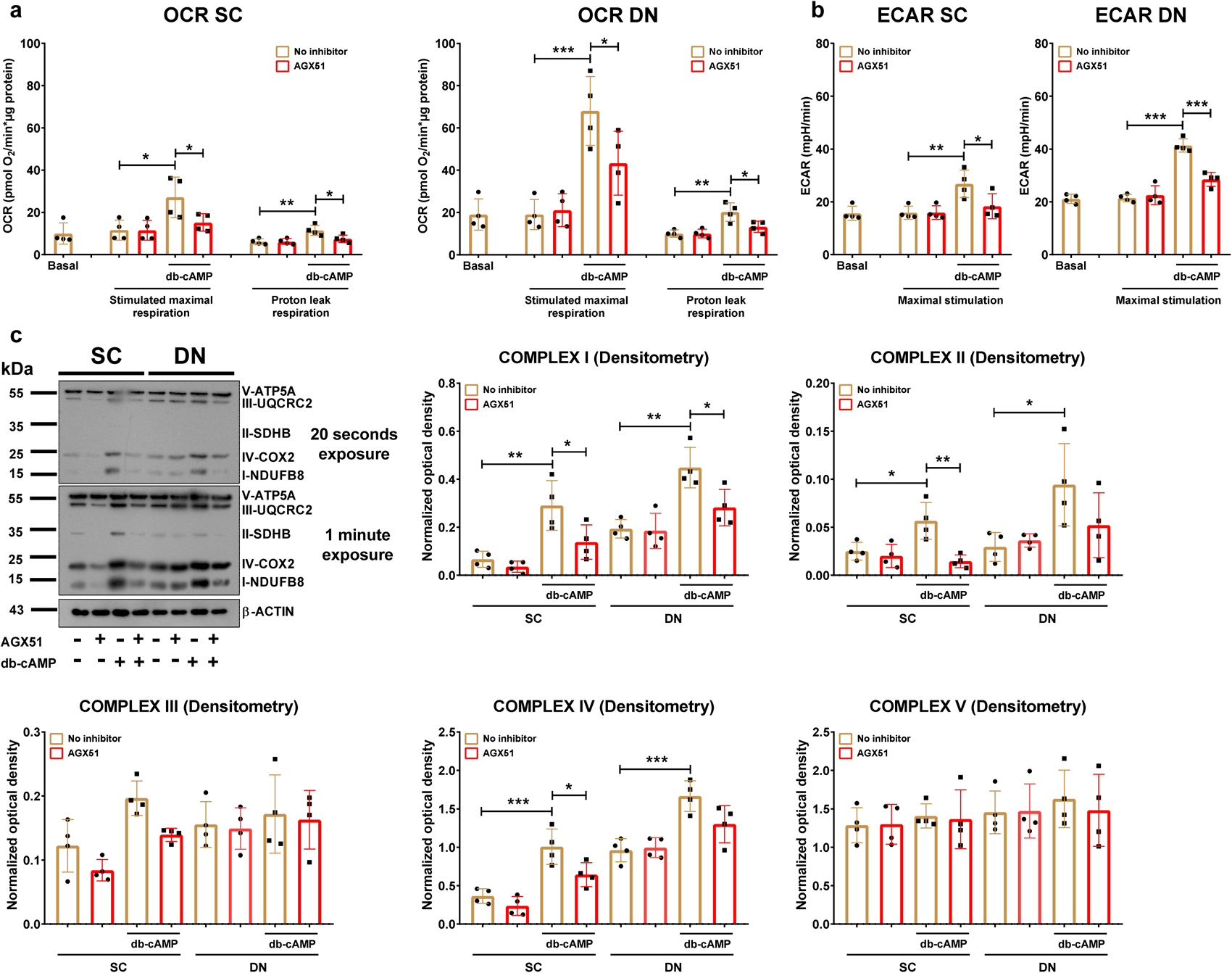
The effect of the Inhibitor of DNA Binding (ID) antagonist, AGX51 on the oxygen consumption and extracellular acidification rates (OCR and ECAR) and expression of mitochondrial complex subunits in subcutaneous (SC) and deep neck (DN)-derived adipocytes after 10 h of dibutyryl-cAMP (db-cAMP)-driven thermogenic activation. (a) Basal, db-cAMP stimulated maximal, and proton leak OCR and (b) basal and stimulated maximal ECAR were quantified by Seahorse extracellular flux analysis. (c) Protein expression of mitochondrial complex subunits detected by immunoblotting, n=4. The original pictures of the full-length blots are displayed in Supplementary Fig. 4. Statistical analysis was performed by one-way ANOVA followed by Tukey’s post-hoc test, *p<0.05, **p<0.01, and ***p<0.001.

### ID inhibition abrogated cAMP analogue-stimulated upregulation of thermogenic genes

Since we observed that proton leak respiration, which is associated with heat generation, was decreased by AGX51 during adrenergic stimulation, we intended to investigate the effect of AGX51 on the expression level of thermogenic genes. We found that the mRNA expression of *UCP1*, *PGC1a*, *DIO2*, and *CITED1* induced by db-cAMP was hindered by AGX51 in both SC and DN-derived adipocytes (Figure 5a). We also found that AGX51 abrogated the cAMP analogue-stimulated elevation of UCP1, PGC1a, and DIO2 expression at protein level in both types of adipocytes (Figure 5b). The ID inhibitor did not influence the expression of the investigated markers in the absence of thermogenic activation. These results are in accordance with our proton leak respiration data suggesting an important role of ID1 and ID3 in the efficient thermogenic response during adrenergic stimulation of human cervical area-derived adipocytes.

**Figure 5.**
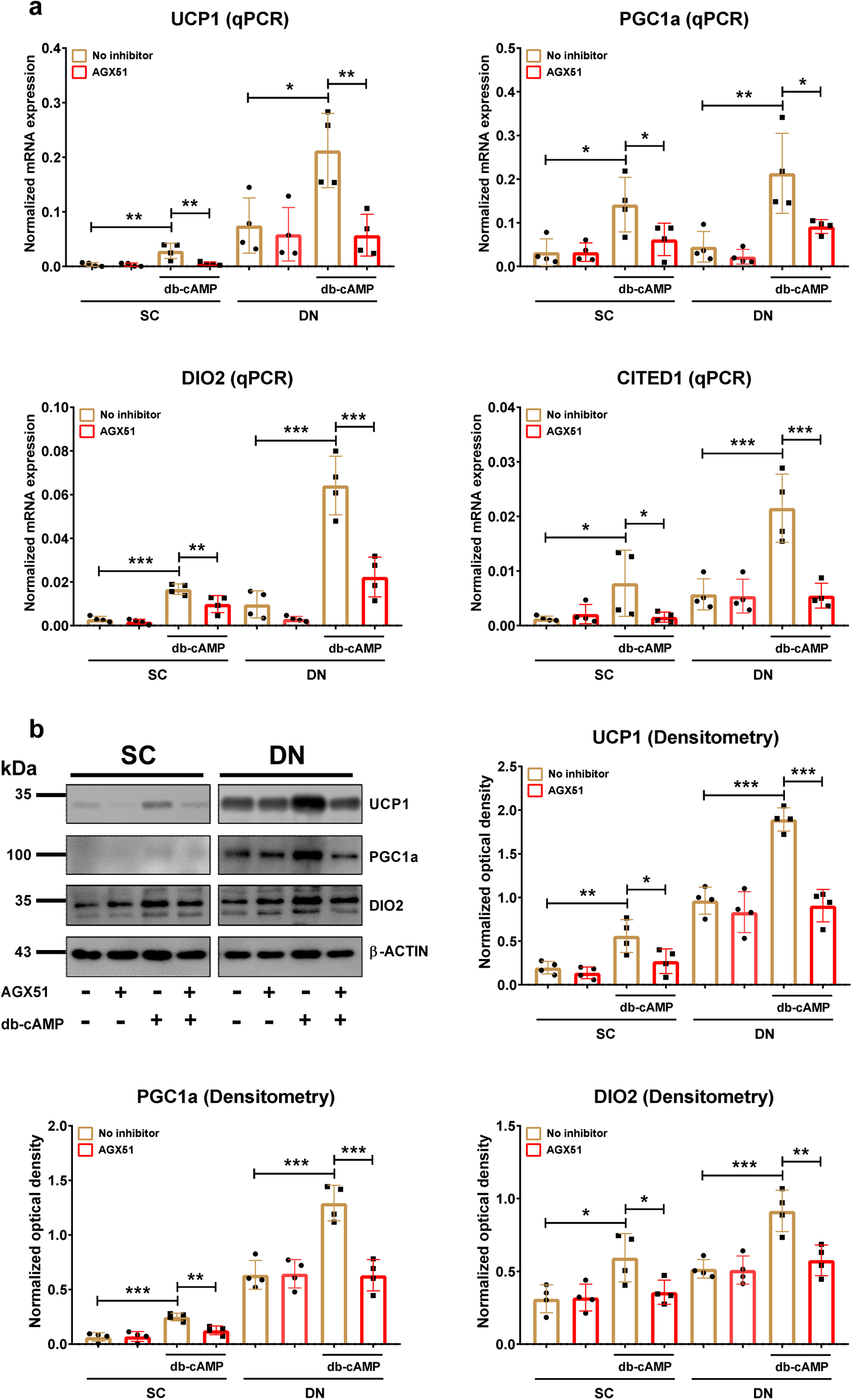
The effect of the Inhibitor of DNA Binding (ID) antagonist, AGX51 on the expression of thermogenic markers in subcutaneous (SC) and deep neck (DN)-derived adipocytes after 10 h of dibutyryl-cAMP (db-cAMP)-driven thermogenic activation. (a) The mRNA expression of *UCP1*, *PGC1a*, *DIO2*, and *CITED1* detected by RT-qPCR. (b) Protein expression of UCP1, PGC1a, and DIO2 detected by immunoblotting, n=4. The original pictures of the full-length blots are displayed in Supplementary Fig. 5. Statistical analysis was performed by one-way ANOVA followed by Tukey’s post-hoc test, *p<0.05, **p<0.01, and ***p<0.001.

## Discussion

The increased prevalence of obesity worldwide has prompted the scientific community to discover novel therapeutic strategies against this threat. Targeting brown and beige adipocytes has been a promising approach to augment energy expenditure due to their capability in dissipating energy as heat primarily via UCP1 activity [28]. The cervical area of adult humans is enriched in thermogenic adipocytes of both types marked by *UCP1, PGC1a, CITED1, PRDM16, LHX8, ZIC1,* and *TBX1* [23, 29]. To our knowledge, this is the first study in which the transcriptional response of SC and DN-derived human adipocytes to thermogenic activation was compared. Our results showed that adrenergic stimulation elevated both browning (especially in SC) and thermogenic capacity in human cervical area-derived adipocytes. Our KEGG pathway analysis showed that the genes whose expressions were induced by db-cAMP in both types of adipocytes were involved in several pathways, such as MAPK, cGMP-PKG, and TNF signaling. It was shown that elevated *UCP1* transcription is dependent on β3-adrenergic receptor-PKA-p38 MAPK pathway which results in the phosphorylation of activating transcription factor 2 (ATF2) and PGC1a [30–32]. Both ATF2 and PGC1a bind to the −2.5 kb enhancer region of the *UCP1* gene thus increasing its expression [33]. cGMP, which is produced by guanylyl cyclase, activates GMP-dependent protein kinase G (PKG) and plays a crucial role in thermogenesis [34]. TNF signaling pathway is involved in the activation of MAPK signaling which is important for non-shivering thermogenesis [35]. Another pathway commonly overrepresented in both SC and DN-derived adipocytes upon adrenergic stimulation by db-cAMP was PTH synthesis, secretion, and action, which promoted browning program in *ex vivo* differentiated human abdominal SC adipocytes [36]. Kovaničová *et al.* [37] reported that circulating PTH showed a positive correlation with the expression of thermogenic genes in human BAT. In addition, PTH level of ice-water swimmers was increased after 15-minute exposure. The Reactome pathway analysis showed that several pathways were overrepresented by genes which were induced by db-cAMP. Among the aforementioned pathways, signal transduction and GPCR ligand binding showed the highest significance. GPCRs play a crucial role in the regulation of thermogenic adipocytes [38]. We also found that the mRNA expression and secretion of IL6 and VEGFA was elevated by db-cAMP. Our previous study found that IL6 secreted by human beige adipocytes enhance browning in an autocrine manner [24]. VEGFA released by brown/beige adipocytes promoted BAT vascularization [39, 40]. Our data indicate orchestrated signaling events through parallel pathways for upregulation of thermogenic genes and batokines secretion to enhance thermogenic activity during db-cAMP treatment in human cervical area-derived adipocytes.

Pathway enrichment analysis by KEGG showed that IL-17 signaling pathway, which was shown to promote adaptive thermogenesis in murine inguinal WAT (iWAT) [41], was overrepresented only in SC-derived adipocytes during adrenergic stimulation. In addition, Reactome analysis showed that carbohydrate metabolism was overrepresented in SC-derived, whereas amino acids metabolism was overrepresented in DN-derived adipocytes suggesting that there are distinct properties between the two cell types in their response to adrenergic stimulation. *Ex vivo* differentiated human adipocytes have been used to reveal important regulatory elements of adipocyte browning and thermogenesis at molecular level [42–45], avoiding the invasive methods in *in vivo* systems. In line with the previous findings, we revealed a distinct regulatory elements between SC and DN-derived adipocytes during adrenergic stimulation, which may be targeted to alleviate obesity and related diseases in a cell type specific manner.

ID family proteins are abundantly expressed in preadipocytes, however, they are undetectable in differentiated adipocytes [46–48]. We previously found that the mRNA expression of ID1 and ID3 were declined in differentiated adipocytes derived from both SC and DN depots as compared to preadipocytes [23]. Our RNA-sequencing data showed that the expressions of *ID1*, *ID3*, and *ID4* were induced by adrenergic stimulation in human cervical area-derived adipocytes suggesting they have regulatory function in browning and thermogenesis. Pathway analysis revealed that the genes encoding ID family members are involved in ‘signaling pathways regulating pluripotency of stem cells’ in which cAMP analogue-driven upregulated genes were enriched in both types of adipocytes. We could confirm the moderate elevation of *ID1* and the strong upregulation of *ID3* by db-cAMP by RT-qPCR and western blot. Moldes *et al.* [49] reported that isoproterenol and forskolin increased the protein expression of Id2 and Id3 in 3T3-L1 cell lines and adipocytes isolated from rats. Our findings indicate different translational regulation of ID2 in human and murine adipocytes since ID2 expression was not affected by db-cAMP. Shinoda *et al.* [42] reported that *ID1-4* expression was higher in clonally-derived adult human brown as compared to white adipocytes. Previously, we also found that bone morphogenic protein 7 (BMP7), which drives brown adipocyte differentiation, elevated the expression of ID1 in both SC and DN-derived adipocytes [50].

The exact role and contribution of IDs, especially ID3, in energy expenditure, browning, and thermogenic activation in human adipocytes, remains to be clarified. Our presented data highlight the importance of ID1 and ID3 for the efficient thermogenic response during adrenergic stimulation in human SC and DN-derived adipocytes. We observed that the ID antagonist, AGX51 treatment led to the decreased protein expression of ID1 and ID3 at db-cAMP-stimulated conditions. In parallel to decreased IDs expression, the cAMP analogue-stimulated elevation of proton leak respiration that reflects heat generation, mitochondrial complex I, and thermogenic gene expression were abrogated by AGX51 pointing to possibly specific regulatory function of the ID proteins in thermogenesis. Research focused on the role of ID1 in adipose tissue, however, studies on ID3 are still lacking. Patil *et al.* [51] reported that murine BAT showed the highest expression of Id1 as compared to WAT and cold strongly induced the Id1 protein expression in BAT. However, in contrast with our findings, they found that iWAT from whole body *Id1^-/-^* mice expressed higher levels of Ucp1 and Dio2 when mice were exposed to cold. They speculated that cold-induced elevation of Id1 may prevent excessive thermogenesis by acting as a thermogenic suppressor [51]. Another group also reported contrasting results with our findings [46]. They performed protein expression analysis from different tissues of adult mice and found that Id1 showed high expression in both WAT and BAT. They revealed that iWAT of *Id1^-/-^*mice had higher energy expenditure along with increased lipolysis and fatty acid oxidation as compared to *Id1^+/+^* mice [46, 51]. These observations suggest that Id1 may play a role in adipogenesis, iWAT metabolism, and BAT-mediated thermogenesis. Zhao *et al*. [52] reported that WAT from whole body Id1-deficient mice had higher expression level of PGC1a.

Id3 regulates adiponectin expression by binding to E47, which potentiates SREBP-1c-mediated adiponectin promoter activation [53]. We searched for interactions of ID3 with other DEGs and found that ID3 interacts with the transcription factor interferon regulatory factor 4 (IRF4) which was upregulated by db-cAMP in both adipocyte types. Kong *et al*. [54] reported that the mRNA and protein expression of IRF4 was induced by cold and cAMP in mouse iBAT, iWAT, and eWAT as well. Furthermore, the physical and functional interaction between IRF4 and PGC1a promoted thermogenesis in mice. ID3 also interacts with other transcription factors, such as BACH2 and SOX4, whose expression was induced by db-cAMP only in DN. In addition, we also found that ID3 interacts with BMP4, which was downregulated by db-cAMP in both SC and DN-derived adipocytes. This secreted factor promotes the whitening of murine BAT and blunts the activity of mature brown adipocytes [55]. ID1 and ID3 have overlapping regulatory roles including angiogenesis [56]. As BAT is highly vascularized, elevated vascular density and angiogenesis are crucial prerequisites for the expansion during WAT browning and thermogenic activation in BAT [57].

The discrepancy between our and other groups’ findings may be explained by different regulatory mechanisms between humans and mice. It is also important to underline that in our experiments, the ID antagonist was only administered to fully differentiated human adipocytes during adrenergic stimulation and therefore, the emerging effect may be different compared to the absence of Id1 from the beginning of adipose tissue development in mice. The advantage of the application of a pharmacological inhibitor was that its effect could be restricted only to the activation period four which the current study focused on. However, we are aware that we could not rule out the possible off-target effects of AGX51 on adipocytes. Therefore, further studies are required for better understanding of transcriptional regulatory functions of ID1 and ID3 in thermogenesis, especially in human adipocytes.

## Materials and methods

### Materials

All chemicals were from Sigma-Aldrich (Munich, Germany) unless stated otherwise.

### Ethics statements and obtained tissue samples

Tissue collection was approved by the Medical Research Council of Hungary (20571-2/2017/EKU) followed by the EU Member States’ Directive 2004/23/EC on presumed consent practice for tissue collection. All experiments were carried out in accordance with the guidelines of the Helsinki Declaration. Written informed consent was obtained from all participants before the surgical procedure. During thyroid surgeries, a pair of DN and SC adipose tissue samples was obtained to rule out inter-individual variations. Patients with known diabetes, malignant tumor, or with abnormal thyroid hormone levels at the time of surgery were excluded. Human adipose-derived stromal cells (hASCs) were isolated from SC and DN fat biopsies as described previously [23, 14]. The *FTO* 1421085 genotype for all involved donors were heterozygous (T/C).

### Differentiation and treatment of hASCs

Primary cervical adipocytes were differentiated from stromal-vascular fractions, isolated from adipose tissue biopsies, containing hASCs according to a described protocol applying insulin, cortisol, T3, dexamethasone, and short-term rosiglitazone treatment [58]. After 14 days of differentiation, adipocytes were treated with a single bolus of 500 µM db-cAMP (D0260) for 10 hours to mimic *in vivo* cold-induced thermogenesis [59], 10 µM AGX51 (Med Chem, NJ, USA, HY-129241) [18], or combination of db-cAMP and AGX51. Adipocytes incubated in Dulbecco’s Modified Eagle Medium/Nutrient Mixture F-12 (DMEM-F12) medium, supplemented with 33 µM biotin, 17 µM pantothenic acid, and 100 U/ml penicillin/streptomycin, was used as control.

### RNA isolation and RNA-sequencing analysis

Cells were collected and total RNA was isolated as described previously [14]. The concentration and purity of the isolated RNA was checked by using Nanodrop 2000 Spectrophotometer (Thermo Fisher, Waltham, MA, USA). RNA-sequencing analysis was carried out as described in our previous study [23]. FASTQ file data were analyzed by Galaxy (https://usegalaxy.org/) [60]. Significant DEGs were defined based on adjusted p values p <0.05 and log_2_ fold change threshold >1. Heatmap was generated by GraphPad 8.0 (GraphPad Software, San Diego, CA, USA) using VST score. Enrichment pathway analysis for KEGG and Reactome was performed by STRING (https://string-db.org/) [61]. Prediction of browning capacity was analyzed using publicly available webtool ProFAT (http://ido.helmholtz-muenchen.de/profat/) [21] and BATLAS (https://fgcz-shiny.uzh.ch/tnb_ethz_BATLAS_app/) [20].

### RT-qPCR

RNA was diluted to 100 ng/µL for all samples and was reverse transcribed to cDNA by using reverse transcription kit (Thermo Fisher Scientific, 4368814) following the manufacturer’s instructions. Validated TaqMan assays used in qPCR were designed and supplied by Thermo Fisher Scientific as listed in Supplementary Table 1. qPCR was performed in Light Cycler® 480 II (Roche). The following conditions were set to perform the reactions: initial denaturation step at 95 °C for 1 min followed by 50 cycles of 95 °C for 12 sec and 60 °C for 30 sec. Gene expression values were calculated by the comparative threshold cycle (Ct) method as described in the previous publication [62]. Gene expressions were normalized to the geometric mean of *ACTB* and *GAPDH*. Normalized gene expression levels equal 2^-ΔCt^.

### Measurement of cytokine release

Conditioned medium was collected after the abovementioned treatments and stored in - 70°C until measurement. The concentration of IL-6 (DY206) and VEGFA (DY293B-05) in the conditioned media was measured by using ELISA DuoSet Kit (R&D Systems, Minneapolis, MN, USA). The concentration was calculated by following the manufacturer’s instructions.

### Immunoblotting and densitometry

Immunoblotting and densitometry were carried out as described previously [14, 63]. Antibodies and working dilutions are listed in Supplementary Table 2. The expression of the visualized immunoreactive proteins were quantified by densitometry using the FIJI ImageJ software (National Institutes of Health, Bethesda, MD, USA) as previously described [63].

### Extracellular flux assay

OCR and ECAR of adipocytes were measured using an XF96 oxymeter (Seahorse Biosciences, North Billerica, MA, USA) as described previously [14, 63]. After recording the baseline OCR, 500 µM db-cAMP, 10 µM AGX51, or combination of the cAMP analogue and the inhibitor were injected to the cells. Then, stimulated OCR was recorded every 30 min for 10 hours. Proton leak respiration was determined after injecting ATP synthase blocker (oligomycin) at 2 μM concentration. Cells received a single bolus of Antimycin A at 10 μM concentration for baseline correction (measuring non-mitochondrial respiration). The OCR was normalized to protein content.

### Statistical analysis

The results are expressed as mean±SD. Normality of distribution of the data was tested by Shapiro–Wilk test. Multiple comparison among groups were analyzed by one-way ANOVA followed by Tukey’s post-hoc test. The data were visualized and analyzed by using GraphPad Prism 8.

## Abbreviations

BAT: brown adipose tissue
Db-cAMP: dibutyryl-cyclic adenosine monophosphate
DEGs: differentially expressed genes
DN: deep neck
ECAR: extracellular acidification rate
hASCs: human adipose-derived stromal cells
ID: inhibitor of DNA binding
iWAT: inguinal white adipose tissue
OCR: oxygen consumption rate
PET-CT: positron emission tomography-computed tomography
SC: subcutaneous
UCP1: uncoupling protein 1
VAT: visceral white adipose tissue
WAT: white adipose tissue

## Data availability

The RNA-sequencing datasets generated and analyzed for this study can be found in the Sequence Read Archive (SRA) database [https://www.ncbi.nlm.nih.gov/sra] under accession number PRJNA1093362.

## Author contributions

R.Ar. — conceptualization, methodology, investigation, data curation, validation, visualization, writing - original draft; B. Á. V. — methodology, investigation; R. Al. — methodology; G. K. — methodology; Y. Q. A. — methodology; S. P. — methodology, investigation; F. G. — methodology, resources; L. F. — conceptualization, funding acquisition, supervision, writing – review and editing; E. K. — conceptualization, project administration, funding acquisition, supervision, writing – review and editing.

## Acknowledgements

We thank Dr. Éva Csősz for her exceptional help in reviewing the manuscript before its submission and Jennifer Nagy for technical assistance. This research was funded by the National Research, Development and Innovation Office (NKFIH-FK145866 and PD146202) of Hungary. BÁV was supported by the ÚNKP-23-3-II-DE-156 New National Excellence Program of the Ministry for Innovation and Technology from the source of the National Research, Development and Innovation Fund. SP was supported by the project TKP2021-NKTA-34, which has been implemented with the support provided by the Ministry of Culture and Innovation of Hungary from the National Research, Development and Innovation Fund, financed under the TKP2021-NKTA funding scheme.

