## Supplementary Figures and Tables for "Upregulation of Inhibitor of DNA Binding 1 and 3 is Important for Efficient Thermogenic Response in Human Adipocytes"

**SUPPLEMENTARY MATERIAL**

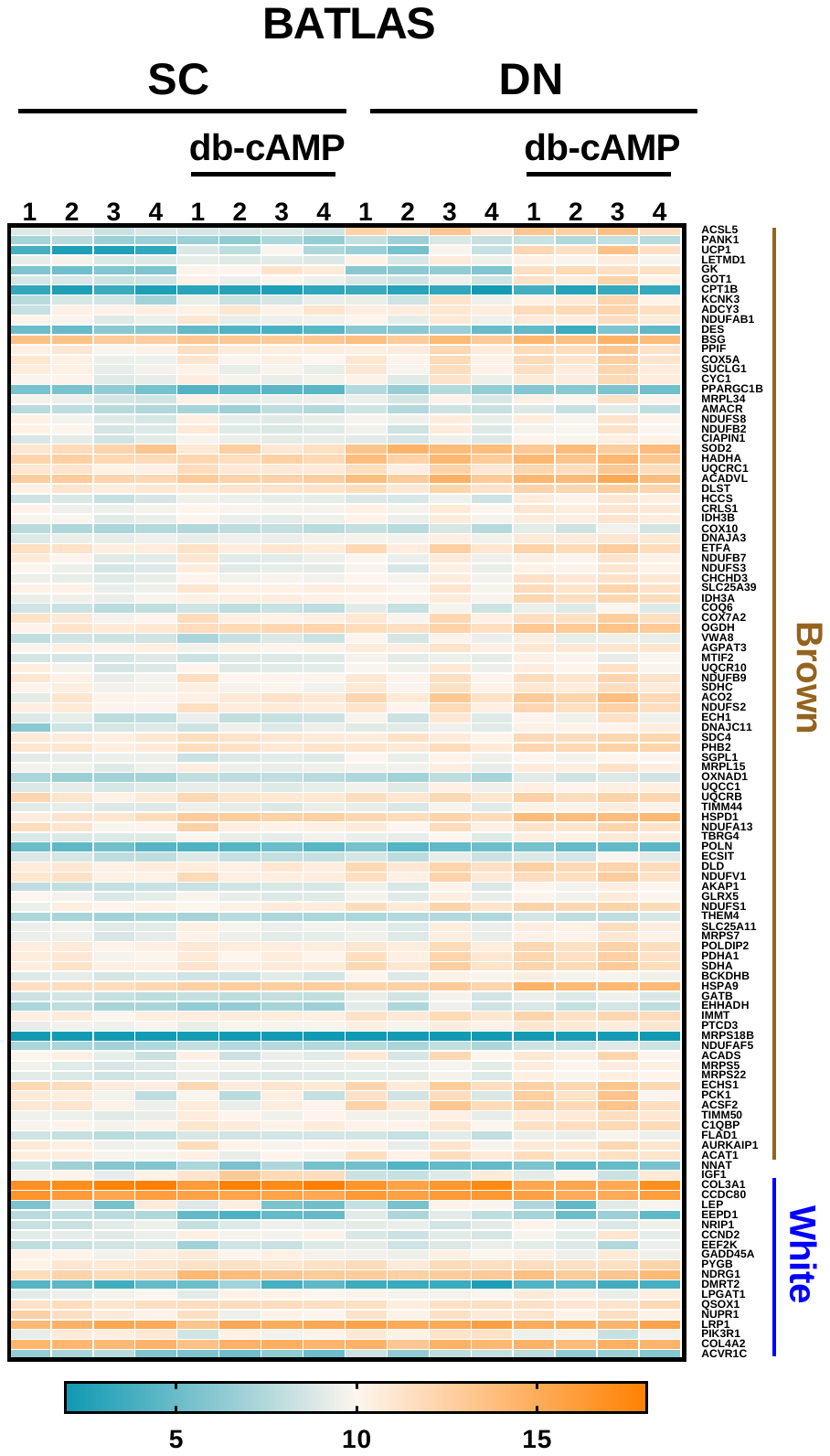

**Supplementary Figure 1.** Heatmap displaying the expression pattern of BATLAS markers, n=4 of each group. Adipocytes were differentiated and treated as in Figures 1-3. VST scores from DESeq2 analysis were used to generate the heatmap.

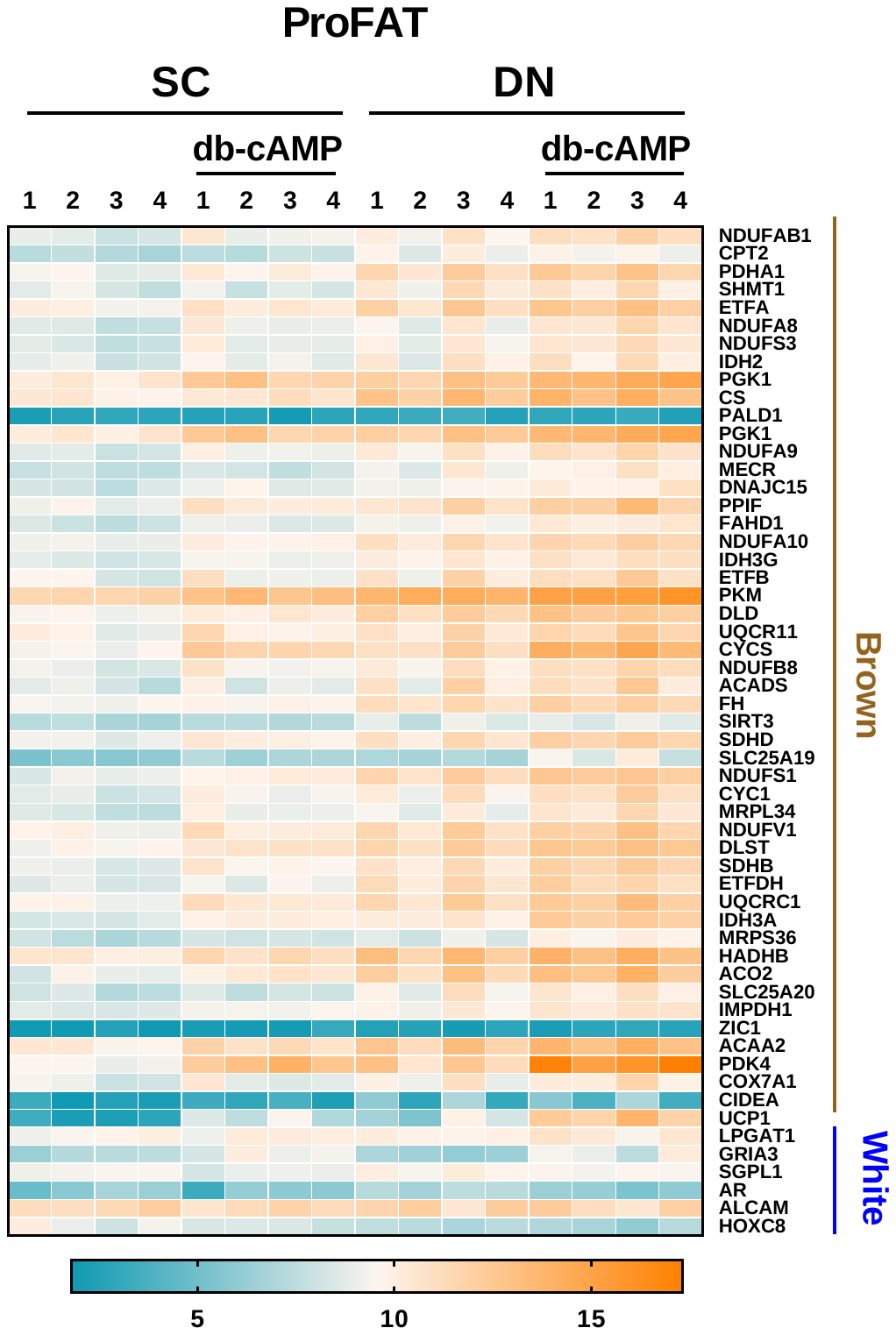

**Supplementary Figure 2.** Heatmap displaying the expression pattern of ProFAT markers, n=4 of each genotype. Adipocytes were differentiated and treated as in Figures 1-3. VST scores from DESeq2 analysis were used to generate the heatmap.

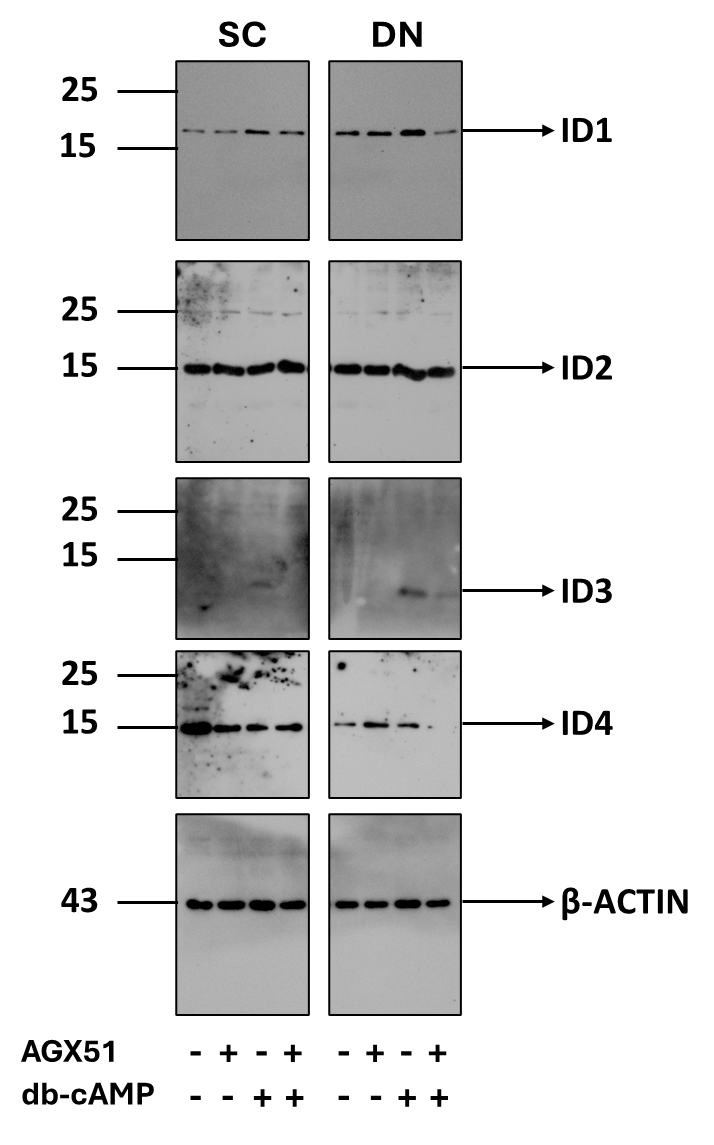

**Supplementary Figure 3. Uncropped images presented with molecular weight ladders for Figure 3c.** β-actin was used as endogenous control. Detailed information regarding the antibodies and working dilution are displayed in Supplementary Table 2.

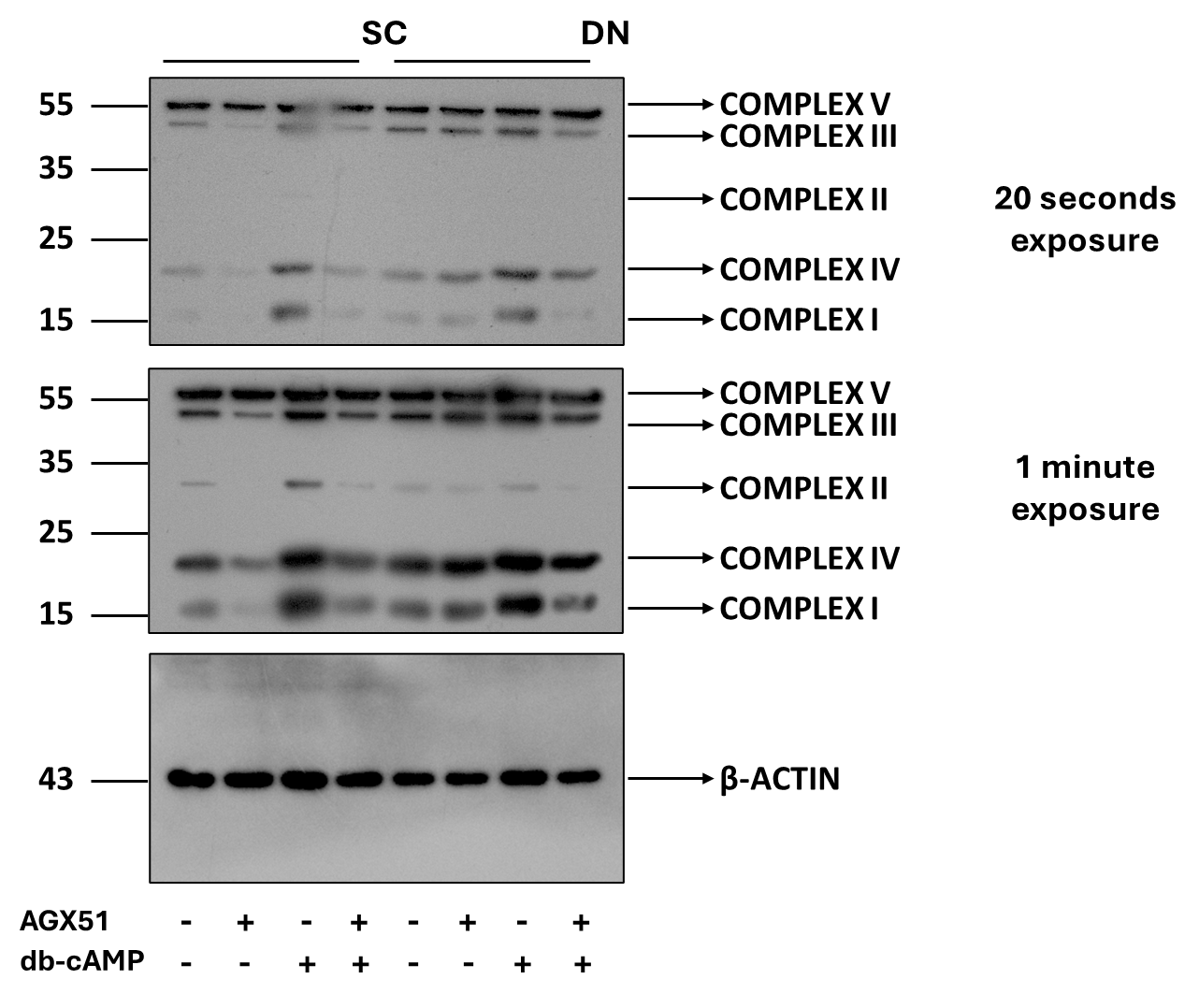

**Supplementary Figure 4. Uncropped images presented with molecular weight ladders for Figure 4c.** β-actin was used as endogenous control. Detailed information regarding the antibodies and working dilution are displayed in Supplementary Table 2.

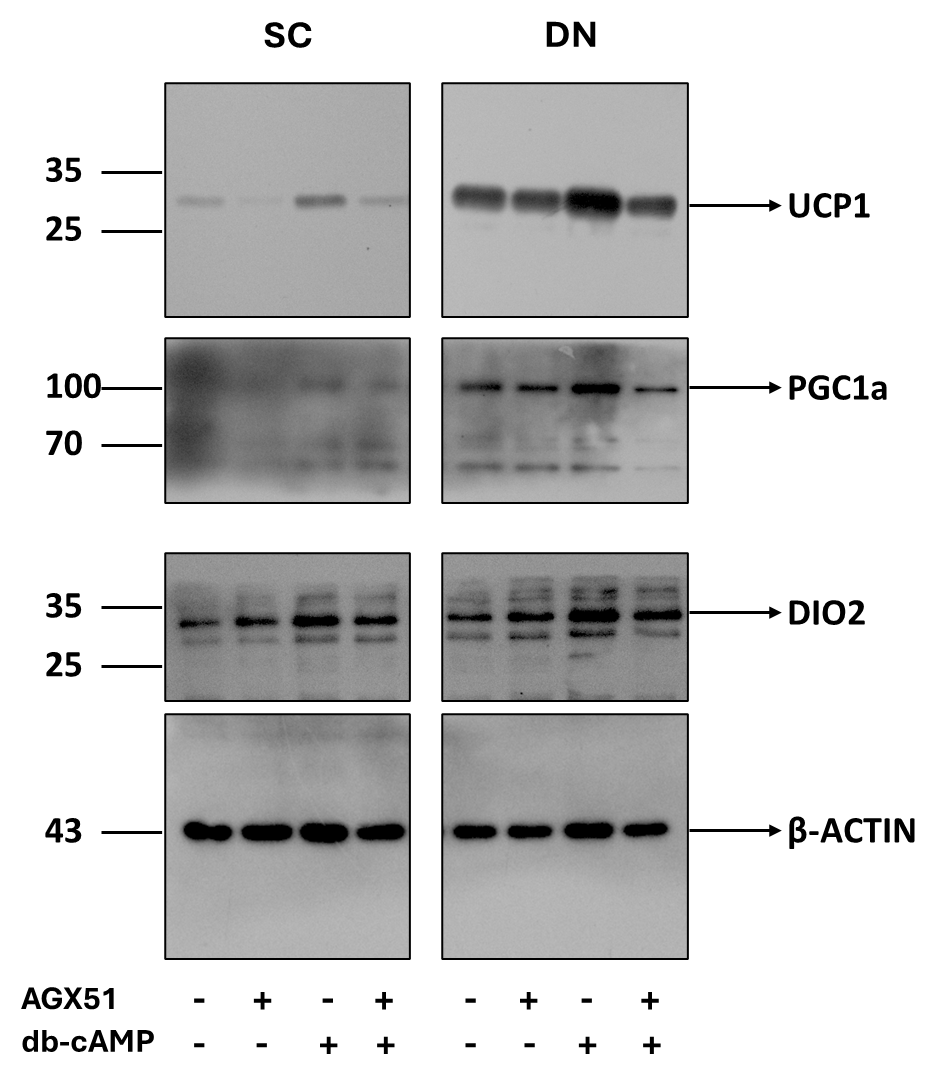

**Supplementary Figure 5. Uncropped images presented with molecular weight ladders for Figure 5b.** β-actin was used as endogenous control. Detailed information regarding the antibodies and working dilution are displayed in Supplementary Table 2.

**Supplementary Table 1.** Gene primers and probes

| **GENES** | **ASSAY ID** |
| --- | --- |
| *UCP1* | Hs00222453_m1 |
| *PGC1a* | Hs01075227_m1 |
| *DIO2* | Hs00399438_m1 |
| *CITED1* | Hs00918445_g1 |
| *ID1* | Hs00179727_m1 |
| *ID2* | Hs00176502_m1 |
| *ID3* | Hs00988962_m1 |
| *ID4* | Hs00160173_m1 |
| *GAPDH* | Hs99999905_m1 |

**Supplementary Table 2.** Antibodies used in immunoblotting

| Antibody | Company | Catalog Number | Dilution |
| --- | --- | --- | --- |
| UCP1 | R&D Systems, Minneapolis, MN, USA | MAB6158 | 1:750 |
| PGC1α | Novus Biologicals, Centennial, CO, USA | NBP1-04676 | 1:1000 |
| DIO2 | Invitrogen, USA | PA5-49631 | 1:2000 |
| ID1 | Novus Biologicals | NBP2-66897 | 1:1000 |
| ID2 | Novus Biologicals | NBP2-27194SS | 1:1000 |
| ID3 | Cell Signaling Technology, Beverly, MA, USA | D16D10 | 1:1000 |
| ID4 | Novus Biologicals | NBP3-10934 | 1:1000 |
| Total OXPHOS | Abcam, Cambridge, MA, USA | ab110411 | 1:1000 |
| Β-Actin | Sigma Aldrich | A2066 | 1:10000 |
| HRP-conjugated goat anti-rabbit IgG | Advansta, San Jose, CA, USA | R-05072-500 | 1:5000 |
| HRP-conjugated goat anti-mouse IgG | Advansta, San Jose, CA, USA | R-05071-500 | 1:5000 |

**Supplementary Table 3.** Upregulated genes by cAMP in SC adipocytes. Log_2_ fold change (FC) ≥1, Adjective p value <0.05

| **Symbol** | **Log2 Fold Change** | **Adj. P Value** |  | **Symbol** | **Log2 Fold Change** | **Adj. P Value** |
| --- | --- | --- | --- | --- | --- | --- |
| *SST* | 6.90 | 2.23E-40 |  | *PLCB1* | 1.82 | 6.95E-06 |
| *LINC00473* | 6.69 | 6.38E-50 |  | *QPCTL* | 1.82 | 8.93E-08 |
| *RHCG* | 6.41 | 1.59E-40 |  | *MESD* | 1.82 | 1.16E-08 |
| *EREG* | 6.24 | 2.80E-38 |  | *AMIGO2* | 1.82 | 3.21E-06 |
| *LYPD3* | 6.19 | 4.99E-42 |  | *SP6* | 1.81 | 0.004006773 |
| *IL11* | 6.08 | 5.20E-50 |  | *CASC15* | 1.81 | 0.000919263 |
| *OASL* | 5.92 | 3.58E-37 |  | *ZC3H12A* | 1.81 | 1.04E-08 |
| *PTPRN* | 5.75 | 2.61E-36 |  | *AVPI1* | 1.81 | 0.000210592 |
| *C11orf86* | 5.71 | 1.04E-22 |  | *BTG3* | 1.81 | 4.26E-05 |
| *NBEA* | 5.38 | 1.71E-57 |  | *KLF17* | 1.81 | 0.023118794 |
| *C2CD4A* | 5.26 | 1.06E-24 |  | *F2RL3* | 1.80 | 0.025651161 |
| *IL1RN* | 5.22 | 3.74E-24 |  | *NKAIN1* | 1.80 | 4.09E-05 |
| *NR4A1* | 5.22 | 2.65E-81 |  | *HYAL3* | 1.80 | 3.15E-06 |
| *UCP1* | 5.20 | 6.30E-22 |  | *CYP11A1* | 1.80 | 4.82E-08 |
| *SERTM1* | 5.11 | 1.12E-20 |  | *LBP* | 1.80 | 1.09E-06 |
| *NTRK1* | 5.08 | 1.21E-24 |  | *CST1* | 1.80 | 0.044335349 |
| *SH2D2A* | 4.98 | 4.46E-33 |  | *BLK* | 1.79 | 0.028956133 |
| *ADAMTS4* | 4.95 | 4.90E-79 |  | *FRMD3* | 1.79 | 0.01189379 |
| *FAM167A* | 4.88 | 3.89E-21 |  | *CHD1* | 1.79 | 0.000377565 |
| *GPR183* | 4.87 | 1.90E-17 |  | *TNFAIP6* | 1.79 | 0.001547207 |
| *NR4A2* | 4.86 | 2.41E-38 |  | *NGEF* | 1.78 | 0.008312217 |
| *GPRC5A* | 4.80 | 1.14E-81 |  | *ANKRD53* | 1.78 | 7.14E-05 |
| *GK* | 4.79 | 4.17E-44 |  | *CDC7* | 1.78 | 1.06E-05 |
| *FCER1G* | 4.78 | 3.93E-20 |  | *MB21D2* | 1.77 | 4.08E-07 |
| *TUBB2B* | 4.78 | 4.47E-30 |  | *SLC19A2* | 1.77 | 1.67E-06 |
| *TRH* | 4.75 | 6.75E-20 |  | *GPCPD1* | 1.76 | 7.83E-07 |
| *CCR7* | 4.70 | 1.82E-18 |  | *CYSLTR2* | 1.76 | 0.046043321 |
| *TAC1* | 4.69 | 5.36E-22 |  | *GMNN* | 1.76 | 6.31E-05 |
| *SEMA6B* | 4.64 | 1.52E-18 |  | *FAM83A* | 1.76 | 0.040382931 |
| *RASD2* | 4.56 | 6.65E-18 |  | *PAQR5* | 1.75 | 0.00016265 |
| *CRISPLD2* | 4.55 | 1.75E-29 |  | *LONRF3* | 1.75 | 0.000530915 |
| *SCG2* | 4.54 | 8.00E-22 |  | *PAPPA2* | 1.75 | 0.005544605 |
| *NPPC* | 4.51 | 2.35E-16 |  | *FMNL1* | 1.75 | 7.48E-06 |
| *CALCA* | 4.50 | 2.94E-16 |  | *PNP* | 1.74 | 6.49E-07 |
| *SMOX* | 4.49 | 1.91E-44 |  | *FOSL2* | 1.74 | 0.002220857 |
| *CYP2S1* | 4.40 | 4.59E-21 |  | *MSX1* | 1.74 | 5.48E-05 |
| *S100P* | 4.39 | 1.35E-13 |  | *TREM1* | 1.74 | 0.038421565 |
| *MUC13* | 4.39 | 2.89E-14 |  | *SAMD11* | 1.74 | 0.002644271 |
| *USP2* | 4.38 | 3.01E-22 |  | *MT1G* | 1.74 | 0.00553787 |
| *CTH* | 4.38 | 5.01E-34 |  | *ST3GAL5* | 1.74 | 7.95E-05 |
| *MT1A* | 4.37 | 1.77E-16 |  | *DAB2IP* | 1.73 | 4.42E-07 |
| *TUBA4A* | 4.36 | 8.85E-23 |  | *WDR86* | 1.73 | 0.042115157 |
| *SLC6A17* | 4.28 | 1.25E-21 |  | *CAMK2N2* | 1.73 | 0.000878317 |
| *KLHL13* | 4.26 | 3.78E-23 |  | *HEYL* | 1.72 | 9.88E-05 |
| *SIK1* | 4.24 | 3.71E-35 |  | *C5orf49* | 1.72 | 0.032265522 |
| *C2CD4B* | 4.18 | 1.26E-11 |  | *NAGS* | 1.72 | 0.041607702 |
| *PDE4D* | 4.14 | 2.31E-31 |  | *SHISA3* | 1.72 | 0.001552699 |
| *IRF4* | 4.13 | 8.15E-19 |  | *PLOD2* | 1.72 | 0.003933082 |
| *MLLT11* | 4.10 | 1.30E-22 |  | *LGR4* | 1.71 | 0.000255608 |
| *C2orf66* | 4.06 | 4.61E-12 |  | *BMP2* | 1.71 | 9.51E-05 |
| *BABAM2-AS1* | 4.04 | 2.93E-20 |  | *AP1S3* | 1.71 | 1.17E-05 |
| *OVOL1* | 4.04 | 1.05E-12 |  | *ACSL4* | 1.71 | 9.43E-05 |
| *AREG* | 4.03 | 1.75E-12 |  | *AADACP1* | 1.71 | 0.033909799 |
| *ID3* | 3.97 | 1.97E-17 |  | *ZBTB32* | 1.71 | 0.026461102 |
| *SDS* | 3.96 | 2.66E-10 |  | *TGFA* | 1.71 | 0.023213637 |
| *SDIM1* | 3.94 | 1.62E-11 |  | *SLC8A2* | 1.71 | 0.034669548 |
| *CCDC177* | 3.91 | 8.24E-11 |  | *OTOF* | 1.71 | 0.004865074 |
| *TMEM100* | 3.87 | 1.03E-13 |  | *SLC7A8* | 1.70 | 0.00012169 |
| *CACNA1G* | 3.84 | 7.08E-10 |  | *PCDH12* | 1.70 | 0.005524973 |
| *ATP2A3* | 3.80 | 2.99E-21 |  | *BHLHE40* | 1.70 | 1.80E-07 |
| *STC1* | 3.78 | 1.55E-09 |  | *SH3BP5* | 1.70 | 0.00747364 |
| *FLRT3* | 3.77 | 1.45E-16 |  | *SLC22A4* | 1.70 | 1.07E-05 |
| *PRR5L* | 3.77 | 1.00E-25 |  | *ZFP92* | 1.70 | 0.041822358 |
| *ECEL1* | 3.75 | 9.16E-10 |  | *B3GNT2* | 1.70 | 4.53E-07 |
| *CHRDL2* | 3.73 | 5.01E-12 |  | *SOCS3* | 1.70 | 1.65E-05 |
| *GAL* | 3.65 | 2.55E-17 |  | *PRPS2* | 1.69 | 0.000411662 |
| *KLHL15* | 3.64 | 3.20E-23 |  | *FZD8* | 1.69 | 3.53E-09 |
| *DIO2* | 3.61 | 8.21E-12 |  | *LINC01128* | 1.69 | 0.000160984 |
| *TFPI2* | 3.58 | 4.79E-32 |  | *ARHGAP32* | 1.69 | 8.56E-06 |
| *GPAT3* | 3.57 | 2.45E-19 |  | *SEMA3A* | 1.68 | 0.000290194 |
| *PITPNC1* | 3.52 | 8.36E-31 |  | *KCTD20* | 1.68 | 0.001893889 |
| *PHYHIP* | 3.48 | 3.92E-13 |  | *NOG* | 1.68 | 0.00017282 |
| *SIX2* | 3.48 | 4.70E-22 |  | *MKRN9P* | 1.68 | 0.024526582 |
| *MFSD2A* | 3.48 | 1.57E-15 |  | *ECE2* | 1.67 | 0.000352704 |
| *SHISA2* | 3.47 | 8.83E-13 |  | *ADAM28* | 1.67 | 0.031883798 |
| *RASL11B* | 3.45 | 1.59E-09 |  | *SNTG2* | 1.67 | 0.00738185 |
| *ACTBL2* | 3.43 | 6.00E-08 |  | *TFCP2L1* | 1.67 | 0.038858768 |
| *REN* | 3.42 | 9.09E-09 |  | *FZD4* | 1.67 | 4.22E-05 |
| *SLC16A6* | 3.41 | 4.44E-10 |  | *HERC4* | 1.66 | 2.55E-05 |
| *SH3TC1* | 3.40 | 4.94E-19 |  | *CH25H* | 1.66 | 0.000121473 |
| *PGAP1* | 3.39 | 8.04E-17 |  | *BMP6* | 1.66 | 0.000143253 |
| *PGAP1* | 3.39 | 5.71E-08 |  | *FAM131A* | 1.66 | 3.47E-05 |
| *P4HA3* | 3.36 | 1.14E-15 |  | *IQSEC3* | 1.65 | 0.007343356 |
| *CHGB* | 3.34 | 2.02E-08 |  | *CBARP* | 1.64 | 0.003241158 |
| *DLK1* | 3.32 | 1.25E-07 |  | *UCN* | 1.64 | 0.012936944 |
| *CHMP1B* | 3.31 | 1.35E-30 |  | *MRPS30-DT* | 1.64 | 0.018344991 |
| *WNT1* | 3.29 | 1.27E-07 |  | *SLC22A23* | 1.64 | 7.63E-05 |
| *TNFRSF18* | 3.28 | 4.67E-08 |  | *SPHK1* | 1.64 | 5.45E-06 |
| *DUSP4* | 3.27 | 2.99E-21 |  | *ENPEP* | 1.64 | 0.000239906 |
| *PPARGC1A* | 3.26 | 2.93E-19 |  | *ENOSF1* | 1.64 | 1.34E-06 |
| *LYVE1* | 3.26 | 9.02E-09 |  | *HOXA5* | 1.63 | 0.02541194 |
| *DNAH17* | 3.25 | 1.96E-17 |  | *GPR161* | 1.63 | 7.69E-05 |
| *KRT75* | 3.25 | 4.43E-07 |  | *CYCS* | 1.63 | 1.81E-06 |
| *CST2* | 3.25 | 1.98E-09 |  | *FHOD3* | 1.62 | 0.000173434 |
| *CASP9* | 3.24 | 2.92E-18 |  | *IL18R1* | 1.62 | 0.01723224 |
| *CD55* | 3.24 | 1.35E-30 |  | *LINC01940* | 1.62 | 0.006717844 |
| *FAM87B* | 3.24 | 4.25E-10 |  | *TSPYL2* | 1.62 | 1.90E-07 |
| *CHST1* | 3.22 | 8.25E-10 |  | *KLRD1* | 1.61 | 0.048091097 |
| *FOXQ1* | 3.22 | 2.29E-07 |  | *PBLD* | 1.61 | 8.10E-07 |
| *TNS4* | 3.20 | 4.47E-09 |  | *DNAJC12* | 1.60 | 0.015916814 |
| *PCSK1* | 3.18 | 2.71E-07 |  | *DLX5* | 1.60 | 0.047843122 |
| *NECTIN1* | 3.18 | 1.71E-23 |  | *PTMAP9* | 1.60 | 0.018475129 |
| *SYT12* | 3.18 | 5.21E-11 |  | *DUSP1* | 1.60 | 1.35E-06 |
| *IGFN1* | 3.17 | 1.88E-07 |  | *GABARAPL1* | 1.60 | 7.76E-08 |
| *PTP4A1* | 3.16 | 6.96E-14 |  | *SLC16A4* | 1.60 | 0.002629514 |
| *KCNH1* | 3.15 | 1.78E-10 |  | *PAPPA* | 1.60 | 0.0105666 |
| *APLN* | 3.13 | 1.05E-06 |  | *METRNL* | 1.59 | 0.000324921 |
| *CMSS1* | 3.12 | 1.05E-14 |  | *LOC730101* | 1.59 | 0.007018787 |
| *OAS1* | 3.12 | 9.79E-09 |  | *RDH8* | 1.59 | 0.005213489 |
| *PKIB* | 3.10 | 7.63E-09 |  | *GNG4* | 1.58 | 0.042629852 |
| *IGFBP1* | 3.05 | 1.22E-05 |  | *SYN3* | 1.58 | 0.048915255 |
| *HTRA3* | 3.04 | 2.17E-15 |  | *FIGN* | 1.58 | 0.010590391 |
| *ADRA2C* | 3.03 | 6.63E-12 |  | *MEG9* | 1.58 | 0.008906256 |
| *RHBDF2* | 3.03 | 2.12E-14 |  | *GLA* | 1.57 | 5.56E-11 |
| *FSTL3* | 3.03 | 1.04E-11 |  | *GSTO2* | 1.57 | 0.008055349 |
| *FPR2* | 3.02 | 1.38E-05 |  | *INSRR* | 1.56 | 0.037053677 |
| *CD24* | 3.00 | 6.11E-09 |  | *AMZ1* | 1.55 | 0.026602727 |
| *PDE4B* | 2.96 | 7.41E-07 |  | *SGIP1* | 1.55 | 0.004951893 |
| *PGM2L1* | 2.96 | 2.97E-16 |  | *SLC38A5* | 1.54 | 0.00437423 |
| *PHLDB2* | 2.95 | 3.91E-20 |  | *BNC1* | 1.53 | 0.000237048 |
| *CREM* | 2.95 | 8.90E-19 |  | *NFIL3* | 1.53 | 3.87E-09 |
| *MELTF* | 2.93 | 4.31E-17 |  | *SESN3* | 1.53 | 0.002447549 |
| *TMOD1* | 2.92 | 4.51E-07 |  | *ZDBF2* | 1.52 | 0.00553787 |
| *PNPLA5* | 2.92 | 1.70E-05 |  | *ADAMTS9* | 1.52 | 0.000300313 |
| *SLC2A13* | 2.91 | 6.48E-10 |  | *PGF* | 1.52 | 0.016520721 |
| *GPR4* | 2.91 | 3.65E-09 |  | *SH3PXD2A* | 1.52 | 0.000319147 |
| *HAS2* | 2.90 | 6.79E-13 |  | *NAP1L5* | 1.52 | 0.000723149 |
| *PDE10A* | 2.89 | 1.79E-09 |  | *SLC2A3* | 1.51 | 0.000822287 |
| *IGLON5* | 2.89 | 3.25E-06 |  | *PLA2G2A* | 1.51 | 0.003286347 |
| *FGF10* | 2.89 | 7.15E-05 |  | *UBE2QL1* | 1.51 | 0.002055314 |
| *MIR614* | 2.85 | 8.01E-07 |  | *NXN* | 1.51 | 0.002972392 |
| *RNF122* | 2.85 | 1.05E-15 |  | *NDRG1* | 1.50 | 4.74E-05 |
| *FIBCD1* | 2.85 | 2.42E-09 |  | *C2orf88* | 1.50 | 2.75E-05 |
| *PDE3A* | 2.83 | 2.96E-05 |  | *GALNT12* | 1.50 | 0.001034343 |
| *C17orf58* | 2.83 | 9.84E-33 |  | *TGFBR3* | 1.49 | 0.0002755 |
| *WNT10B* | 2.82 | 2.17E-08 |  | *ST3GAL1* | 1.49 | 2.10E-08 |
| *PDK4* | 2.82 | 9.73E-12 |  | *HS3ST3A1* | 1.49 | 0.000369461 |
| *IL1A* | 2.82 | 5.63E-07 |  | *SIK3* | 1.49 | 0.000367563 |
| *LINC00887* | 2.82 | 4.85E-06 |  | *CACNB2* | 1.48 | 0.046370238 |
| *DPYSL3* | 2.81 | 3.61E-23 |  | *APBA1* | 1.48 | 0.004079426 |
| *LINC00664* | 2.80 | 7.76E-07 |  | *CBLB* | 1.48 | 0.011005579 |
| *ITPRIP* | 2.79 | 4.49E-18 |  | *RAB3A* | 1.47 | 2.84E-05 |
| *SNAI1* | 2.79 | 1.26E-10 |  | *KIF26B* | 1.47 | 0.024526582 |
| *PMEPA1* | 2.79 | 1.21E-16 |  | *SCARA5* | 1.47 | 0.014699458 |
| *DIRAS3* | 2.79 | 2.60E-07 |  | *MSI1* | 1.46 | 0.045719748 |
| *CCK* | 2.78 | 5.33E-05 |  | *TDP2* | 1.46 | 1.63E-06 |
| *PRND* | 2.78 | 3.01E-05 |  | *FGF7* | 1.46 | 0.012577418 |
| *KRT17* | 2.78 | 2.95E-05 |  | *CHST15* | 1.45 | 0.003333112 |
| *C8A* | 2.77 | 1.14E-05 |  | *IL15* | 1.45 | 9.64E-06 |
| *PPDPFL* | 2.75 | 0.000258703 |  | *PRG4* | 1.45 | 0.023088744 |
| *KCNK5* | 2.72 | 1.49E-05 |  | *UAP1* | 1.45 | 1.34E-06 |
| *ATP1B3* | 2.72 | 5.10E-15 |  | *ODC1* | 1.45 | 0.000145404 |
| *IL33* | 2.72 | 3.82E-05 |  | *GAS1* | 1.44 | 0.000474365 |
| *TCIM* | 2.71 | 2.99E-06 |  | *DNAJB9* | 1.44 | 0.000406144 |
| *TMEM151A* | 2.71 | 1.07E-06 |  | *IL4R* | 1.44 | 0.000396705 |
| *ATOH7* | 2.71 | 1.36E-05 |  | *ENTPD7* | 1.44 | 0.026897485 |
| *BDKRB1* | 2.70 | 5.21E-08 |  | *ABCA4* | 1.43 | 0.041349496 |
| *ANXA10* | 2.70 | 2.57E-07 |  | *GAB2* | 1.43 | 0.000158429 |
| *BEX1* | 2.68 | 6.25E-06 |  | *LINC01588* | 1.43 | 0.022348186 |
| *CASZ1* | 2.68 | 4.86E-07 |  | *EPOP* | 1.43 | 0.001187525 |
| *CYP19A1* | 2.68 | 7.70E-07 |  | *CLMP* | 1.42 | 0.024878478 |
| *EXOC3L2* | 2.67 | 6.91E-06 |  | *DLL1* | 1.42 | 0.01933454 |
| *PDE4C* | 2.67 | 3.48E-07 |  | *TTYH2* | 1.41 | 0.017081024 |
| *NAMPT* | 2.66 | 3.24E-13 |  | *MEG3* | 1.41 | 0.017007687 |
| *SV2B* | 2.66 | 0.000106739 |  | *GPC2* | 1.41 | 0.020093522 |
| *WFDC21P* | 2.66 | 0.000813079 |  | *TRAF4* | 1.41 | 1.80E-05 |
| *AADACL2* | 2.66 | 5.01E-05 |  | *MTFP1* | 1.41 | 0.005955292 |
| *HYAL1* | 2.63 | 2.52E-14 |  | *ABTB2* | 1.41 | 0.011521879 |
| *NCCRP1* | 2.61 | 3.96E-05 |  | *SPDL1* | 1.40 | 0.000417451 |
| *TMEM217* | 2.61 | 1.18E-10 |  | *TRIM29* | 1.40 | 0.027164728 |
| *WNT5A* | 2.61 | 5.56E-11 |  | *HSPD1* | 1.40 | 2.24E-05 |
| *KCNG1* | 2.60 | 5.51E-14 |  | *ABL1* | 1.40 | 0.001778489 |
| *HMX3* | 2.58 | 0.000355168 |  | *MEG8* | 1.40 | 0.025995813 |
| *GPR157* | 2.57 | 1.20E-05 |  | *CYRIA* | 1.39 | 0.037942912 |
| *CREB5* | 2.57 | 1.02E-08 |  | *ZBTB16* | 1.39 | 0.007358473 |
| *TLNRD1* | 2.57 | 4.70E-22 |  | *CARD8-AS1* | 1.39 | 6.95E-05 |
| *DEFB1* | 2.57 | 0.000140451 |  | *HBEGF* | 1.38 | 0.001838064 |
| *LINC02274* | 2.56 | 0.000103499 |  | *DUSP5* | 1.38 | 0.00024244 |
| *PITPNM1* | 2.56 | 7.61E-27 |  | *FKBP4* | 1.38 | 6.47E-06 |
| *RASD1* | 2.55 | 8.67E-08 |  | *MAPKAPK2* | 1.38 | 4.35E-06 |
| *KSR1* | 2.55 | 8.75E-06 |  | *CDK2AP2* | 1.37 | 0.000103499 |
| *GADD45G* | 2.54 | 1.67E-06 |  | *CENPN* | 1.36 | 1.82E-06 |
| *B4GALT1* | 2.54 | 1.78E-15 |  | *NFATC4* | 1.36 | 0.000364716 |
| *BDNF* | 2.54 | 6.81E-08 |  | *SDF2L1* | 1.36 | 0.007589073 |
| *SUCNR1* | 2.53 | 8.84E-05 |  | *AGPS* | 1.35 | 0.000621107 |
| *ARFGEF3* | 2.53 | 0.001759224 |  | *CYGB* | 1.35 | 0.001251001 |
| *MT1L* | 2.53 | 2.13E-06 |  | *LGALS9* | 1.35 | 0.003317465 |
| *C12orf60* | 2.53 | 1.04E-06 |  | *CITED1* | 1.35 | 0.037698851 |
| *CXCR4* | 2.52 | 7.18E-06 |  | *LOC100287387* | 1.34 | 0.027560741 |
| *RFK* | 2.52 | 5.27E-11 |  | *CUL4B* | 1.34 | 0.000257561 |
| *IL1B* | 2.52 | 3.49E-06 |  | *CISH* | 1.34 | 0.003336355 |
| *LIPG* | 2.51 | 6.86E-06 |  | *CHST6* | 1.34 | 0.046254303 |
| *ISG20* | 2.51 | 3.76E-06 |  | *FAM122C* | 1.34 | 0.001232551 |
| *BMP8B* | 2.51 | 9.08E-05 |  | *PENK* | 1.34 | 0.027108929 |
| *S1PR1* | 2.50 | 2.61E-07 |  | *MAP7* | 1.34 | 0.009152838 |
| *RAMP1* | 2.50 | 4.39E-08 |  | *PBX1* | 1.34 | 0.000210414 |
| *BCL2L11* | 2.50 | 2.19E-13 |  | *AKAP6* | 1.34 | 0.003200592 |
| *TUBB3* | 2.50 | 1.33E-17 |  | *AURKA* | 1.33 | 1.35E-05 |
| *FOS* | 2.50 | 1.39E-15 |  | *RAB21* | 1.33 | 0.000363935 |
| *ITK* | 2.50 | 0.0002336 |  | *AKAP13* | 1.33 | 0.005543966 |
| *KCNJ8* | 2.49 | 2.48E-13 |  | *KPNA2* | 1.32 | 1.66E-06 |
| *VASH2* | 2.49 | 9.64E-06 |  | *DNAJA1* | 1.32 | 6.70E-05 |
| *CXCL8* | 2.49 | 3.37E-08 |  | *IL1R1* | 1.32 | 0.016736565 |
| *MT2A* | 2.47 | 7.59E-09 |  | *PAX8* | 1.32 | 0.015347156 |
| *NECTIN4* | 2.47 | 9.67E-06 |  | *ENO3* | 1.32 | 0.026796395 |
| *KCNA10* | 2.46 | 0.000655926 |  | *MEX3A* | 1.31 | 0.012436666 |
| *TGM2* | 2.46 | 9.93E-10 |  | *WIPI1* | 1.31 | 1.18E-05 |
| *C11orf96* | 2.45 | 1.14E-08 |  | *BLOC1S3* | 1.31 | 1.14E-07 |
| *BMP8A* | 2.45 | 3.76E-05 |  | *JUND* | 1.31 | 0.000892695 |
| *SULT4A1* | 2.44 | 8.28E-06 |  | *HSPE1* | 1.31 | 0.000796776 |
| *LIF* | 2.42 | 0.000268509 |  | *LARP4* | 1.31 | 0.003895215 |
| *CHSY1* | 2.42 | 7.38E-09 |  | *GMPPB* | 1.31 | 5.95E-05 |
| *MAP3K14-AS1* | 2.41 | 1.50E-05 |  | *JUP* | 1.30 | 0.000355168 |
| *KBTBD8* | 2.41 | 4.59E-09 |  | *PDLIM4* | 1.30 | 0.001077458 |
| *ISM1* | 2.41 | 0.00024389 |  | *CRY1* | 1.30 | 0.000243114 |
| *RCSD1* | 2.40 | 5.77E-05 |  | *TOB1* | 1.30 | 0.000114626 |
| *SPON1* | 2.40 | 1.14E-06 |  | *ADAM12* | 1.30 | 0.027213646 |
| *SLC1A2* | 2.39 | 0.000189436 |  | *LARGE1* | 1.30 | 0.001111235 |
| *ACTL8* | 2.39 | 0.000485177 |  | *PLAT* | 1.29 | 0.000682006 |
| *PAX1* | 2.39 | 0.00091642 |  | *ESYT2* | 1.29 | 0.000396705 |
| *TUBB2A* | 2.39 | 1.53E-07 |  | *SELENOK* | 1.29 | 0.002800308 |
| *C15orf48* | 2.38 | 0.000290194 |  | *HMOX1* | 1.29 | 0.00437423 |
| *IFI30* | 2.37 | 5.84E-08 |  | *BZW2* | 1.29 | 6.24E-06 |
| *B3GNT5* | 2.37 | 4.32E-07 |  | *TRABD2A* | 1.28 | 0.008863102 |
| *GFPT2* | 2.37 | 3.82E-08 |  | *PPIAP46* | 1.28 | 0.049528 |
| *PPP1R3C* | 2.36 | 1.66E-07 |  | *OXSR1* | 1.28 | 5.50E-05 |
| *KCNE4* | 2.36 | 2.74E-07 |  | *MT1X* | 1.28 | 0.00786273 |
| *PTGES* | 2.36 | 2.10E-13 |  | *PTGFRN* | 1.28 | 0.005073146 |
| *CXCL2* | 2.35 | 1.95E-05 |  | *ARRDC2* | 1.27 | 0.004001886 |
| *LINC00322* | 2.35 | 0.00017767 |  | *UBASH3B* | 1.27 | 0.008356588 |
| *PER1* | 2.35 | 4.67E-11 |  | *STK26* | 1.27 | 0.013328548 |
| *RELT* | 2.35 | 3.24E-07 |  | *TIMP3* | 1.27 | 9.33E-07 |
| *CXCL3* | 2.35 | 3.04E-05 |  | *CRLF1* | 1.27 | 0.028176406 |
| *MASP1* | 2.35 | 1.47E-06 |  | *SPTSSA* | 1.27 | 1.13E-05 |
| *MAPK4* | 2.34 | 0.000769067 |  | *LOC100379224* | 1.26 | 0.012491821 |
| *SPOCK2* | 2.34 | 3.43E-06 |  | *FGF18* | 1.26 | 0.029490347 |
| *PRL* | 2.34 | 0.000305704 |  | *EFR3B* | 1.26 | 0.038144841 |
| *RAB20* | 2.33 | 1.47E-17 |  | *RCE1* | 1.26 | 1.95E-05 |
| *LINC00673* | 2.33 | 1.39E-06 |  | *LY6K* | 1.26 | 0.042575068 |
| *IGSF21* | 2.33 | 0.00142371 |  | *PHF10* | 1.25 | 0.015253361 |
| *DLGAP1* | 2.32 | 0.000396544 |  | *RNF152* | 1.25 | 0.024526582 |
| *SPAG4* | 2.32 | 6.33E-06 |  | *LINC00941* | 1.25 | 0.003539364 |
| *FER1L6* | 2.32 | 0.000601532 |  | *THBS2* | 1.25 | 0.040229099 |
| *MYPN* | 2.32 | 4.74E-07 |  | *KLF4* | 1.25 | 1.02E-05 |
| *DUSP8* | 2.31 | 7.32E-08 |  | *ZNF281* | 1.24 | 0.01893133 |
| *EDNRB* | 2.31 | 1.51E-05 |  | *GNAL* | 1.24 | 0.020224258 |
| *TRABD2B* | 2.31 | 1.35E-07 |  | *SLC31A1* | 1.24 | 0.000742523 |
| *SV2C* | 2.30 | 0.001142588 |  | *TGIF1* | 1.24 | 3.99E-06 |
| *FOSB* | 2.30 | 3.17E-09 |  | *CELSR3* | 1.23 | 0.046451493 |
| *GAP43* | 2.30 | 7.63E-05 |  | *VEGFD* | 1.23 | 0.005589301 |
| *LETM2* | 2.30 | 2.88E-06 |  | *MIR635* | 1.22 | 0.048709144 |
| *PGBD5* | 2.29 | 7.51E-06 |  | *CSRNP1* | 1.22 | 0.000136359 |
| *FOXE1* | 2.28 | 3.98E-05 |  | *SHISAL1* | 1.22 | 0.01933454 |
| *TMEM71* | 2.28 | 1.96E-06 |  | *SIN3B* | 1.22 | 0.000925552 |
| *SOWAHD* | 2.27 | 0.000315825 |  | *ARL4A* | 1.22 | 0.000671601 |
| *PRDM1* | 2.27 | 8.80E-06 |  | *HYOU1* | 1.22 | 0.024588619 |
| *JUNB* | 2.26 | 7.70E-09 |  | *YRDC* | 1.21 | 0.00069507 |
| *IMPA2* | 2.26 | 1.89E-09 |  | *TMEM158* | 1.21 | 0.005543966 |
| *NFATC2* | 2.26 | 0.001226544 |  | *SIAH1* | 1.21 | 0.001541961 |
| *AGAP11* | 2.25 | 0.000359719 |  | *FNDC3A* | 1.21 | 0.022817412 |
| *CDH24* | 2.25 | 7.82E-06 |  | *SIPA1L2* | 1.21 | 0.030524334 |
| *SORL1* | 2.25 | 3.13E-05 |  | *LDLRAD4* | 1.21 | 0.001320188 |
| *ZNF703* | 2.24 | 4.75E-08 |  | *FABP5* | 1.21 | 0.023516126 |
| *CFAP69* | 2.24 | 6.10E-08 |  | *TMC7* | 1.20 | 0.048915255 |
| *C5AR2* | 2.24 | 0.002178741 |  | *RNF113A* | 1.20 | 0.003557232 |
| *NR4A3* | 2.24 | 6.97E-05 |  | *GPR3* | 1.20 | 0.040014512 |
| *LAMC3* | 2.23 | 0.000204623 |  | *MXI1* | 1.19 | 0.031844744 |
| *SFTPB* | 2.22 | 0.000299483 |  | *KALRN* | 1.19 | 0.039526888 |
| *TP53AIP1* | 2.22 | 0.004714992 |  | *PTHLH* | 1.19 | 0.017081426 |
| *XYLT1* | 2.21 | 1.24E-06 |  | *TMEM51* | 1.19 | 0.019845689 |
| *ABLIM1* | 2.21 | 7.75E-07 |  | *CXCL1* | 1.19 | 0.040217538 |
| *FOXF1* | 2.21 | 0.001126269 |  | *NCS1* | 1.19 | 2.88E-07 |
| *GRAMD1C* | 2.20 | 2.07E-05 |  | *GNA12* | 1.18 | 0.012358459 |
| *CELA2B* | 2.19 | 0.000892695 |  | *ATP2A2* | 1.18 | 0.020185428 |
| *AJAP1* | 2.19 | 0.000264793 |  | *ECE1* | 1.18 | 0.00064781 |
| *RADIL* | 2.18 | 4.70E-12 |  | *TOMM40* | 1.18 | 0.000919629 |
| *ZNF331* | 2.18 | 1.62E-06 |  | *TAF13* | 1.17 | 0.026192052 |
| *THAP2* | 2.17 | 6.40E-10 |  | *MAP3K4* | 1.17 | 0.017145146 |
| *TYMP* | 2.17 | 4.10E-08 |  | *ZNF521* | 1.17 | 0.017592865 |
| *C1QL2* | 2.16 | 0.002954487 |  | *TMUB1* | 1.17 | 0.03740958 |
| *LRFN4* | 2.16 | 4.23E-12 |  | *GOT1* | 1.17 | 0.000308677 |
| *DNER* | 2.16 | 0.000258171 |  | *CHD7* | 1.15 | 0.032012478 |
| *HES4* | 2.16 | 0.000273136 |  | *BIN1* | 1.15 | 0.003310672 |
| *QRFP* | 2.15 | 0.00035808 |  | *DYRK3* | 1.15 | 4.58E-05 |
| *AADAC* | 2.15 | 4.57E-06 |  | *RELL1* | 1.15 | 0.00373454 |
| *AGPAT4* | 2.15 | 8.80E-06 |  | *CDH4* | 1.15 | 0.045278336 |
| *ARRDC3* | 2.15 | 1.82E-07 |  | *CRABP2* | 1.15 | 0.022431583 |
| *GALNT15* | 2.14 | 9.49E-06 |  | *MFSD12* | 1.15 | 0.000536923 |
| *ELOVL7* | 2.14 | 0.000114626 |  | *RRN3* | 1.14 | 0.000359719 |
| *TENT5C* | 2.14 | 0.000382017 |  | *CYTH1* | 1.14 | 2.32E-05 |
| *GALNT2* | 2.14 | 2.09E-08 |  | *KIF21A* | 1.14 | 0.013085698 |
| *FGF2* | 2.14 | 5.80E-08 |  | *ESPL1* | 1.14 | 0.023783109 |
| *CGA* | 2.12 | 0.008349517 |  | *SPTLC3* | 1.14 | 0.004514269 |
| *SLCO4A1* | 2.12 | 0.000204623 |  | *FASTKD1* | 1.14 | 9.29E-06 |
| *INHBA* | 2.10 | 0.001678306 |  | *ZSCAN5A* | 1.14 | 0.008382806 |
| *CFAP58-DT* | 2.10 | 0.003142105 |  | *SYAP1* | 1.13 | 0.000103389 |
| *TSKU* | 2.10 | 8.38E-08 |  | *EAF1* | 1.13 | 0.007589073 |
| *HES2* | 2.09 | 0.007619139 |  | *BNIP3L* | 1.13 | 0.005972661 |
| *AP5B1* | 2.09 | 3.84E-10 |  | *EIF4E* | 1.13 | 0.000547519 |
| *CSGALNACT1* | 2.09 | 1.12E-10 |  | *WARS1* | 1.12 | 4.49E-05 |
| *PXDNL* | 2.09 | 0.003290414 |  | *CDR2L* | 1.12 | 0.000253325 |
| *SEPTIN6* | 2.09 | 6.60E-06 |  | *NCALD* | 1.12 | 0.015845811 |
| *DDIT4* | 2.08 | 7.90E-08 |  | *OGDH* | 1.12 | 0.000726239 |
| *SPINK1* | 2.07 | 0.001134305 |  | *RFLNB* | 1.12 | 0.011433625 |
| *SLC7A5* | 2.06 | 2.62E-06 |  | *NAA50* | 1.12 | 0.002071417 |
| *SYNDIG1* | 2.06 | 5.33E-05 |  | *ZBTB21* | 1.12 | 0.022507866 |
| *SHC4* | 2.06 | 0.000197522 |  | *CAMKK1* | 1.11 | 0.028954979 |
| *RHOB* | 2.05 | 2.57E-12 |  | *SMYD4* | 1.11 | 9.16E-05 |
| *DOC2B* | 2.05 | 0.010726514 |  | *HSPH1* | 1.11 | 0.012193975 |
| *PIGA* | 2.05 | 3.76E-06 |  | *DKK1* | 1.11 | 0.007948265 |
| *PSD4* | 2.04 | 2.44E-05 |  | *HES1* | 1.11 | 0.017356097 |
| *ZMAT4* | 2.04 | 0.007358473 |  | *SEC23B* | 1.11 | 0.001480354 |
| *SPSB4* | 2.03 | 0.011457951 |  | *USP35* | 1.11 | 0.003917772 |
| *CTNNB1* | 2.02 | 9.80E-10 |  | *ELL2* | 1.10 | 0.01164505 |
| *KLHDC7B* | 2.01 | 0.000457693 |  | *EIF2AK3* | 1.10 | 0.042596712 |
| *CD14* | 2.01 | 8.12E-06 |  | *TPI1* | 1.10 | 0.026602727 |
| *IGF1* | 2.01 | 8.61E-06 |  | *KDM7A-DT* | 1.10 | 0.004131575 |
| *NIM1K* | 2.01 | 0.000719598 |  | *JMJD6* | 1.10 | 0.001673141 |
| *TFRC* | 2.00 | 0.000270213 |  | *ZDHHC14* | 1.09 | 0.003480859 |
| *VMO1* | 2.00 | 0.00114907 |  | *MTHFD1L* | 1.09 | 0.00031242 |
| *KYNU* | 1.99 | 0.002655456 |  | *DGKD* | 1.09 | 0.01989904 |
| *CCL20* | 1.99 | 0.001164398 |  | *PAICS* | 1.09 | 0.042596712 |
| *PCSK1N* | 1.99 | 0.001352687 |  | *MARK4* | 1.09 | 0.000277909 |
| *GRIA3* | 1.99 | 6.97E-06 |  | *C5AR1* | 1.09 | 0.03511211 |
| *FLVCR2* | 1.98 | 9.51E-06 |  | *SSBP3* | 1.09 | 0.019028173 |
| *C10orf90* | 1.97 | 1.95E-06 |  | *PHLDA1* | 1.09 | 0.019860707 |
| *GLP2R* | 1.97 | 0.001759174 |  | *APOLD1* | 1.09 | 0.009701183 |
| *MEDAG* | 1.97 | 2.05E-13 |  | *MOSPD1* | 1.09 | 0.00214579 |
| *TLE3* | 1.97 | 2.37E-06 |  | *LXN* | 1.09 | 0.041164893 |
| *ESYT3* | 1.96 | 0.001589368 |  | *SCO2* | 1.09 | 0.039256381 |
| *MT1M* | 1.96 | 0.002677625 |  | *KDM6B* | 1.09 | 0.034995907 |
| *ATF3* | 1.96 | 1.10E-11 |  | *MPHOSPH6* | 1.08 | 0.000806563 |
| *BDKRB2* | 1.96 | 0.003446315 |  | *HYAL2* | 1.08 | 0.000386709 |
| *PRKAR1A* | 1.95 | 2.26E-09 |  | *MYD88* | 1.08 | 0.00109209 |
| *IGFBP4* | 1.95 | 7.61E-05 |  | *HSPA5* | 1.07 | 0.025083466 |
| *ACKR3* | 1.95 | 3.37E-09 |  | *ARSG* | 1.07 | 0.026060799 |
| *PTGS2* | 1.95 | 0.002646187 |  | *PTPN1* | 1.07 | 0.003205386 |
| *CACNA1G-AS1* | 1.95 | 0.012323586 |  | *ZNF672* | 1.07 | 6.31E-05 |
| *SIX3* | 1.95 | 0.017081426 |  | *TMED5* | 1.06 | 0.000295381 |
| *DAW1* | 1.93 | 0.000435546 |  | *DIP2C* | 1.06 | 0.003877089 |
| *TACC2* | 1.93 | 1.11E-12 |  | *ADAT3* | 1.06 | 0.010692364 |
| *INSYN2B* | 1.93 | 0.005949911 |  | *SMIM12* | 1.06 | 0.006601133 |
| *SLC22A11* | 1.92 | 0.007320291 |  | *UBE2S* | 1.06 | 0.036347201 |
| *GFOD1* | 1.92 | 2.96E-05 |  | *PPME1* | 1.06 | 0.01933454 |
| *HCRTR1* | 1.92 | 0.01933454 |  | *CTSL* | 1.06 | 0.001104936 |
| *CYFIP2* | 1.92 | 0.000314227 |  | *PHC2* | 1.05 | 0.00344748 |
| *EPB41L3* | 1.92 | 1.17E-06 |  | *HDAC4* | 1.05 | 0.017592865 |
| *ALPK3* | 1.92 | 0.000475541 |  | *KLF9* | 1.05 | 0.02592823 |
| *BTBD11* | 1.91 | 0.005708184 |  | *BAG1* | 1.05 | 0.037146255 |
| *VEGFA* | 1.91 | 1.35E-05 |  | *AMMECR1* | 1.05 | 0.032230683 |
| *PITX1* | 1.91 | 0.005922444 |  | *MARCKSL1* | 1.04 | 0.005394022 |
| *CREB3L2* | 1.90 | 0.000121473 |  | *INO80C* | 1.04 | 0.027478268 |
| *FAM124A* | 1.89 | 0.000642114 |  | *TXNDC11* | 1.03 | 0.000913214 |
| *ADAMTS6* | 1.89 | 0.001528176 |  | *ARHGEF40* | 1.03 | 0.012683586 |
| *GASK1A* | 1.89 | 5.39E-05 |  | *BAG3* | 1.03 | 0.000400725 |
| *JPH1* | 1.89 | 0.007115825 |  | *NFKBIA* | 1.03 | 0.005334078 |
| *PCSK9* | 1.89 | 2.77E-13 |  | *FARSA* | 1.03 | 0.000406144 |
| *FGFR1* | 1.88 | 1.06E-06 |  | *MANF* | 1.03 | 0.007496923 |
| *HAL* | 1.87 | 0.0058507 |  | *SLC33A1* | 1.03 | 0.006109415 |
| *SH3PXD2B* | 1.87 | 6.98E-05 |  | *MAGEH1* | 1.03 | 0.001247802 |
| *PKDCC* | 1.87 | 0.000264793 |  | *AGMAT* | 1.03 | 0.032693139 |
| *C9orf72* | 1.86 | 2.02E-05 |  | *CEBPD* | 1.02 | 0.009348043 |
| *CLCF1* | 1.86 | 5.28E-10 |  | *SLC22A3* | 1.02 | 0.034729275 |
| *CDC6* | 1.86 | 1.04E-06 |  | *EGLN1* | 1.02 | 0.028001561 |
| *ATOH8* | 1.85 | 2.29E-07 |  | *KAZN* | 1.02 | 0.02040943 |
| *CPEB4* | 1.85 | 1.02E-05 |  | *ID1* | 1.02 | 0.048413276 |
| *MOB1B* | 1.85 | 1.66E-07 |  | *RAPGEF1* | 1.01 | 0.028433052 |
| *ZC2HC1C* | 1.85 | 0.007638358 |  | *CCT2* | 1.01 | 0.000522452 |
| *RPRM* | 1.84 | 0.014562199 |  | *FAM214B* | 1.01 | 1.65E-05 |
| *PID1* | 1.84 | 4.32E-05 |  | *DNAJC1* | 1.01 | 0.009873377 |
| *LINC01121* | 1.84 | 0.00184402 |  | *LRRC59* | 1.01 | 0.007093368 |
| *SMTNL2* | 1.83 | 0.023366655 |  | *TBX3* | 1.01 | 0.046811538 |
| *FXYD6* | 1.83 | 0.023213637 |  | *SMPDL3A* | 1.01 | 0.029016107 |
| *LMLN2* | 1.83 | 0.007227563 |  | *TMEM198* | 1.01 | 0.017850856 |
| *RHOU* | 1.83 | 3.27E-07 |  | *AMPD2* | 1.00 | 0.032817387 |
| *KLHL26* | 1.83 | 1.68E-08 |  | *PGK1* | 1.00 | 0.046368645 |
| *TWIST2* | 1.83 | 9.82E-11 |  | *CLUH* | 1.00 | 0.003086945 |
| *PALMD* | 1.83 | 6.29E-06 |  | *PFKP* | 1.00 | 0.042362006 |
| *LONRF2* | 1.82 | 0.003605484 |  | *ST3GAL4* | 1.00 | 0.016402291 |

**Supplementary Table 4.** Downregulated genes by cAMP in SC adipocytes. Log_2_ FC ≤-1, Adjective P value <0.05.

| **Symbol** | **Log2 Fold Change** | **Adj. P Value** |  | **Symbol** | **Log2 Fold Change** | **Adj. P Value** |
| --- | --- | --- | --- | --- | --- | --- |
| *SERPINE1* | -4.70 | 7.21E-33 |  | *PLA2G5* | -1.51 | 3.76E-02 |
| *PTX3* | -4.70 | 1.81E-36 |  | *N4BP2L1* | -1.51 | 1.42E-05 |
| *B3GALT2* | -4.70 | 2.51E-25 |  | *MAP4K3* | -1.51 | 0.002049239 |
| *RPSAP52* | -4.23 | 2.61E-17 |  | *LOC100129034* | -1.50 | 0.000479264 |
| *DRD1* | -4.22 | 1.79E-13 |  | *ARRB1* | -1.50 | 1.54498E-05 |
| *ANKRD1* | -3.97 | 9.33E-12 |  | *RXRG* | -1.50 | 0.049218112 |
| *ADRA2A* | -3.68 | 3.49E-10 |  | *FMO4* | -1.50 | 1.75771E-05 |
| *LINC01085* | -3.67 | 1.27E-10 |  | *MBP* | -1.49 | 3.59E-04 |
| *KRTAP1-5* | -3.66 | 3.93E-12 |  | *RIMS3* | -1.49 | 9.33E-04 |
| *BMP4* | -3.66 | 4.90E-16 |  | *CHST7* | -1.48 | 1.16E-05 |
| *ADAMTS8* | -3.62 | 8.17E-13 |  | *MAN1C1* | -1.48 | 4.11E-03 |
| *PPL* | -3.53 | 5.10E-22 |  | *PLAU* | -1.48 | 4.94E-03 |
| *PLCE1-AS1* | -3.50 | 6.97E-12 |  | *CYP4F22* | -1.48 | 0.002251249 |
| *ST8SIA1* | -3.49 | 2.05E-26 |  | *TJP2* | -1.48 | 6.42E-03 |
| *ELOVL2* | -3.48 | 1.12E-07 |  | *ADM* | -1.48 | 0.000589896 |
| *HMGA2-AS1* | -3.43 | 2.65E-09 |  | *THRB* | -1.48 | 0.000475541 |
| *HECW2* | -3.39 | 4.62E-09 |  | *SCHIP1* | -1.48 | 0.000355168 |
| *NTF3* | -3.25 | 2.23E-12 |  | *MITF* | -1.47 | 4.20425E-05 |
| *GPAM* | -3.23 | 8.50E-18 |  | *MKX* | -1.47 | 2.02E-02 |
| *MEOX2* | -3.21 | 1.50E-10 |  | *TNFAIP8* | -1.47 | 4.53E-04 |
| *TRIB2* | -3.19 | 1.61E-12 |  | *CCDC85A* | -1.47 | 1.45417E-05 |
| *ANK3* | -3.12 | 3.95E-08 |  | *CRYBG1* | -1.47 | 0.003539777 |
| *FMO2* | -3.10 | 1.82E-07 |  | *KCNB1* | -1.46 | 3.21E-02 |
| *EEPD1* | -3.10 | 3.91E-22 |  | *TNFAIP8L1* | -1.46 | 0.000950759 |
| *ATP8B1* | -3.07 | 6.46E-10 |  | *FLNC* | -1.46 | 0.009910856 |
| *RGS4* | -2.99 | 9.96E-07 |  | *TRIL* | -1.46 | 0.006174411 |
| *VGLL3* | -2.97 | 2.52E-09 |  | *LYPLAL1* | -1.46 | 2.14E-04 |
| *C1orf198* | -2.96 | 1.82E-16 |  | *CEBPA* | -1.46 | 8.01E-03 |
| *NCKAP5* | -2.96 | 1.25E-11 |  | *FANCE* | -1.46 | 0.00121178 |
| *EVA1A* | -2.90 | 5.53E-14 |  | *CAND2* | -1.46 | 0.020830306 |
| *PTGIR* | -2.90 | 7.31E-15 |  | *MAP2* | -1.46 | 0.031250635 |
| *MAP3K7CL* | -2.89 | 1.69E-13 |  | *CDC42EP1* | -1.45 | 1.34E-08 |
| *CX3CL1* | -2.89 | 1.36E-05 |  | *DDX60L* | -1.45 | 0.00464779 |
| *MSC* | -2.86 | 7.65E-21 |  | *TCEA3* | -1.45 | 0.003633751 |
| *RGS3* | -2.84 | 1.21E-16 |  | *CALHM5* | -1.45 | 0.00437423 |
| *MAFB* | -2.83 | 2.68E-12 |  | *LINC00886* | -1.45 | 0.007592704 |
| *NAV3* | -2.82 | 6.33E-12 |  | *AHNAK2* | -1.45 | 0.001484775 |
| *KIT* | -2.82 | 1.20E-06 |  | *GUCY1B1* | -1.44 | 2.24E-03 |
| *CDC42EP3* | -2.81 | 1.30E-09 |  | *CHADL* | -1.44 | 1.52E-02 |
| *PRR15* | -2.80 | 2.16E-08 |  | *CDT1* | -1.44 | 1.24E-02 |
| *CCN2* | -2.79 | 2.38E-08 |  | *SYTL3* | -1.44 | 0.002760532 |
| *SLC16A12* | -2.78 | 5.76E-05 |  | *SYNE1* | -1.44 | 0.040472782 |
| *DBP* | -2.77 | 4.74E-07 |  | *TNFRSF14* | -1.44 | 3.01029E-05 |
| *ACKR4* | -2.70 | 3.51E-16 |  | *AP3M2* | -1.44 | 0.000690473 |
| *LINC02709* | -2.70 | 2.03E-08 |  | *COLGALT2* | -1.44 | 0.017162374 |
| *LZTS1* | -2.68 | 7.34E-06 |  | *TRAK1* | -1.44 | 0.001091224 |
| *GAS6-AS1* | -2.65 | 3.10E-06 |  | *TOLLIP-AS1* | -1.44 | 0.028176406 |
| *OXTR* | -2.65 | 4.53E-06 |  | *INHBB* | -1.43 | 1.65E-02 |
| *POU2F2* | -2.65 | 5.73E-06 |  | *AARD* | -1.43 | 0.018247591 |
| *CCN1* | -2.65 | 4.99E-10 |  | *RASSF9* | -1.43 | 2.17E-02 |
| *ZFHX4-AS1* | -2.64 | 5.10E-08 |  | *LGALS12* | -1.43 | 0.019576565 |
| *STING1* | -2.64 | 1.02E-23 |  | *LOC145694* | -1.43 | 2.21E-02 |
| *DEPTOR* | -2.63 | 9.98E-13 |  | *XAF1* | -1.43 | 1.91E-02 |
| *CNTN3* | -2.61 | 8.78E-06 |  | *NR2F2* | -1.43 | 3.55656E-05 |
| *FGF9* | -2.60 | 3.56E-05 |  | *HOGA1* | -1.42 | 2.42E-02 |
| *LOXL4* | -2.60 | 1.06E-29 |  | *DDO* | -1.42 | 4.80E-02 |
| *CCDC81* | -2.57 | 1.58E-07 |  | *TFEB* | -1.42 | 0.000314875 |
| *NEDD9* | -2.56 | 2.86E-11 |  | *PRKAR2A-AS1* | -1.42 | 0.025931835 |
| *SLC9A9* | -2.55 | 2.45E-14 |  | *LRRC75A* | -1.42 | 0.018099677 |
| *THRSP* | -2.52 | 9.03E-06 |  | *SLC2A4* | -1.41 | 0.046398271 |
| *IL34* | -2.52 | 1.25E-07 |  | *PPP1R13L* | -1.41 | 0.00125741 |
| *SEC14L5* | -2.51 | 8.56E-06 |  | *CLDN11* | -1.41 | 5.77444E-05 |
| *RGS7* | -2.50 | 7.37E-06 |  | *LYRM9* | -1.41 | 0.000634553 |
| *KIAA0040* | -2.50 | 1.54E-06 |  | *VWA5A* | -1.41 | 1.80488E-06 |
| *IFIT3* | -2.49 | 2.59E-11 |  | *EFNB1* | -1.41 | 1.63E-04 |
| *TENT5B* | -2.48 | 8.29E-07 |  | *SAMD4A* | -1.41 | 8.88E-03 |
| *TSPAN15* | -2.47 | 1.27E-05 |  | *OPRL1* | -1.41 | 0.016736565 |
| *IL16* | -2.47 | 2.17E-08 |  | *GAB1* | -1.41 | 0.010290628 |
| *S1PR5* | -2.46 | 1.94E-05 |  | *LIFR-AS1* | -1.40 | 0.008008076 |
| *TRHDE-AS1* | -2.46 | 6.95E-07 |  | *EPG5* | -1.40 | 1.30E-02 |
| *DDAH1* | -2.45 | 7.86E-09 |  | *LINC01094* | -1.40 | 0.039571877 |
| *IFIT1* | -2.45 | 8.72E-13 |  | *CABYR* | -1.40 | 0.005832224 |
| *NPAS1* | -2.45 | 1.67E-08 |  | *HAVCR2* | -1.40 | 0.0240893 |
| *LMOD1* | -2.44 | 1.38E-08 |  | *SOX8* | -1.40 | 0.022348186 |
| *CACNB4* | -2.44 | 6.01E-07 |  | *MMP11* | -1.40 | 0.037921496 |
| *CMAHP* | -2.42 | 1.58E-09 |  | *PBX4* | -1.40 | 3.78E-02 |
| *HMGA2* | -2.41 | 6.25E-06 |  | *DAB2* | -1.40 | 6.81E-03 |
| *DTX4* | -2.41 | 1.08E-05 |  | *APOL6* | -1.40 | 0.002862469 |
| *RTL3* | -2.39 | 1.18E-03 |  | *GHR* | -1.39 | 6.08E-05 |
| *KCTD16* | -2.38 | 4.05E-05 |  | *TMEM30B* | -1.39 | 0.022697306 |
| *NRK* | -2.38 | 7.45E-05 |  | *SH2D5* | -1.39 | 4.50E-02 |
| *CYTH3* | -2.37 | 5.52E-09 |  | *MAP4K3-DT* | -1.39 | 2.17E-03 |
| *PTGER4* | -2.37 | 2.84E-06 |  | *F2R* | -1.39 | 0.000214787 |
| *IRAG1* | -2.37 | 3.64E-06 |  | *FBLIM1* | -1.38 | 3.2599E-05 |
| *KIAA1755* | -2.36 | 3.56E-05 |  | *CCDC9B* | -1.38 | 0.003713451 |
| *KRT34* | -2.36 | 1.64E-05 |  | *SELPLG* | -1.38 | 1.15E-03 |
| *PDE5A* | -2.35 | 3.18E-06 |  | *SMURF2* | -1.38 | 0.018502726 |
| *KCNS2* | -2.34 | 3.55E-04 |  | *CCBE1* | -1.38 | 0.001165456 |
| *GATA6-AS1* | -2.33 | 4.57E-05 |  | *PTK2B* | -1.37 | 1.21E-04 |
| *BBC3* | -2.31 | 1.06E-10 |  | *PPARGC1B* | -1.37 | 0.002055237 |
| *USP53* | -2.31 | 5.06E-07 |  | *GLS* | -1.37 | 0.012098202 |
| *JCAD* | -2.31 | 4.22E-05 |  | *GPC4* | -1.37 | 7.74815E-07 |
| *NKX3-2* | -2.31 | 4.36E-04 |  | *THSD1* | -1.37 | 0.002178586 |
| *CHN2* | -2.30 | 4.37E-06 |  | *BMERB1* | -1.37 | 3.42E-06 |
| *EDN1* | -2.30 | 2.38E-06 |  | *TOX* | -1.37 | 3.09E-02 |
| *SLC40A1* | -2.30 | 4.78E-09 |  | *NCOA3* | -1.37 | 0.015218057 |
| *LINC00640* | -2.29 | 6.48E-04 |  | *SLC27A1* | -1.36 | 1.57464E-06 |
| *TMEM37* | -2.29 | 2.35E-04 |  | *LRIG3* | -1.36 | 0.015685618 |
| *AK5* | -2.29 | 4.93E-09 |  | *CCDC121* | -1.36 | 0.005949911 |
| *TRPA1* | -2.28 | 3.32E-04 |  | *ANKRD35* | -1.36 | 0.00208686 |
| *MAP1LC3C* | -2.27 | 2.14E-04 |  | *TRPC6* | -1.36 | 0.035466535 |
| *TTPA* | -2.26 | 6.60E-05 |  | *FRMD6-AS1* | -1.36 | 0.028956133 |
| *LOC100507477* | -2.25 | 3.08E-03 |  | *FMN2* | -1.36 | 0.013754754 |
| *RPARP-AS1* | -2.25 | 2.29E-05 |  | *GBP2* | -1.35 | 3.25E-06 |
| *FRMD6* | -2.25 | 4.76E-07 |  | *FAM200B* | -1.35 | 4.49773E-05 |
| *CCL2* | -2.24 | 1.95E-06 |  | *DAAM2* | -1.34 | 0.017164014 |
| *NR0B1* | -2.21 | 2.97E-03 |  | *GLIPR2* | -1.34 | 0.000377614 |
| *MGARP* | -2.21 | 3.82E-08 |  | *SEMA7A* | -1.34 | 0.000297191 |
| *NEXN* | -2.20 | 3.40E-05 |  | *ALPK2* | -1.34 | 0.00727637 |
| *P2RY14* | -2.20 | 2.75E-03 |  | *SOS1-IT1* | -1.34 | 2.33E-02 |
| *ADM2* | -2.19 | 2.98E-08 |  | *NAT8L* | -1.34 | 0.018755613 |
| *HCAR1* | -2.18 | 5.05E-04 |  | *IL20RA* | -1.33 | 0.00437423 |
| *LRRC4C* | -2.18 | 1.56E-03 |  | *NUAK2* | -1.33 | 0.014497216 |
| *ACSS1* | -2.18 | 2.44E-05 |  | *LYPD6B* | -1.33 | 0.040797333 |
| *ARHGAP18* | -2.17 | 2.08E-06 |  | *SYNPO2* | -1.33 | 0.03959495 |
| *INMT* | -2.17 | 1.08E-03 |  | *MID1* | -1.33 | 0.021274123 |
| *RMRP* | -2.16 | 1.83E-03 |  | *RIMS4* | -1.33 | 0.021220931 |
| *MICAL2* | -2.16 | 7.32E-04 |  | *CNN3* | -1.33 | 0.000155532 |
| *DAPK2* | -2.15 | 9.06E-05 |  | *SEMA3G* | -1.33 | 0.026730754 |
| *COQ8A* | -2.15 | 4.64E-11 |  | *SMAGP* | -1.33 | 1.56E-02 |
| *RANBP3L* | -2.15 | 1.97E-03 |  | *ARHGAP10* | -1.33 | 6.86E-04 |
| *VCAM1* | -2.15 | 1.37E-02 |  | *ABTB1* | -1.33 | 0.001532581 |
| *TRIM47* | -2.15 | 5.36E-07 |  | *FAM78A* | -1.32 | 0.002440232 |
| *ARHGAP24* | -2.14 | 8.81E-06 |  | *DES* | -1.32 | 1.25E-02 |
| *GAS6-DT* | -2.13 | 2.15E-05 |  | *SKP2* | -1.32 | 0.009131617 |
| *METTL7B* | -2.13 | 8.19E-07 |  | *VSIG10* | -1.32 | 0.000165452 |
| *LFNG* | -2.13 | 5.94E-11 |  | *SLC9A3R2* | -1.32 | 0.006820887 |
| *RGCC* | -2.12 | 2.50E-04 |  | *TPM1* | -1.32 | 0.010416372 |
| *RNF144B* | -2.12 | 3.65E-05 |  | *LINC01140* | -1.32 | 9.51E-05 |
| *ROBO4* | -2.12 | 1.01E-05 |  | *AHR* | -1.32 | 0.038297301 |
| *PHKG1* | -2.12 | 1.28E-05 |  | *PTPRJ* | -1.32 | 0.004560513 |
| *DUSP14* | -2.12 | 9.44E-12 |  | *MYOZ2* | -1.32 | 0.026709793 |
| *LBH* | -2.12 | 4.76E-07 |  | *IKBKE* | -1.32 | 1.63E-04 |
| *AFF3* | -2.12 | 6.59E-10 |  | *SSBP2* | -1.32 | 2.45095E-06 |
| *WEE1* | -2.12 | 1.47E-06 |  | *PSTPIP2* | -1.31 | 7.63E-05 |
| *IFIT2* | -2.11 | 4.09E-06 |  | *PIK3IP1* | -1.31 | 0.030778978 |
| *LINC01750* | -2.11 | 7.63E-05 |  | *SNX10* | -1.31 | 4.24E-02 |
| *ROR1* | -2.10 | 8.80E-05 |  | *JHY* | -1.31 | 0.000769067 |
| *HCN1* | -2.10 | 3.73E-03 |  | *ELOVL6* | -1.31 | 0.005903647 |
| *CYTOR* | -2.09 | 3.87E-06 |  | *ZNF674* | -1.30 | 0.005794158 |
| *LINC02615* | -2.09 | 1.16E-03 |  | *MEGF9* | -1.30 | 0.004131113 |
| *LINC01013* | -2.07 | 4.11E-03 |  | *ITPKB* | -1.30 | 0.006434482 |
| *MGAM* | -2.07 | 9.92E-04 |  | *STARD5* | -1.30 | 0.007104118 |
| *NRXN2* | -2.06 | 2.68E-06 |  | *FADS3* | -1.30 | 0.000645835 |
| *CYS1* | -2.05 | 1.23E-07 |  | *DUBR* | -1.29 | 0.002804222 |
| *FRY* | -2.05 | 9.79E-05 |  | *MVK* | -1.29 | 0.007900441 |
| *RAB3IL1* | -2.05 | 1.81E-09 |  | *ACVRL1* | -1.29 | 3.87224E-05 |
| *RGMB* | -2.05 | 6.16E-06 |  | *MAP2K3* | -1.29 | 6.92367E-05 |
| *ITGA3* | -2.05 | 2.55924E-07 |  | *MGLL* | -1.29 | 7.68087E-06 |
| *CYP7B1* | -2.03 | 3.21E-03 |  | *AHNAK* | -1.29 | 0.025658482 |
| *VDR* | -2.03 | 1.10E-06 |  | *ZNF575* | -1.29 | 0.016697692 |
| *P2RX7* | -2.02 | 2.91E-04 |  | *TLR3* | -1.28 | 0.006355339 |
| *MSTN* | -2.01 | 4.49E-04 |  | *TMEM170B* | -1.28 | 2.32E-03 |
| *LRRC2* | -2.01 | 9.91E-06 |  | *PER3* | -1.28 | 0.001199714 |
| *STUM* | -2.00 | 7.85E-03 |  | *STAT1* | -1.28 | 0.000991949 |
| *SERTAD4-AS1* | -2.00 | 2.14E-04 |  | *EIF3J-DT* | -1.28 | 0.00125449 |
| *ANKRD33B* | -1.99 | 1.72E-05 |  | *PML* | -1.28 | 0.00150542 |
| *MAP1A* | -1.99 | 5.32E-08 |  | *APPL2* | -1.28 | 0.007027299 |
| *KCNJ2* | -1.99 | 2.09E-04 |  | *SGCG* | -1.28 | 0.007174834 |
| *HTR7P1* | -1.99 | 2.90E-06 |  | *FLI1* | -1.28 | 0.000818275 |
| *RNU4-2* | -1.98 | 1.70E-02 |  | *ERMAP* | -1.27 | 0.000475541 |
| *SQOR* | -1.97 | 4.23178E-09 |  | *IRF1* | -1.27 | 1.26E-03 |
| *ACKR2* | -1.97 | 1.91E-03 |  | *GPR137C* | -1.27 | 0.041612266 |
| *SHROOM3* | -1.96 | 2.93323E-05 |  | *SLC4A4* | -1.27 | 0.040164146 |
| *IQGAP2* | -1.96 | 8.94E-04 |  | *TYRO3* | -1.27 | 7.39E-03 |
| *RIMS1* | -1.96 | 3.87E-05 |  | *FERMT3* | -1.27 | 4.85E-04 |
| *RAB3B* | -1.95 | 5.01E-04 |  | *LPP-AS2* | -1.27 | 0.001473245 |
| *SLC7A14* | -1.95 | 3.67E-04 |  | *RFX8* | -1.27 | 4.44E-02 |
| *ARID5B* | -1.95 | 4.01E-04 |  | *CNKSR2* | -1.26 | 0.043700667 |
| *IFFO1* | -1.94 | 2.19E-08 |  | *CFL2* | -1.26 | 0.001213476 |
| *DCLK2* | -1.94 | 7.80222E-06 |  | *NMI* | -1.26 | 0.000193658 |
| *SERPINB9P1* | -1.94 | 8.67E-03 |  | *CAMK1D* | -1.26 | 0.033592093 |
| *XIRP1* | -1.94 | 5.54E-03 |  | *SLC46A3* | -1.25 | 0.002655456 |
| *LOC100506725* | -1.94 | 3.08E-03 |  | *JAG2* | -1.25 | 0.023125048 |
| *UACA* | -1.93 | 0.000243114 |  | *NEK6* | -1.25 | 2.83537E-06 |
| *BMF* | -1.93 | 0.002067342 |  | *C11orf68* | -1.25 | 3.80569E-07 |
| *RGS16* | -1.93 | 3.27E-03 |  | *NFKBIE* | -1.25 | 0.002287131 |
| *SPP1* | -1.92 | 2.48E-03 |  | *PIWIL4* | -1.25 | 0.004126177 |
| *C8orf34* | -1.92 | 1.70E-03 |  | *FRMD4A* | -1.25 | 0.004154628 |
| *JADE1* | -1.92 | 2.79E-07 |  | *PGRMC2* | -1.24 | 0.000300821 |
| *SNCAIP* | -1.91 | 8.93E-04 |  | *PTCHD4* | -1.24 | 0.032012478 |
| *PAG1* | -1.91 | 7.55E-08 |  | *SWAP70* | -1.24 | 0.004845679 |
| *KRTAP2-3* | -1.91 | 1.93E-02 |  | *C4orf46* | -1.24 | 0.00049967 |
| *CARD10* | -1.91 | 9.38148E-06 |  | *TXNIP* | -1.24 | 0.007766901 |
| *LOC100126784* | -1.91 | 3.01E-04 |  | *OMA1* | -1.23 | 2.25E-04 |
| *DGAT2* | -1.91 | 2.81E-05 |  | *EPHA4* | -1.23 | 0.032012478 |
| *RNU2-2P* | -1.90 | 2.17E-02 |  | *CPNE2* | -1.23 | 1.14E-03 |
| *NR1D1* | -1.90 | 4.70E-08 |  | *MYOM1* | -1.23 | 1.61E-03 |
| *RFLNA* | -1.89 | 5.97E-03 |  | *S100A3* | -1.23 | 0.010324749 |
| *RGMB-AS1* | -1.89 | 1.39E-03 |  | *FAM126A* | -1.23 | 0.010194526 |
| *CITED2* | -1.89 | 6.58E-08 |  | *PDLIM2* | -1.23 | 0.017877223 |
| *RCAN2* | -1.88 | 2.83E-08 |  | *IER5L* | -1.23 | 0.003106336 |
| *GDF5* | -1.88 | 1.46E-02 |  | *UST* | -1.22 | 6.70E-03 |
| *TLR4* | -1.88 | 1.25E-04 |  | *LOC100287837* | -1.22 | 4.24E-02 |
| *FAXDC2* | -1.87 | 4.07E-05 |  | *CTHRC1* | -1.22 | 0.007358473 |
| *SLC29A1* | -1.87 | 8.10E-08 |  | *RIN3* | -1.22 | 0.000749652 |
| *EVI2A* | -1.87 | 8.59E-04 |  | *GSC* | -1.22 | 0.000716249 |
| *GBP3* | -1.87 | 0.00021758 |  | *CMTM3* | -1.22 | 4.05E-04 |
| *INKA1* | -1.86 | 6.78E-04 |  | *CHN1* | -1.21 | 0.042115157 |
| *AMOTL2* | -1.86 | 2.08E-05 |  | *LOC257396* | -1.21 | 0.020704412 |
| *FMO1* | -1.86 | 8.20E-04 |  | *PKD2* | -1.21 | 0.031407494 |
| *PDGFRA* | -1.85 | 5.03E-04 |  | *SLC1A5* | -1.21 | 2.32276E-05 |
| *NHSL2* | -1.85 | 0.026796395 |  | *PPIP5K1* | -1.21 | 0.009765195 |
| *POLH* | -1.84 | 1.15E-07 |  | *LINC01503* | -1.21 | 0.028356479 |
| *SLC38A4* | -1.84 | 1.35E-03 |  | *PSEN2* | -1.21 | 0.000275474 |
| *IFIH1* | -1.83 | 1.35E-05 |  | *AXIN2* | -1.21 | 0.040623057 |
| *PRR16* | -1.83 | 2.22E-05 |  | *YPEL3* | -1.21 | 0.008960308 |
| *TMEM171* | -1.83 | 0.000305704 |  | *CTF1* | -1.21 | 0.000638253 |
| *FGD4* | -1.83 | 7.63E-05 |  | *SNTB2* | -1.20 | 0.001074508 |
| *TDO2* | -1.82 | 7.71E-03 |  | *EPS8L2* | -1.20 | 1.45E-04 |
| *NUDT7* | -1.82 | 2.65E-04 |  | *PYROXD2* | -1.20 | 0.000515175 |
| *PKD1L2* | -1.82 | 5.35609E-05 |  | *FAM131B* | -1.20 | 0.049440902 |
| *IL17RE* | -1.81 | 1.40E-04 |  | *GDPD5* | -1.20 | 5.71E-03 |
| *ZNF367* | -1.81 | 3.46E-04 |  | *ACVR1C* | -1.20 | 0.040855451 |
| *PPFIBP2* | -1.81 | 2.12628E-11 |  | *TRIM59* | -1.19 | 0.035019615 |
| *STOML1* | -1.81 | 4.79079E-07 |  | *LOXL1-AS1* | -1.19 | 0.032207402 |
| *NR3C2* | -1.81 | 0.00018338 |  | *PPP2R5A* | -1.19 | 1.36222E-05 |
| *KCNQ5* | -1.81 | 1.43E-02 |  | *CASP4LP* | -1.19 | 3.81E-02 |
| *HRCT1* | -1.81 | 0.007882351 |  | *RAB40B* | -1.19 | 2.74597E-05 |
| *NR2F1-AS1* | -1.81 | 5.97E-06 |  | *TMEM17* | -1.19 | 1.72E-03 |
| *SAMD3* | -1.81 | 3.18E-02 |  | *MEST* | -1.19 | 0.003187611 |
| *MYCT1* | -1.80 | 2.22E-02 |  | *DPYSL2* | -1.19 | 3.18664E-06 |
| *NRXN3* | -1.79 | 8.88E-03 |  | *LIMD2* | -1.19 | 0.010604678 |
| *RASL11A* | -1.79 | 7.99E-05 |  | *ESR1* | -1.19 | 3.01E-02 |
| *SERTAD4* | -1.79 | 5.03E-04 |  | *ZBED5-AS1* | -1.18 | 0.032568549 |
| *RERG* | -1.79 | 0.009951826 |  | *ZC3H6* | -1.18 | 0.034606616 |
| *ADH1B* | -1.79 | 7.85E-04 |  | *CPA4* | -1.18 | 0.004227115 |
| *TSPAN5* | -1.79 | 1.53E-04 |  | *ZSWIM4* | -1.18 | 0.045269874 |
| *SLC1A3* | -1.79 | 2.11E-04 |  | *SERPINB9* | -1.18 | 0.001394523 |
| *PLEKHG2* | -1.78 | 1.12E-04 |  | *SBF2-AS1* | -1.18 | 1.45E-02 |
| *DDB2* | -1.78 | 5.32279E-05 |  | *IGF2BP2* | -1.18 | 0.014211375 |
| *MSC-AS1* | -1.78 | 7.57E-06 |  | *PRELID3A* | -1.18 | 0.000971158 |
| *KLF2* | -1.78 | 7.91431E-09 |  | *VCL* | -1.18 | 0.030062886 |
| *G0S2* | -1.78 | 1.02E-02 |  | *CCDC89* | -1.18 | 0.023516126 |
| *KLHDC9* | -1.77 | 1.42E-02 |  | *XPC* | -1.18 | 6.22E-03 |
| *DACT3* | -1.77 | 5.94465E-06 |  | *LAYN* | -1.17 | 0.000122239 |
| *TMEM200A* | -1.77 | 0.00047751 |  | *ANO4* | -1.17 | 0.005524973 |
| *PTGER2* | -1.76 | 8.80E-06 |  | *PLCE1* | -1.17 | 0.039461566 |
| *SYNE3* | -1.76 | 0.001176561 |  | *TBCK* | -1.17 | 0.001237254 |
| *GNA14* | -1.76 | 5.23E-05 |  | *SORBS1* | -1.17 | 0.031883798 |
| *OSR2* | -1.75 | 9.73E-09 |  | *CD9* | -1.17 | 0.010628255 |
| *KLHL30* | -1.75 | 1.93E-02 |  | *CORO6* | -1.17 | 0.005643011 |
| *GIPC2* | -1.75 | 7.71E-03 |  | *GNAI1* | -1.16 | 0.001592879 |
| *ST3GAL6* | -1.75 | 0.001813734 |  | *SEMA4F* | -1.16 | 8.19094E-05 |
| *RHOBTB1* | -1.75 | 1.31E-04 |  | *MYO1D* | -1.16 | 0.003974759 |
| *NUAK1* | -1.75 | 9.44E-05 |  | *ATP23* | -1.16 | 0.001402721 |
| *LINC00957* | -1.74 | 1.42E-02 |  | *HIRIP3* | -1.16 | 0.002655456 |
| *DOK5* | -1.74 | 1.47E-06 |  | *SLC22A15* | -1.16 | 0.023695774 |
| *KLHL31* | -1.74 | 0.011465469 |  | *ROBO3* | -1.16 | 0.021049753 |
| *SYT17* | -1.74 | 2.26E-03 |  | *FAM110A* | -1.16 | 0.007589073 |
| *KCNJ16* | -1.74 | 2.60E-02 |  | *HSPB1* | -1.16 | 0.049768676 |
| *PRODH* | -1.74 | 0.021266702 |  | *CREB3L1* | -1.15 | 0.003821251 |
| *POPDC2* | -1.74 | 7.34E-03 |  | *HEBP1* | -1.15 | 0.01981626 |
| *KCNE3* | -1.74 | 5.55E-05 |  | *SLCO3A1* | -1.15 | 0.00655849 |
| *SVIL* | -1.74 | 1.05E-03 |  | *PGBD3* | -1.15 | 0.003003496 |
| *PLCXD3* | -1.74 | 0.012956883 |  | *RP9P* | -1.15 | 0.006744327 |
| *CSF1* | -1.74 | 0.005410661 |  | *AKAP7* | -1.15 | 0.001227676 |
| *CHDH* | -1.73 | 2.21E-03 |  | *TENM4* | -1.15 | 0.037003597 |
| *HMGA1* | -1.73 | 8.36E-05 |  | *TMEM159* | -1.15 | 8.38E-03 |
| *P2RX6* | -1.72 | 1.70E-03 |  | *TPD52L1* | -1.15 | 0.01905072 |
| *FYB1* | -1.72 | 0.008588358 |  | *ACOT1* | -1.15 | 0.024274908 |
| *TMEM132C* | -1.71 | 1.86E-02 |  | *TMOD2* | -1.15 | 0.00952951 |
| *KCNA1* | -1.71 | 0.017166541 |  | *RASAL2-AS1* | -1.15 | 0.049482543 |
| *VEGFC* | -1.71 | 2.30517E-06 |  | *GBP1* | -1.14 | 0.016022908 |
| *REXO5* | -1.70 | 4.3459E-06 |  | *NANOS1* | -1.14 | 0.000373185 |
| *MYO1B* | -1.70 | 3.15E-04 |  | *DHX58* | -1.14 | 0.011465469 |
| *LMO7* | -1.70 | 1.42E-04 |  | *PRXL2C* | -1.14 | 0.013533989 |
| *CAV1* | -1.70 | 1.46685E-10 |  | *CAMK2D* | -1.14 | 0.010868121 |
| *RNF150* | -1.70 | 8.68E-05 |  | *DENND3* | -1.14 | 0.003265791 |
| *DAPK1* | -1.70 | 0.005429816 |  | *GLIS2* | -1.14 | 0.003998403 |
| *ARHGEF28* | -1.70 | 0.001270088 |  | *PNMA8B* | -1.14 | 3.37E-02 |
| *FLT3LG* | -1.69 | 3.18E-04 |  | *TIAM1* | -1.14 | 0.036058341 |
| *C8orf58* | -1.69 | 1.57E-07 |  | *SLC2A12* | -1.14 | 0.027559812 |
| *PLA2G4C* | -1.69 | 3.54E-03 |  | *ARHGEF17* | -1.14 | 0.033775715 |
| *SPEG* | -1.69 | 4.30E-07 |  | *OSR1* | -1.13 | 0.041069014 |
| *LURAP1L* | -1.69 | 1.09064E-08 |  | *CEBPA-DT* | -1.13 | 0.039960693 |
| *ARHGEF3* | -1.68 | 2.06E-03 |  | *ZNF365* | -1.13 | 4.54E-02 |
| *ABCB4* | -1.68 | 0.00609278 |  | *MBNL1* | -1.13 | 0.029202208 |
| *NHLRC4* | -1.67 | 0.003607399 |  | *IER5* | -1.13 | 0.003527621 |
| *ICAM1* | -1.67 | 0.013884768 |  | *C2orf27A* | -1.13 | 0.005524973 |
| *SOD2-OT1* | -1.67 | 1.07E-02 |  | *TMEM187* | -1.13 | 4.83E-02 |
| *SCUBE3* | -1.67 | 3.24E-02 |  | *LMCD1* | -1.13 | 0.017488692 |
| *ACVR2A* | -1.67 | 2.85E-05 |  | *APBB2* | -1.13 | 0.041096869 |
| *GATA6* | -1.67 | 1.38639E-06 |  | *CPED1* | -1.13 | 0.036009959 |
| *LINC01415* | -1.67 | 0.011805898 |  | *TNFRSF1B* | -1.13 | 0.024476804 |
| *NR1D2* | -1.67 | 2.99E-04 |  | *CD27-AS1* | -1.12 | 0.005348164 |
| *DDX60* | -1.66 | 1.24E-03 |  | *MTSS2* | -1.12 | 2.07E-03 |
| *LINGO1* | -1.66 | 0.007951582 |  | *FLJ32255* | -1.12 | 0.005196408 |
| *PRKD1* | -1.66 | 1.16167E-05 |  | *PAQR4* | -1.12 | 0.005543966 |
| *TSC22D1-AS1* | -1.66 | 0.007320291 |  | *RAP2B* | -1.12 | 2.44391E-05 |
| *HNMT* | -1.66 | 4.03095E-06 |  | *ZYX* | -1.12 | 2.83994E-05 |
| *TANC1* | -1.65 | 0.001292233 |  | *TCEANC* | -1.12 | 0.045577433 |
| *LDB3* | -1.65 | 0.034669548 |  | *AJUBA* | -1.12 | 0.012597951 |
| *RMI2* | -1.65 | 2.79E-03 |  | *WDSUB1* | -1.12 | 0.000588235 |
| *ARNT2* | -1.65 | 0.000638253 |  | *CNKSR3* | -1.11 | 1.86E-02 |
| *SLC9A7* | -1.65 | 9.49E-04 |  | *TMEM119* | -1.11 | 0.003539364 |
| *BAALC* | -1.65 | 2.70E-03 |  | *TMEM97* | -1.11 | 9.71442E-05 |
| *SH2D3C* | -1.65 | 0.015004857 |  | *PKP4* | -1.11 | 0.004288975 |
| *GALNT14* | -1.64 | 3.58E-02 |  | *CACNA2D4* | -1.11 | 0.048300495 |
| *TSC22D3* | -1.64 | 1.53E-05 |  | *JAZF1* | -1.11 | 3.33221E-05 |
| *LINC01504* | -1.64 | 0.014772599 |  | *ID2* | -1.11 | 0.005894876 |
| *CDH6* | -1.64 | 5.18E-03 |  | *H1-10-AS1* | -1.11 | 0.006730108 |
| *TGFB2* | -1.64 | 9.75E-06 |  | *ACAN* | -1.10 | 0.049658425 |
| *BDNF-AS* | -1.64 | 0.000324921 |  | *EZH1* | -1.10 | 0.010928979 |
| *CRIM1-DT* | -1.64 | 1.90E-03 |  | *CAVIN2* | -1.10 | 0.003773418 |
| *RNF125* | -1.63 | 0.004865074 |  | *PDP1* | -1.10 | 0.009850756 |
| *ABCD2* | -1.63 | 2.79E-03 |  | *FNBP1* | -1.10 | 0.002245854 |
| *RDH10* | -1.63 | 9.51E-05 |  | *EXT1* | -1.10 | 0.028132936 |
| *RN7SK* | -1.63 | 0.0453971 |  | *PLXDC1* | -1.10 | 0.047999543 |
| *SAMD12* | -1.62 | 0.001424619 |  | *PTGIS* | -1.09 | 0.030644345 |
| *NEK10* | -1.62 | 2.31E-02 |  | *MIR23AHG* | -1.09 | 0.005050049 |
| *LOXL1* | -1.62 | 3.08096E-07 |  | *PPM1H* | -1.09 | 0.017592865 |
| *ARHGAP33* | -1.62 | 1.78E-03 |  | *DTNBP1* | -1.09 | 0.003713451 |
| *LINC01119* | -1.62 | 2.76E-03 |  | *BCL7A* | -1.09 | 0.006167642 |
| *ADRA1D* | -1.62 | 0.016491968 |  | *NMT2* | -1.09 | 0.001626213 |
| *KLF5* | -1.61 | 0.005367455 |  | *TEF* | -1.09 | 0.02257391 |
| *C10orf55* | -1.61 | 0.037686389 |  | *SYPL2* | -1.09 | 0.033123103 |
| *ZNF467* | -1.61 | 0.032158444 |  | *FBXO30-DT* | -1.09 | 0.017592865 |
| *FGF5* | -1.61 | 0.001142588 |  | *PLS3* | -1.08 | 0.007158496 |
| *FHOD1* | -1.61 | 8.40E-07 |  | *ANKRD13A* | -1.08 | 0.009714691 |
| *ZNF704* | -1.61 | 9.77E-06 |  | *RAB29* | -1.08 | 0.000198927 |
| *FGD6* | -1.61 | 4.40E-03 |  | *TNS2* | -1.08 | 0.042362006 |
| *ID4* | -1.60 | 0.003678477 |  | *SPRY1* | -1.08 | 0.00977677 |
| *ARL4C* | -1.60 | 6.16E-05 |  | *VWCE* | -1.08 | 0.032495619 |
| *RRAS2* | -1.60 | 8.46E-08 |  | *TEAD4* | -1.08 | 2.20E-02 |
| *CNN2* | -1.60 | 4.97712E-07 |  | *C5* | -1.08 | 0.049528 |
| *LINC01354* | -1.60 | 0.004174737 |  | *SMAD6* | -1.07 | 0.002677625 |
| *SVIL2P* | -1.60 | 2.43E-02 |  | *WTIP* | -1.07 | 0.000929611 |
| *RPPH1* | -1.60 | 1.24E-02 |  | *PPM1M* | -1.06 | 0.004006325 |
| *ARHGAP26* | -1.60 | 2.61E-04 |  | *IRF1-AS1* | -1.06 | 0.037710249 |
| *AOPEP* | -1.60 | 1.47E-05 |  | *CSPG4* | -1.06 | 0.011008105 |
| *CCDC136* | -1.59 | 3.41794E-08 |  | *IPO5* | -1.06 | 0.012285709 |
| *LINC01569* | -1.59 | 0.01554794 |  | *CTIF* | -1.06 | 0.000547519 |
| *JPH2* | -1.59 | 0.003564473 |  | *LOC646762* | -1.06 | 0.00119377 |
| *TIAM2* | -1.59 | 0.003318327 |  | *SH3BP4* | -1.06 | 0.028433052 |
| *COBLL1* | -1.59 | 3.15E-03 |  | *CBX7* | -1.06 | 0.003107414 |
| *LVRN* | -1.58 | 0.000448551 |  | *AFAP1L1* | -1.06 | 0.035345626 |
| *MID1IP1* | -1.58 | 0.001900015 |  | *FHL3* | -1.06 | 0.00066297 |
| *NEURL1B* | -1.58 | 2.81E-02 |  | *F2RL2* | -1.06 | 0.019650546 |
| *OCLN* | -1.57 | 0.043371668 |  | *RBPJ* | -1.06 | 0.035621357 |
| *FMN1* | -1.57 | 0.013466034 |  | *MINDY4* | -1.06 | 0.022025012 |
| *FGF1* | -1.57 | 1.47E-06 |  | *ULBP1* | -1.05 | 0.046398271 |
| *AQP1* | -1.57 | 0.000723149 |  | *VMAC* | -1.05 | 0.019842226 |
| *PADI2* | -1.57 | 0.026796395 |  | *NLRC5* | -1.05 | 0.035602957 |
| *ADH1A* | -1.56 | 6.24E-03 |  | *DPYD* | -1.05 | 0.017948378 |
| *RFPL4B* | -1.56 | 0.049697288 |  | *PIGV* | -1.04 | 0.000364412 |
| *KCNS3* | -1.56 | 0.024237156 |  | *SLC9A5* | -1.04 | 0.00767924 |
| *MTSS1* | -1.56 | 1.0747E-05 |  | *KLF6* | -1.04 | 0.000391346 |
| *LINC00899* | -1.56 | 0.00066005 |  | *OSBPL10* | -1.04 | 0.028624859 |
| *THSD7B* | -1.56 | 0.018268825 |  | *STAT5A* | -1.04 | 0.002767883 |
| *LIMS2* | -1.56 | 1.84E-05 |  | *LBX2-AS1* | -1.04 | 0.017002946 |
| *WIPF3* | -1.56 | 7.47E-03 |  | *PPP1R13B* | -1.04 | 0.021162642 |
| *ADAMTS3* | -1.56 | 0.030636177 |  | *CCDC146* | -1.04 | 0.020322374 |
| *MAP3K5* | -1.55 | 1.31E-03 |  | *ZMAT3* | -1.04 | 0.001213476 |
| *SLCO4C1* | -1.55 | 0.011740764 |  | *PSMG3-AS1* | -1.04 | 0.014251881 |
| *OLFML2B* | -1.55 | 7.37E-04 |  | *EPS8* | -1.03 | 0.044729118 |
| *OLFM4* | -1.55 | 4.16E-02 |  | *P2RX4* | -1.03 | 1.85E-02 |
| *CHI3L1* | -1.55 | 7.49E-04 |  | *ADAP1* | -1.03 | 0.040086475 |
| *ST6GAL1* | -1.55 | 1.44E-02 |  | *CCDC92* | -1.03 | 0.00727637 |
| *ADORA1* | -1.54 | 1.49E-02 |  | *TP53I11* | -1.03 | 0.005589301 |
| *C14orf180* | -1.54 | 3.55E-02 |  | *GDF6* | -1.03 | 0.041109821 |
| *CCDC170* | -1.54 | 0.000690447 |  | *FAM241A* | -1.03 | 0.001272618 |
| *CCDC15* | -1.54 | 0.007085496 |  | *EMX2OS* | -1.03 | 0.037969858 |
| *MAP7D3* | -1.54 | 9.76745E-06 |  | *CORO1C* | -1.03 | 0.012150346 |
| *LPIN1* | -1.54 | 9.72326E-06 |  | *AFF1-AS1* | -1.03 | 0.022705108 |
| *ARHGAP22* | -1.53 | 0.00028066 |  | *LINC02381* | -1.03 | 0.00307254 |
| *PTH1R* | -1.53 | 0.000886208 |  | *TRAF3IP2-AS1* | -1.02 | 0.010862652 |
| *SPATA9* | -1.53 | 1.64E-02 |  | *TGFB1I1* | -1.02 | 0.004865074 |
| *LOC100240735* | -1.53 | 6.38E-03 |  | *UBE2E2* | -1.02 | 0.000317738 |
| *CALCRL* | -1.53 | 2.58E-04 |  | *CALHM2* | -1.02 | 0.003766744 |
| *TREX1* | -1.53 | 2.61E-04 |  | *MACIR* | -1.02 | 0.002022984 |
| *PRDM16* | -1.53 | 0.000630299 |  | *FAM117A* | -1.02 | 0.011861133 |
| *RPLP0P2* | -1.53 | 2.33E-02 |  | *TRIM22* | -1.02 | 0.011220722 |
| *PEAK1* | -1.52 | 9.10E-03 |  | *CAMK2G* | -1.02 | 0.001972097 |
| *RHEBL1* | -1.52 | 2.11E-02 |  | *OSER1-DT* | -1.01 | 0.014797841 |
| *SUSD3* | -1.52 | 9.85E-03 |  | *REPS2* | -1.01 | 0.020186721 |
| *THBS1* | -1.52 | 0.011803786 |  | *ADHFE1* | -1.01 | 0.004311028 |
| *MAML3* | -1.52 | 0.002322855 |  | *RPL23AP7* | -1.01 | 0.042051681 |
| *TMEM254-AS1* | -1.51 | 2.41E-03 |  | *RIN2* | -1.01 | 0.01670249 |
| *LOC100505942* | -1.51 | 0.048915255 |  | *SCAI* | -1.01 | 0.020704412 |
| *BAMBI* | -1.51 | 9.77E-04 |  | *BCL2L1* | -1.01 | 0.000933318 |
| *TBC1D2* | -1.51 | 0.001018572 |  | *UBA7* | -1.01 | 0.000199561 |
| *NATD1* | -1.51 | 4.05358E-05 |  | *AMN1* | -1.00 | 0.003773171 |
| *LINC00310* | -1.51 | 4.31E-02 |  | *LINC01116* | -1.00 | 0.010316229 |
| *ALS2CL* | -1.51 | 6.66E-03 |  |  |  |  |

**Supplementary Table 5.** Upregulated genes by cAMP in DN adipocytes. Log_2_ fold FC ≥1, Adjective p value <0.05

| **Symbol** | **Log2 Fold Change** | **Adj. P Value** |  | **Symbol** | **Log2 Fold Change** | **Adj. P Value** |
| --- | --- | --- | --- | --- | --- | --- |
| *HAS1* | 7.78 | 4.08E-85 |  | *SLCO5A1* | 1.70 | 2.65E-02 |
| *LINC00473* | 6.70 | 9.61E-53 |  | *MEX3A* | 1.70 | 0.000576396 |
| *RHCG* | 6.43 | 9.65E-36 |  | *SPART-AS1* | 1.70 | 0.017629407 |
| *IL11* | 6.12 | 4.30E-40 |  | *SLC6A2* | 1.70 | 0.028393146 |
| *NR4A1* | 5.92 | 7.20E-86 |  | *GAL* | 1.69 | 0.0100891 |
| *C11orf86* | 5.90 | 1.29E-26 |  | *EXOC3L2* | 1.69 | 0.017183453 |
| *GPR183* | 5.74 | 3.00E-24 |  | *MMP16* | 1.69 | 0.022798499 |
| *FCER1G* | 5.62 | 3.89E-26 |  | *AGAP11* | 1.69 | 0.004233363 |
| *TRH* | 5.62 | 5.65E-28 |  | *SLC17A9* | 1.69 | 0.00013756 |
| *IRF4* | 5.40 | 2.69E-30 |  | *FAM43B* | 1.69 | 6.80E-03 |
| *PTPRN* | 5.37 | 2.12E-36 |  | *RGS2* | 1.68 | 0.001325337 |
| *CTH* | 5.21 | 4.66E-42 |  | *FGF7* | 1.68 | 0.002375387 |
| *SST* | 5.16 | 4.01E-30 |  | *ZDBF2* | 1.68 | 3.26654E-05 |
| *ADAMTS4* | 5.13 | 1.23E-36 |  | *IFI30* | 1.68 | 0.00243957 |
| *SERTM1* | 5.08 | 4.95E-21 |  | *GABARAPL1* | 1.68 | 5.74306E-05 |
| *SH2D2A* | 5.03 | 1.46E-27 |  | *MT2A* | 1.68 | 0.004732332 |
| *LYPD3* | 5.00 | 7.94E-19 |  | *SLC16A4* | 1.67 | 0.000910246 |
| *SIK1* | 5.00 | 7.90E-46 |  | *SCIN* | 1.67 | 0.03445074 |
| *C2CD4A* | 4.96 | 7.29E-22 |  | *TMC7* | 1.67 | 0.003945097 |
| *OASL* | 4.95 | 6.17E-18 |  | *CELA2B* | 1.67 | 0.025917818 |
| *USP2* | 4.94 | 6.57E-21 |  | *GALNT12* | 1.66 | 7.63148E-05 |
| *TAC1* | 4.83 | 1.06E-18 |  | *B3GNT5* | 1.66 | 0.000526482 |
| *GK* | 4.81 | 1.80E-85 |  | *BCL2L11* | 1.66 | 2.19586E-08 |
| *NR4A2* | 4.80 | 5.37E-27 |  | *KLHL29* | 1.65 | 7.07962E-08 |
| *RASD2* | 4.80 | 8.48E-21 |  | *CGA* | 1.65 | 0.047317045 |
| *NBEA* | 4.79 | 1.75E-31 |  | *LIF* | 1.65 | 0.021153916 |
| *BABAM2-AS1* | 4.74 | 4.61E-20 |  | *DYRK3* | 1.65 | 5.28E-12 |
| *AREG* | 4.74 | 4.88E-22 |  | *LINC00887* | 1.65 | 1.55E-02 |
| *SLC6A17* | 4.71 | 5.51E-54 |  | *TGIF1* | 1.64 | 5.91289E-07 |
| *NTRK1* | 4.71 | 9.83E-30 |  | *ADAMTS6* | 1.64 | 0.008847348 |
| *ANXA10* | 4.64 | 2.47E-24 |  | *APBB3* | 1.64 | 3.50E-05 |
| *PKIB* | 4.64 | 8.65E-18 |  | *SLC38A5* | 1.64 | 0.001756056 |
| *SMOX* | 4.64 | 4.29E-49 |  | *ARHGAP32* | 1.63 | 4.78546E-08 |
| *CALCA* | 4.62 | 1.01E-17 |  | *BLK* | 1.63 | 0.046993053 |
| *MT1A* | 4.61 | 1.39E-27 |  | *SDF2L1* | 1.62 | 1.23519E-05 |
| *EREG* | 4.60 | 2.37E-17 |  | *VASH2* | 1.62 | 0.002974913 |
| *C2orf66* | 4.48 | 2.53E-22 |  | *ABTB2* | 1.62 | 5.33E-06 |
| *DIO2* | 4.47 | 1.24E-52 |  | *INO80C* | 1.62 | 1.18736E-09 |
| *INHBA* | 4.46 | 7.18E-28 |  | *PELI1* | 1.62 | 5.98028E-05 |
| *CCR7* | 4.46 | 9.23E-20 |  | *JMJD6* | 1.62 | 2.72359E-08 |
| *C2CD4B* | 4.40 | 2.92E-15 |  | *SNHG15* | 1.62 | 4.56E-07 |
| *MUC13* | 4.38 | 4.20E-16 |  | *C5AR1* | 1.62 | 2.73174E-06 |
| *SCG2* | 4.37 | 1.31E-17 |  | *KCNE5* | 1.61 | 0.004189907 |
| *CACNA1G* | 4.36 | 6.60E-17 |  | *OXSR1* | 1.61 | 1.12E-10 |
| *DUSP4* | 4.25 | 2.75E-17 |  | *LRRC73* | 1.61 | 0.000354202 |
| *ID3* | 4.23 | 2.61E-25 |  | *TDP2* | 1.61 | 4.53E-08 |
| *NPPC* | 4.22 | 1.76E-14 |  | *ITK* | 1.61 | 0.041738055 |
| *FAM167A* | 4.17 | 8.48E-24 |  | *TNFSF4* | 1.61 | 0.004402571 |
| *ADRA2C* | 4.16 | 3.82E-22 |  | *RFK* | 1.61 | 3.19001E-09 |
| *HMX3* | 4.14 | 3.15E-11 |  | *SEMA3A* | 1.61 | 0.000484461 |
| *TUBB2B* | 4.12 | 1.59E-15 |  | *SLC3A2* | 1.61 | 4.91341E-05 |
| *REN* | 4.10 | 2.74E-13 |  | *CYP11A1* | 1.60 | 0.006728978 |
| *PDE4D* | 4.05 | 1.11E-19 |  | *FKBP4* | 1.60 | 2.24802E-08 |
| *FPR2* | 4.05 | 3.52E-11 |  | *FOXC1* | 1.60 | 1.22503E-06 |
| *SDS* | 4.00 | 1.03E-10 |  | *SELENOK* | 1.60 | 2.18757E-05 |
| *KCNG1* | 3.98 | 9.06E-39 |  | *RSAD2* | 1.60 | 0.018542717 |
| *SULT4A1* | 3.98 | 2.35E-11 |  | *CACNB2* | 1.60 | 0.006279246 |
| *APOLD1* | 3.95 | 1.08E-20 |  | *HMOX1* | 1.60 | 8.56055E-05 |
| *LINC00664* | 3.90 | 5.68E-20 |  | *HYOU1* | 1.59 | 3.45692E-05 |
| *KLHL13* | 3.89 | 5.99E-20 |  | *SELENOKP1* | 1.59 | 0.003163487 |
| *WNT1* | 3.88 | 8.85E-10 |  | *BTBD11* | 1.59 | 3.90E-02 |
| *CCDC177* | 3.88 | 2.04E-11 |  | *CARD8-AS1* | 1.59 | 5.40898E-05 |
| *MLLT11* | 3.87 | 2.64E-28 |  | *CFAP69* | 1.59 | 0.000319808 |
| *NR4A3* | 3.87 | 3.75E-18 |  | *SAMD11* | 1.59 | 0.008235373 |
| *ACTBL2* | 3.87 | 9.81E-11 |  | *RAB21* | 1.58 | 3.69E-13 |
| *CCK* | 3.87 | 8.25E-11 |  | *TMEM233* | 1.58 | 0.010714416 |
| *FLRT3* | 3.84 | 7.40E-14 |  | *METRNL* | 1.58 | 5.31846E-12 |
| *GPRC5A* | 3.82 | 2.33E-17 |  | *ATOH7* | 1.58 | 0.035729726 |
| *PITPNC1* | 3.81 | 1.24E-52 |  | *HOMER1* | 1.57 | 8.61815E-05 |
| *LINC00322* | 3.77 | 1.34E-09 |  | *SIAH1* | 1.57 | 0.000738523 |
| *CRISPLD2* | 3.77 | 4.51E-49 |  | *SIX2* | 1.57 | 0.001370558 |
| *P4HA3* | 3.76 | 1.37E-14 |  | *HAS3* | 1.56 | 7.34E-03 |
| *ECEL1* | 3.73 | 2.87E-13 |  | *ADGRG2* | 1.56 | 1.56E-03 |
| *PGAP1* | 3.69 | 8.19E-25 |  | *PRPS2* | 1.56 | 1.49016E-07 |
| *SHISA2* | 3.67 | 9.59E-12 |  | *MYO7A* | 1.56 | 2.53E-03 |
| *PCSK1* | 3.67 | 2.88E-24 |  | *RDH8* | 1.56 | 0.003752262 |
| *SLC2A13* | 3.64 | 4.37E-21 |  | *DAB2IP* | 1.56 | 2.46966E-06 |
| *ARFGEF3* | 3.63 | 6.12E-11 |  | *ALPK3* | 1.56 | 0.003113274 |
| *DNAH17* | 3.61 | 1.19E-14 |  | *LIPG* | 1.56 | 0.008403139 |
| *TMEM100* | 3.59 | 2.34E-21 |  | *ADAM28* | 1.56 | 0.049141767 |
| *TMOD1* | 3.57 | 2.20E-16 |  | *DDIT3* | 1.55 | 0.00040056 |
| *CCL20* | 3.55 | 1.22E-17 |  | *SGSM1* | 1.55 | 0.038480238 |
| *CASP9* | 3.50 | 2.57E-14 |  | *SLC7A8* | 1.55 | 4.54735E-08 |
| *C11orf96* | 3.49 | 3.55E-28 |  | *KDM7A-DT* | 1.55 | 6.04791E-07 |
| *SYT12* | 3.48 | 8.17E-22 |  | *ISG15* | 1.55 | 6.35576E-05 |
| *TUBA4A* | 3.48 | 1.01E-15 |  | *ID4* | 1.55 | 0.039199987 |
| *PDE4B* | 3.46 | 1.69E-13 |  | *FNDC11* | 1.55 | 0.03415051 |
| *LYVE1* | 3.40 | 4.80E-12 |  | *TTYH2* | 1.55 | 0.001153396 |
| *PGM2L1* | 3.39 | 3.69E-14 |  | *JUND* | 1.55 | 0.000147022 |
| *LOC100507403* | 3.36 | 8.41E-08 |  | *UBASH3B* | 1.55 | 9.42E-06 |
| *ISG20* | 3.34 | 1.68E-13 |  | *FICD* | 1.55 | 8.80909E-09 |
| *PAX1* | 3.34 | 1.17E-06 |  | *NUDT9P1* | 1.54 | 0.016264185 |
| *PDE3A* | 3.34 | 6.59E-12 |  | *ECE2* | 1.54 | 1.29E-05 |
| *MIR614* | 3.30 | 4.06E-11 |  | *UBE2QL1* | 1.54 | 4.88E-02 |
| *KLHL15* | 3.29 | 9.38E-32 |  | *WNK3* | 1.54 | 0.007880776 |
| *SYNDIG1* | 3.28 | 8.65E-17 |  | *QPCTL* | 1.53 | 1.1212E-05 |
| *HES4* | 3.28 | 2.83E-10 |  | *MIR635* | 1.53 | 0.000303183 |
| *CHMP1B* | 3.23 | 6.57E-44 |  | *HSPA5* | 1.53 | 4.41121E-05 |
| *CHRDL2* | 3.22 | 3.77E-10 |  | *EDNRB* | 1.53 | 5.79E-03 |
| *MMP12* | 3.19 | 3.01E-10 |  | *CYCS* | 1.52 | 6.26E-04 |
| *GAP43* | 3.17 | 5.23E-09 |  | *ZDHHC23* | 1.52 | 0.001137971 |
| *KCNH1* | 3.17 | 3.20E-08 |  | *SLC6A15* | 1.52 | 0.000764884 |
| *CYP2S1* | 3.16 | 3.88E-09 |  | *KDM6B* | 1.52 | 1.65126E-13 |
| *AJAP1* | 3.15 | 2.19E-07 |  | *SLC25A19* | 1.51 | 2.62E-03 |
| *GADD45G* | 3.15 | 3.94E-08 |  | *INSRR* | 1.51 | 0.038259131 |
| *CREM* | 3.14 | 2.78E-21 |  | *C9orf131* | 1.51 | 0.049469094 |
| *RNF152* | 3.13 | 1.37E-14 |  | *CHD7* | 1.51 | 2.7555E-05 |
| *ITPRIP* | 3.12 | 5.79E-25 |  | *MEG9* | 1.51 | 0.014053725 |
| *LINC00313* | 3.12 | 4.25E-06 |  | *ABCC4* | 1.51 | 0.000731949 |
| *FIBCD1* | 3.12 | 1.03E-10 |  | *LXN* | 1.51 | 0.001617858 |
| *PRR5L* | 3.11 | 5.74E-15 |  | *DUSP5* | 1.51 | 0.004296244 |
| *CST2* | 3.09 | 2.35E-12 |  | *SYN3* | 1.50 | 0.016737966 |
| *FAM87B* | 3.05 | 6.66E-13 |  | *SOWAHD* | 1.50 | 0.033027129 |
| *HEYL* | 3.05 | 1.39E-11 |  | *SNORA24* | 1.50 | 0.034195359 |
| *PMEPA1* | 3.04 | 1.38E-36 |  | *CYGB* | 1.50 | 5.11E-06 |
| *PDE4C* | 3.04 | 1.59E-06 |  | *ENOSF1* | 1.50 | 4.18115E-06 |
| *ATF3* | 3.02 | 2.42E-10 |  | *PPP1R15B* | 1.50 | 3.7065E-06 |
| *LINC02274* | 3.02 | 2.84E-06 |  | *TLE3* | 1.50 | 1.60E-06 |
| *ZNF331* | 3.01 | 2.78E-12 |  | *UAP1* | 1.50 | 7.49751E-08 |
| *INHBA-AS1* | 3.01 | 2.45E-08 |  | *USP2-AS1* | 1.50 | 0.042466667 |
| *BDNF* | 2.99 | 1.14E-13 |  | *DERL3* | 1.49 | 0.043687752 |
| *NECTIN1* | 2.97 | 4.64E-35 |  | *LYPD1* | 1.49 | 0.049469094 |
| *AADACP1* | 2.96 | 1.13E-05 |  | *CHST15* | 1.49 | 1.48711E-05 |
| *CHGB* | 2.96 | 1.02E-07 |  | *ARL5B* | 1.49 | 9.92E-06 |
| *PHYHIP* | 2.94 | 4.41E-12 |  | *CDA* | 1.49 | 0.0191888 |
| *MSI1* | 2.94 | 5.34E-06 |  | *TOMM40* | 1.48 | 7.93E-05 |
| *SDIM1* | 2.93 | 5.58E-07 |  | *ZBTB21* | 1.48 | 1.59404E-06 |
| *TNFRSF18* | 2.92 | 8.06E-07 |  | *RIPOR2* | 1.48 | 0.005055726 |
| *FSTL3* | 2.92 | 6.19E-12 |  | *AGPAT4-IT1* | 1.48 | 0.020893108 |
| *CAMK2N2* | 2.91 | 9.37E-07 |  | *TMUB1* | 1.48 | 6.45E-03 |
| *C17orf58* | 2.90 | 3.19E-15 |  | *BCAS4* | 1.48 | 0.000732157 |
| *RAMP1* | 2.89 | 8.65E-17 |  | *KL* | 1.47 | 0.017776319 |
| *IGFN1* | 2.89 | 1.41E-04 |  | *FAM131A* | 1.47 | 7.22846E-07 |
| *PTP4A1* | 2.88 | 3.43E-35 |  | *PCDH12* | 1.47 | 0.007306358 |
| *KRT75* | 2.86 | 1.02E-07 |  | *PLOD2* | 1.47 | 0.008598832 |
| *RCSD1* | 2.85 | 4.31E-06 |  | *KPNA2* | 1.47 | 1.83E-06 |
| *SPOCK2* | 2.84 | 8.81E-09 |  | *EGR2* | 1.46 | 0.020488574 |
| *C8A* | 2.84 | 1.41E-05 |  | *BACH2* | 1.46 | 0.00350481 |
| *TLNRD1* | 2.83 | 4.40E-32 |  | *ZSWIM5* | 1.46 | 0.000480196 |
| *CXCR4* | 2.82 | 1.30E-06 |  | *C2orf88* | 1.46 | 1.06E-07 |
| *CD55* | 2.82 | 8.42E-15 |  | *TSKU* | 1.46 | 0.014492757 |
| *RNF122* | 2.82 | 8.74E-17 |  | *ARRDC3* | 1.46 | 0.003786306 |
| *KCNK5* | 2.80 | 1.11E-07 |  | *METTL1* | 1.46 | 7.21003E-08 |
| *SNTG2* | 2.80 | 2.67E-07 |  | *SULT1C2* | 1.45 | 0.030225928 |
| *FOS* | 2.79 | 1.4956E-09 |  | *TMEM88* | 1.45 | 0.028237321 |
| *HAS2* | 2.77 | 2.35E-09 |  | *LINC00941* | 1.45 | 0.006765478 |
| *FOSB* | 2.77 | 4.95E-07 |  | *FN3K* | 1.44 | 0.020296951 |
| *PDK4* | 2.77 | 2.14E-07 |  | *ATG101* | 1.44 | 2.31211E-06 |
| *SNAI1* | 2.74 | 6.05E-13 |  | *GFOD1* | 1.44 | 0.003546572 |
| *MFSD2A* | 2.73 | 6.43E-07 |  | *TMEM158* | 1.44 | 0.004915536 |
| *THAP2* | 2.72 | 9.71E-13 |  | *EPB41L3* | 1.44 | 0.000249319 |
| *ATP1B3* | 2.71 | 2.23E-18 |  | *DLL1* | 1.44 | 0.002037212 |
| *CASC15* | 2.71 | 5.66E-07 |  | *SLC22A4* | 1.44 | 0.001757092 |
| *BMP6* | 2.71 | 6.87E-08 |  | *TNFAIP6* | 1.44 | 0.011784108 |
| *MAP3K14-AS1* | 2.70 | 5.38E-07 |  | *TENT5C* | 1.43 | 0.011618081 |
| *CYFIP2* | 2.69 | 2.42E-07 |  | *CASP7* | 1.43 | 3.79982E-07 |
| *KSR1* | 2.69 | 4.28E-12 |  | *ZNF462* | 1.43 | 0.000254062 |
| *DDIT4* | 2.68 | 9.37E-15 |  | *MEDAG* | 1.43 | 1.91585E-06 |
| *DPYSL3* | 2.68 | 2.85E-12 |  | *ADAMTS9-AS2* | 1.43 | 0.020525765 |
| *BMP8A* | 2.67 | 2.2544E-09 |  | *FKBP11* | 1.43 | 1.6318E-05 |
| *DLK1* | 2.67 | 3.2657E-05 |  | *AMMECR1* | 1.43 | 0.009139465 |
| *HEY2* | 2.66 | 2.67E-05 |  | *DNAJA1* | 1.42 | 2.66063E-07 |
| *KCNE4* | 2.66 | 3.84E-09 |  | *SPTLC3* | 1.42 | 0.000426008 |
| *SH3TC1* | 2.66 | 7.88E-11 |  | *SDE2* | 1.42 | 4.34E-11 |
| *IL1RN* | 2.63 | 6.00E-04 |  | *EAF2* | 1.42 | 0.003786306 |
| *TMEM217* | 2.63 | 2.51E-10 |  | *CDR2L* | 1.42 | 3.52851E-09 |
| *FGF10* | 2.60 | 2.60E-04 |  | *KCNJ8* | 1.42 | 0.004391075 |
| *ETNPPL* | 2.60 | 0.00033938 |  | *KLKB1* | 1.42 | 0.045746845 |
| *FOXQ1* | 2.59 | 1.20E-03 |  | *HSPA8* | 1.42 | 4.48828E-07 |
| *OAS1* | 2.59 | 1.64E-08 |  | *DCHS1* | 1.42 | 0.007679852 |
| *TREM1* | 2.59 | 2.22E-04 |  | *SBSN* | 1.41 | 0.003083764 |
| *HBEGF* | 2.59 | 5.9479E-07 |  | *PDE2A* | 1.41 | 0.038535773 |
| *CMSS1* | 2.57 | 1.016E-12 |  | *PSD4* | 1.41 | 0.015760261 |
| *MELTF* | 2.56 | 2.42E-12 |  | *CFAP58* | 1.41 | 0.010551463 |
| *DNAJC12* | 2.55 | 8.02E-09 |  | *SDSL* | 1.41 | 2.27E-02 |
| *STC1* | 2.55 | 1.46E-03 |  | *FRMPD4* | 1.41 | 0.005282765 |
| *B4GALT1* | 2.55 | 4.93E-20 |  | *HERC4* | 1.40 | 0.002093798 |
| *INSYN2A* | 2.55 | 2.11E-07 |  | *STAR* | 1.40 | 0.03195408 |
| *RHBDF2* | 2.55 | 8.02E-10 |  | *DNAJC27* | 1.40 | 0.000738728 |
| *NRG1* | 2.55 | 2.74E-06 |  | *NEFM* | 1.40 | 0.049544172 |
| *KCNA10* | 2.55 | 0.00014372 |  | *CKS2* | 1.40 | 4.59E-03 |
| *C10orf90* | 2.54 | 1.35E-06 |  | *WIPI1* | 1.40 | 7.30412E-10 |
| *PGBD5* | 2.53 | 4.27E-07 |  | *LARP4* | 1.40 | 6.25175E-09 |
| *MSX1* | 2.53 | 3.72E-20 |  | *SLC33A1* | 1.40 | 6.6178E-06 |
| *PPARGC1A* | 2.53 | 3.46E-06 |  | *PGM3* | 1.39 | 6.61E-07 |
| *TGM2* | 2.53 | 1.93E-05 |  | *TREH* | 1.39 | 0.037374001 |
| *TUBB3* | 2.52 | 8.42E-15 |  | *XBP1* | 1.39 | 1.87165E-06 |
| *TUBB2A* | 2.52 | 5.45E-08 |  | *EIF2AK3* | 1.38 | 0.000103782 |
| *RELT* | 2.52 | 1.82E-09 |  | *LINC00622* | 1.37 | 0.035537712 |
| *TMEM151A* | 2.52 | 2.82E-06 |  | *NCR3LG1* | 1.37 | 1.36E-06 |
| *CHST1* | 2.50 | 1.25E-05 |  | *FABP5* | 1.37 | 0.002696217 |
| *BTG3* | 2.50 | 3.47E-07 |  | *IL1RL2* | 1.37 | 0.000343101 |
| *ATP2A3* | 2.50 | 8.28E-08 |  | *HSPH1* | 1.37 | 0.000195596 |
| *GRIA3* | 2.50 | 2.24E-07 |  | *FZD4* | 1.36 | 0.001320853 |
| *S1PR1* | 2.50 | 6.7811E-09 |  | *MAPKAPK2* | 1.36 | 2.09475E-05 |
| *LETM2* | 2.50 | 1.15E-06 |  | *ST6GALNAC2* | 1.36 | 0.003747165 |
| *LRFN4* | 2.49 | 3.76E-21 |  | *BHLHE40* | 1.36 | 1.64E-04 |
| *FGF2* | 2.49 | 4.65E-10 |  | *CRLF1* | 1.36 | 0.014942309 |
| *TFPI2* | 2.49 | 7.74E-05 |  | *EMD* | 1.36 | 0.00010594 |
| *C5AR2* | 2.49 | 1.72E-04 |  | *LDLRAD4* | 1.35 | 0.007539255 |
| *GPAT3* | 2.47 | 3.5586E-07 |  | *NCS1* | 1.35 | 4.80851E-10 |
| *RAB20* | 2.47 | 3.67E-12 |  | *SH3PXD2A* | 1.35 | 2.17473E-05 |
| *DAW1* | 2.47 | 1.24E-07 |  | *JARID2* | 1.35 | 0.006781989 |
| *LONRF3* | 2.46 | 1.26E-08 |  | *PAX9* | 1.35 | 0.022832996 |
| *PDE10A* | 2.46 | 3.25E-08 |  | *KALRN* | 1.35 | 0.000550049 |
| *KRT16* | 2.46 | 1.2375E-05 |  | *NAA50* | 1.34 | 1.66265E-09 |
| *CBARP* | 2.44 | 1.03E-07 |  | *ZC3H12A* | 1.34 | 4.10201E-05 |
| *LMLN2* | 2.43 | 2.08E-05 |  | *BLOC1S3* | 1.33 | 3.29123E-05 |
| *LBP* | 2.42 | 1.24E-05 |  | *GREM1* | 1.33 | 0.00373159 |
| *HUNK* | 2.42 | 1.222E-06 |  | *RNF113A* | 1.33 | 9.46326E-07 |
| *PCSK1N* | 2.42 | 3.66E-04 |  | *EIF4A3* | 1.32 | 3.72281E-08 |
| *PTPN5* | 2.41 | 5.88E-04 |  | *CDV3* | 1.32 | 1.21879E-05 |
| *CLCF1* | 2.41 | 2.1216E-09 |  | *UPF3B* | 1.32 | 0.002870593 |
| *DUSP2* | 2.41 | 3.1275E-05 |  | *AGPS* | 1.32 | 0.00075704 |
| *PNPLA5* | 2.41 | 0.00078544 |  | *GJC2* | 1.32 | 0.007021073 |
| *ZBTB32* | 2.41 | 1.32E-04 |  | *MOSPD1* | 1.32 | 0.002790854 |
| *SPAG4* | 2.40 | 8.5316E-07 |  | *ESYT2* | 1.31 | 6.07847E-06 |
| *TMEM71* | 2.40 | 8.66E-09 |  | *KAZN* | 1.31 | 1.1212E-05 |
| *PER1* | 2.39 | 5.35E-19 |  | *KCNK6* | 1.31 | 0.004915536 |
| *SEMA6A* | 2.39 | 4.07E-06 |  | *TWIST1* | 1.31 | 0.002365673 |
| *DOC2B* | 2.38 | 7.06E-04 |  | *UBE2S* | 1.31 | 1.46213E-06 |
| *RHOB* | 2.37 | 3.62E-12 |  | *LINC01347* | 1.31 | 0.013252681 |
| *FAM183BP* | 2.37 | 8.17E-04 |  | *HYAL3* | 1.31 | 0.003451516 |
| *FAM122C* | 2.37 | 7.70E-06 |  | *IL4R* | 1.30 | 0.003066916 |
| *GPR161* | 2.37 | 1.3756E-09 |  | *STX11* | 1.30 | 0.014734491 |
| *GPR3* | 2.36 | 1.96E-04 |  | *MTHFD1L* | 1.30 | 0.00051224 |
| *WFDC21P* | 2.36 | 2.59E-04 |  | *SLC22A23* | 1.30 | 7.26E-06 |
| *JUNB* | 2.35 | 1.33E-11 |  | *RAPGEF1* | 1.29 | 5.18093E-05 |
| *PHLDB2* | 2.35 | 5.30E-12 |  | *TMEM106A* | 1.29 | 0.001184507 |
| *PRL* | 2.35 | 0.00068846 |  | *DIRAS1* | 1.29 | 0.038344391 |
| *IL1A* | 2.34 | 4.48E-05 |  | *STK32B* | 1.29 | 0.007444854 |
| *CFAP58-DT* | 2.34 | 0.00052731 |  | *POLB* | 1.29 | 0.00881808 |
| *CHSY1* | 2.34 | 1.79E-11 |  | *SLC2A1* | 1.29 | 0.022718147 |
| *KLHDC7B* | 2.33 | 8.90E-06 |  | *MSX2* | 1.29 | 0.005038146 |
| *GNAO1* | 2.33 | 2.077E-05 |  | *HSPE1* | 1.29 | 7.85774E-05 |
| *CFAP74* | 2.32 | 0.0016235 |  | *CISH* | 1.28 | 0.002542989 |
| *ZNF184* | 2.31 | 1.25E-04 |  | *PPME1* | 1.28 | 3.23359E-06 |
| *ENPEP* | 2.31 | 1.302E-06 |  | *SELENOS* | 1.28 | 4.74789E-09 |
| *PTGS2* | 2.30 | 5.84E-04 |  | *RELL1* | 1.28 | 0.000739298 |
| *PIGA* | 2.30 | 9.22E-11 |  | *PFKP* | 1.27 | 0.000785153 |
| *AADAC* | 2.29 | 1.58E-04 |  | *KDM7A* | 1.27 | 0.00601872 |
| *BMP8B* | 2.28 | 6.62E-06 |  | *SPTY2D1* | 1.27 | 4.85541E-05 |
| *PITPNM1* | 2.28 | 4.961E-21 |  | *CRY2* | 1.27 | 0.015722169 |
| *JPH1* | 2.27 | 7.97E-04 |  | *ELL2* | 1.27 | 0.000506079 |
| *ZFP92* | 2.27 | 4.47E-04 |  | *MANF* | 1.26 | 0.001932932 |
| *FXYD6* | 2.27 | 9.76E-04 |  | *RIPK2* | 1.26 | 1.18567E-05 |
| *SV2C* | 2.26 | 1.62E-03 |  | *KIRREL3* | 1.26 | 0.02289763 |
| *ADAMTS9* | 2.26 | 3.05E-09 |  | *HSPD1* | 1.26 | 2.00978E-07 |
| *SLC16A6* | 2.25 | 0.00043421 |  | *PKDCC* | 1.26 | 0.000361194 |
| *DEFB1* | 2.25 | 2.28E-03 |  | *USP36* | 1.26 | 0.000323506 |
| *DBX2* | 2.24 | 1.82E-03 |  | *FAM107B* | 1.26 | 0.000984358 |
| *CD14* | 2.24 | 7.63E-10 |  | *SLC31A1* | 1.26 | 8.32725E-05 |
| *PAPPA* | 2.23 | 5.3663E-05 |  | *CCT2* | 1.25 | 1.28219E-06 |
| *SEPTIN6* | 2.22 | 3.11E-08 |  | *ZPR1* | 1.25 | 0.000127276 |
| *IGFBP1* | 2.22 | 7.79E-03 |  | *CDC7* | 1.24 | 0.014307322 |
| *FHOD3* | 2.22 | 8.66E-08 |  | *NOP16* | 1.24 | 0.000769223 |
| *SLC7A5* | 2.21 | 9.67E-06 |  | *SEC23B* | 1.24 | 3.10E-06 |
| *SLC6A12* | 2.21 | 0.00124731 |  | *HSPA7* | 1.24 | 0.04372309 |
| *KCTD20* | 2.21 | 2.93E-13 |  | *LRRC59* | 1.24 | 3.79009E-06 |
| *NKAIN1* | 2.21 | 4.1974E-06 |  | *SDC1* | 1.24 | 0.016191245 |
| *AMIGO2* | 2.20 | 5.1468E-09 |  | *ATAD3B* | 1.24 | 0.000166443 |
| *LINC00673* | 2.20 | 3.8205E-05 |  | *NKRF* | 1.24 | 0.000164858 |
| *SPHK1* | 2.18 | 4.24E-14 |  | *GAB2* | 1.23 | 5.728E-05 |
| *CSGALNACT1* | 2.17 | 2.87E-05 |  | *IVNS1ABP* | 1.23 | 5.04196E-06 |
| *UCN* | 2.17 | 0.00157755 |  | *GNL3* | 1.23 | 4.5815E-05 |
| *IL1B* | 2.17 | 1.75E-04 |  | *AP1S3* | 1.23 | 0.006824017 |
| *DUSP8* | 2.17 | 4.6874E-05 |  | *MT1E* | 1.23 | 0.010810631 |
| *UCP1* | 2.17 | 0.00655959 |  | *ATP2A2* | 1.22 | 3.36741E-06 |
| *THBD* | 2.17 | 1.24E-03 |  | *VASN* | 1.22 | 2.08435E-05 |
| *SLAMF9* | 2.17 | 6.22E-04 |  | *POLR3D* | 1.22 | 1.0188E-07 |
| *MT1L* | 2.16 | 6.37E-06 |  | *DDX21* | 1.22 | 0.00450945 |
| *CHAC1* | 2.16 | 2.10E-04 |  | *AKAP13* | 1.22 | 0.016232511 |
| *DIRAS3* | 2.16 | 1.2301E-05 |  | *GPCPD1* | 1.22 | 0.000311582 |
| *SPINK1* | 2.16 | 3.49E-04 |  | *PTHLH* | 1.22 | 0.016713636 |
| *PIP5K1B* | 2.16 | 0.00024208 |  | *BZW2* | 1.22 | 3.41572E-06 |
| *C6orf223* | 2.15 | 0.00352868 |  | *GJA1* | 1.21 | 0.034346884 |
| *GFPT2* | 2.15 | 2.718E-09 |  | *KLF4* | 1.21 | 0.002848654 |
| *OR1F1* | 2.15 | 5.06E-03 |  | *MFHAS1* | 1.21 | 3.26698E-06 |
| *CSRNP1* | 2.14 | 2.01E-12 |  | *MEIS1* | 1.21 | 0.020720718 |
| *SOCS3* | 2.14 | 5.10E-13 |  | *PPTC7* | 1.21 | 1.38567E-06 |
| *STAC2* | 2.14 | 2.48E-05 |  | *ZNF697* | 1.21 | 0.000206954 |
| *RDH12* | 2.14 | 0.0005471 |  | *NPIPB4* | 1.21 | 0.030101853 |
| *C9orf72* | 2.13 | 3.2631E-08 |  | *PAX8* | 1.21 | 0.004187886 |
| *FLVCR2* | 2.13 | 2.54E-05 |  | *WDR55* | 1.20 | 0.000454417 |
| *PAQR5* | 2.13 | 2.77E-04 |  | *CUL4B* | 1.20 | 0.00893866 |
| *HES1* | 2.13 | 2.4245E-16 |  | *ATAD3A* | 1.20 | 2.56442E-05 |
| *DUSP1* | 2.13 | 6.8354E-09 |  | *TMEM39A* | 1.20 | 1.56964E-05 |
| *RASL11B* | 2.13 | 0.00026908 |  | *CCDC130* | 1.20 | 4.38548E-07 |
| *FLT1* | 2.13 | 0.00065119 |  | *RARA* | 1.20 | 0.000122064 |
| *KLHL26* | 2.13 | 8.28E-08 |  | *NUS1* | 1.19 | 0.000286897 |
| *SUCNR1* | 2.13 | 0.00185832 |  | *RPL22L1* | 1.19 | 4.13744E-05 |
| *GADD45B* | 2.12 | 3.72E-06 |  | *PYCR1* | 1.19 | 0.00213755 |
| *FAM124A* | 2.10 | 1.754E-08 |  | *GPRC5D-AS1* | 1.19 | 0.043671218 |
| *SLC8A2* | 2.10 | 6.02E-03 |  | *RABGGTB* | 1.19 | 9.03236E-06 |
| *CXCL5* | 2.09 | 1.32E-02 |  | *SEC61G* | 1.19 | 0.000430195 |
| *MKRN9P* | 2.09 | 0.00052583 |  | *NOCT* | 1.19 | 0.026384975 |
| *MAPK4* | 2.08 | 6.36E-03 |  | *SOX4* | 1.19 | 0.00081286 |
| *B3GNT2* | 2.08 | 4.64E-11 |  | *DOCK8* | 1.18 | 0.006419227 |
| *RAB3A* | 2.08 | 0.00147421 |  | *ARRDC2* | 1.18 | 0.011236551 |
| *IGFBP3* | 2.08 | 9.07E-04 |  | *EPOP* | 1.18 | 0.006924578 |
| *PPDPFL* | 2.08 | 0.00426088 |  | *TUBB6* | 1.18 | 0.000504337 |
| *CPEB4* | 2.08 | 6.34E-14 |  | *COQ10B* | 1.18 | 6.66447E-06 |
| *AADACL2* | 2.08 | 1.33E-03 |  | *CCNH* | 1.18 | 0.000711086 |
| *GPR157* | 2.08 | 2.1735E-05 |  | *DNAJC1* | 1.18 | 0.00020426 |
| *HAL* | 2.07 | 0.00045516 |  | *MT1X* | 1.17 | 0.012267449 |
| *PPP1R3C* | 2.07 | 4.74E-05 |  | *PICALM* | 1.17 | 1.91471E-05 |
| *SGIP1* | 2.06 | 0.00074121 |  | *TAF13* | 1.17 | 2.43637E-07 |
| *LAMC3* | 2.06 | 1.50E-04 |  | *PHLDA1* | 1.17 | 0.028702337 |
| *ACE2* | 2.05 | 0.00449003 |  | *HSP90B3P* | 1.17 | 0.000519536 |
| *C1QL2* | 2.05 | 0.00544436 |  | *KLF16* | 1.17 | 5.87701E-05 |
| *CDC6* | 2.05 | 3.6483E-07 |  | *GUK1* | 1.16 | 0.008948624 |
| *NFATC2* | 2.05 | 0.00016663 |  | *PPP3CA* | 1.16 | 0.016657106 |
| *PPP1R14C* | 2.04 | 0.0004614 |  | *EIF4A1* | 1.16 | 5.26736E-06 |
| *FZD8* | 2.04 | 7.6975E-06 |  | *SLC7A1* | 1.16 | 0.001108109 |
| *GPRC5D* | 2.04 | 2.06E-03 |  | *ANKH* | 1.16 | 0.006208855 |
| *C17orf107* | 2.04 | 4.71E-03 |  | *ARFIP2* | 1.16 | 2.30023E-06 |
| *SV2B* | 2.04 | 3.94E-03 |  | *SYAP1* | 1.16 | 0.000652565 |
| *QRFP* | 2.03 | 0.00068464 |  | *ESPL1* | 1.16 | 0.016432376 |
| *NXN* | 2.03 | 1.32E-03 |  | *SIN3B* | 1.16 | 5.0153E-07 |
| *LCE2C* | 2.03 | 1.11E-02 |  | *ABLIM1* | 1.16 | 0.020775798 |
| *IL6* | 2.03 | 0.00069648 |  | *SLC52A2* | 1.16 | 8.31371E-05 |
| *APLN* | 2.03 | 0.00171099 |  | *CMIP* | 1.15 | 1.65829E-05 |
| *NECTIN4* | 2.02 | 7.85E-04 |  | *JOSD1* | 1.15 | 1.10958E-06 |
| *ELOVL7* | 2.01 | 0.00049946 |  | *RBM15* | 1.15 | 0.005586579 |
| *BDKRB1* | 2.01 | 1.27E-05 |  | *GNG7* | 1.14 | 0.043671218 |
| *SP6* | 2.01 | 2.44E-04 |  | *PAICS* | 1.14 | 0.003616189 |
| *HSPA6* | 2.01 | 1.65E-05 |  | *EAF1* | 1.14 | 1.11054E-06 |
| *ADGRV1* | 2.01 | 0.00505573 |  | *ZFC3H1* | 1.14 | 0.005215778 |
| *SPON1* | 2.01 | 9.4992E-05 |  | *RRP9* | 1.14 | 0.000919273 |
| *TP53AIP1* | 2.00 | 0.00622962 |  | *ZNF460* | 1.14 | 0.017183453 |
| *SLC1A2* | 2.00 | 3.6927E-05 |  | *CCND2* | 1.14 | 0.028013625 |
| *KBTBD8* | 2.00 | 1.30E-08 |  | *ZSCAN5A* | 1.13 | 0.000502859 |
| *LONRF2* | 1.99 | 3.214E-05 |  | *HSPA13* | 1.13 | 0.000693683 |
| *GALNT9* | 1.98 | 0.01147052 |  | *NDEL1* | 1.13 | 0.00137088 |
| *APBA1* | 1.98 | 1.15E-05 |  | *LINC01128* | 1.13 | 0.001713754 |
| *MT1G* | 1.98 | 0.00138627 |  | *CHST3* | 1.13 | 8.58235E-05 |
| *SLC19A2* | 1.98 | 8.2542E-10 |  | *MORF4L2* | 1.13 | 3.90868E-05 |
| *GLA* | 1.98 | 4.63E-15 |  | *STIP1* | 1.13 | 0.00010594 |
| *SH3PXD2B* | 1.97 | 4.1781E-13 |  | *ST3GAL4* | 1.13 | 0.015633056 |
| *FOXF1* | 1.96 | 0.00024317 |  | *SELENOI* | 1.13 | 5.59427E-06 |
| *CACNA1H* | 1.96 | 4.16E-03 |  | *EVA1C* | 1.13 | 0.027453613 |
| *MMP10* | 1.96 | 0.00326209 |  | *RORA* | 1.13 | 0.014376447 |
| *HYAL1* | 1.96 | 5.7109E-08 |  | *PTRH2* | 1.13 | 0.00024342 |
| *MESD* | 1.96 | 4.1295E-16 |  | *PELO* | 1.13 | 8.98383E-05 |
| *RSPO3* | 1.96 | 6.6178E-06 |  | *CSRP1* | 1.12 | 0.000325109 |
| *CDH24* | 1.95 | 1.30E-08 |  | *EPOR* | 1.12 | 0.001250457 |
| *SPSB4* | 1.95 | 0.0068573 |  | *CBX4* | 1.12 | 0.00864867 |
| *AGPAT4* | 1.95 | 1.06E-04 |  | *PTPN1* | 1.12 | 0.000362018 |
| *PTGES* | 1.94 | 8.83E-12 |  | *PHF10* | 1.11 | 0.001697316 |
| *AKAP6* | 1.94 | 0.00018765 |  | *PNO1* | 1.11 | 9.60744E-05 |
| *CREB5* | 1.94 | 2.06E-04 |  | *DPH2* | 1.11 | 7.05134E-05 |
| *RFLNB* | 1.94 | 9.6312E-10 |  | *ICOSLG* | 1.11 | 0.017170638 |
| *CTNNB1* | 1.93 | 2.68E-14 |  | *ARFGAP3* | 1.11 | 0.0001566 |
| *SFRP4* | 1.93 | 4.54E-04 |  | *PLAT* | 1.11 | 0.006471402 |
| *IGLON5* | 1.93 | 3.62E-03 |  | *NAT9* | 1.11 | 2.17473E-05 |
| *RASD1* | 1.93 | 5.22E-09 |  | *SERP1* | 1.11 | 7.74071E-06 |
| *AP5B1* | 1.92 | 6.36E-05 |  | *RRP12* | 1.11 | 8.29219E-06 |
| *FOSL2* | 1.92 | 2.84E-14 |  | *CBLB* | 1.11 | 0.015218542 |
| *MEG8* | 1.92 | 0.0003279 |  | *TIPARP* | 1.11 | 0.011236551 |
| *DNAJB9* | 1.92 | 1.8083E-09 |  | *FARSA* | 1.10 | 0.000288394 |
| *LINC01011* | 1.92 | 2.69E-02 |  | *ITGA5* | 1.10 | 0.013560672 |
| *ENTPD7* | 1.91 | 5.4384E-07 |  | *WHRN* | 1.10 | 0.022417544 |
| *WARS1* | 1.91 | 6.1234E-12 |  | *GOSR2* | 1.10 | 2.0324E-05 |
| *PI15* | 1.91 | 0.01359597 |  | *ARMCX2* | 1.10 | 0.002616712 |
| *MOB1B* | 1.90 | 1.00E-09 |  | *PFDN2* | 1.10 | 0.000487833 |
| *SLCO4A1* | 1.90 | 1.35E-04 |  | *PNPLA8* | 1.10 | 0.00036203 |
| *ST3GAL5* | 1.90 | 3.96E-07 |  | *APBB1IP* | 1.10 | 0.001054515 |
| *GALNT2* | 1.90 | 1.0311E-10 |  | *ELOVL4* | 1.10 | 0.027908129 |
| *GEM* | 1.89 | 1.27E-07 |  | *SFXN2* | 1.10 | 0.034866907 |
| *C12orf60* | 1.89 | 1.47E-04 |  | *ATP13A3* | 1.10 | 0.022321678 |
| *HTRA3* | 1.89 | 2.32E-06 |  | *AIF1L* | 1.09 | 0.002554175 |
| *IMPA2* | 1.89 | 2.67E-06 |  | *USP35* | 1.09 | 0.002375387 |
| *LGALS9* | 1.89 | 0.00114609 |  | *IPO4* | 1.09 | 0.000622209 |
| *CHD1* | 1.88 | 6.7221E-05 |  | *TIMM8A* | 1.09 | 0.015443122 |
| *BMP2* | 1.88 | 5.06E-03 |  | *TCEAL9* | 1.09 | 0.007569437 |
| *WNT10B* | 1.87 | 0.00132534 |  | *LONRF1* | 1.09 | 0.000676578 |
| *CEBPE* | 1.87 | 1.87E-02 |  | *EIF4E* | 1.09 | 0.000681504 |
| *FGFR1* | 1.87 | 2.7427E-09 |  | *HERPUD1* | 1.09 | 0.006094195 |
| *RCE1* | 1.86 | 2.1161E-14 |  | *LINC01137* | 1.09 | 0.008385946 |
| *PID1* | 1.86 | 9.32E-05 |  | *SSTR1* | 1.08 | 0.020625985 |
| *SLC25A25* | 1.85 | 1.35E-06 |  | *BAG3* | 1.08 | 0.000471879 |
| *TCIM* | 1.85 | 1.31E-02 |  | *HIC2* | 1.08 | 0.000702341 |
| *RTL9* | 1.85 | 4.50E-05 |  | *TVP23B* | 1.08 | 0.000119062 |
| *CACNA1G-AS1* | 1.85 | 0.01692417 |  | *RADIL* | 1.08 | 0.000484686 |
| *WNT5A* | 1.84 | 3.9087E-05 |  | *SESN2* | 1.08 | 0.010477785 |
| *PNP* | 1.84 | 3.1566E-06 |  | *NOLC1* | 1.08 | 0.001515446 |
| *ESYT3* | 1.84 | 0.00167957 |  | *TXNDC11* | 1.07 | 2.91494E-05 |
| *ACKR3* | 1.84 | 1.0405E-10 |  | *MAPK13* | 1.07 | 0.006279246 |
| *SLC26A9* | 1.84 | 0.01673945 |  | *GTPBP4* | 1.07 | 7.51912E-05 |
| *CHL1* | 1.84 | 8.38E-03 |  | *DLGAP1-AS1* | 1.07 | 0.002334399 |
| *KLRD1* | 1.84 | 9.06E-04 |  | *LINC01605* | 1.07 | 0.028237321 |
| *CYP19A1* | 1.84 | 8.63E-03 |  | *PALMD* | 1.07 | 0.048394068 |
| *XYLT1* | 1.84 | 9.58E-13 |  | *USPL1* | 1.07 | 0.017839215 |
| *PRKAR1A* | 1.84 | 5.28E-12 |  | *BYSL* | 1.07 | 0.000695576 |
| *QRICH2* | 1.84 | 0.00048446 |  | *CLMP* | 1.07 | 0.011943866 |
| *CREB3L2* | 1.84 | 2.80E-06 |  | *ALKBH5* | 1.07 | 0.000587511 |
| *ST3GAL1* | 1.83 | 2.1247E-09 |  | *GRPEL1* | 1.07 | 0.004758655 |
| *CYSLTR2* | 1.83 | 0.00408442 |  | *HCCS* | 1.07 | 0.003831342 |
| *INSYN2B* | 1.82 | 0.0028086 |  | *FASTKD1* | 1.06 | 2.44077E-05 |
| *TYMP* | 1.82 | 1.0481E-06 |  | *PPAN* | 1.06 | 0.009731564 |
| *PAPPA2* | 1.82 | 7.71E-03 |  | *SLC25A32* | 1.06 | 5.7605E-05 |
| *MEFV* | 1.82 | 2.75E-02 |  | *RNF126* | 1.06 | 0.000354949 |
| *IGFBP4* | 1.82 | 2.068E-10 |  | *BDKRB2* | 1.06 | 0.044355997 |
| *CASZ1* | 1.81 | 1.28E-02 |  | *ABL1* | 1.06 | 0.000187714 |
| *DLX3* | 1.81 | 0.00097298 |  | *TUBA1C* | 1.06 | 0.002525886 |
| *GMPPB* | 1.80 | 1.0801E-08 |  | *ALDH18A1* | 1.06 | 0.001298858 |
| *NAMPT* | 1.80 | 4.7342E-10 |  | *TMED5* | 1.06 | 0.001065599 |
| *OLR1* | 1.80 | 9.70E-03 |  | *THBS2* | 1.05 | 0.023028438 |
| *ZNF703* | 1.79 | 2.53E-08 |  | *DKK1* | 1.05 | 0.016924167 |
| *MRPS30-DT* | 1.79 | 0.00344049 |  | *PDSS1* | 1.05 | 0.01222922 |
| *HS3ST3A1* | 1.79 | 0.01687313 |  | *KIF21A* | 1.05 | 0.00247406 |
| *ODC1* | 1.79 | 2.23E-08 |  | *BAG1* | 1.05 | 0.012878773 |
| *ACTL8* | 1.79 | 0.00846501 |  | *MYO10* | 1.05 | 0.044908475 |
| *CST1* | 1.78 | 0.04451026 |  | *ZNF672* | 1.04 | 0.005665813 |
| *RRN3* | 1.78 | 5.9451E-08 |  | *IBTK* | 1.04 | 0.001098296 |
| *KCNK1* | 1.78 | 0.00068658 |  | *USP37* | 1.04 | 0.007539255 |
| *PLCZ1* | 1.78 | 1.07E-02 |  | *GOLT1B* | 1.04 | 0.000259338 |
| *ASPG* | 1.78 | 2.99E-02 |  | *COLEC12* | 1.04 | 0.010612827 |
| *TSPYL2* | 1.78 | 6.6056E-07 |  | *FOXC2* | 1.04 | 0.03742387 |
| *ACSL4* | 1.78 | 2.6155E-08 |  | *RRN3P2* | 1.04 | 0.049017069 |
| *DUSP15* | 1.78 | 0.02984289 |  | *ZNF10* | 1.04 | 0.029138053 |
| *KIF19* | 1.78 | 0.03729327 |  | *DNAJA4* | 1.04 | 0.000262011 |
| *PKNOX2* | 1.77 | 8.9564E-06 |  | *LYSMD3* | 1.04 | 0.011931322 |
| *NAP1L5* | 1.77 | 3.2007E-08 |  | *URB2* | 1.04 | 0.000174544 |
| *VEGFA* | 1.77 | 1.4664E-08 |  | *PRELID3B* | 1.04 | 0.000878624 |
| *ANKRD53* | 1.76 | 2.93E-04 |  | *KCTD5* | 1.03 | 0.000928992 |
| *OXCT2* | 1.76 | 0.02249976 |  | *MAB21L1* | 1.03 | 0.018009923 |
| *GRAMD1C* | 1.76 | 1.26E-03 |  | *DDX39A* | 1.03 | 4.22576E-06 |
| *NCALD* | 1.76 | 8.93E-07 |  | *TACC2* | 1.03 | 0.02792594 |
| *SOCS2* | 1.76 | 2.1016E-06 |  | *UPP1* | 1.03 | 0.036053871 |
| *MT1M* | 1.76 | 1.02E-04 |  | *PHLDB3* | 1.03 | 0.020525765 |
| *CRY1* | 1.76 | 3.6247E-09 |  | *COP1* | 1.03 | 0.00543116 |
| *SH3BP5* | 1.76 | 1.45E-06 |  | *RANBP2* | 1.03 | 0.019786734 |
| *PLCB1* | 1.75 | 7.0981E-05 |  | *CHORDC1* | 1.03 | 0.013651273 |
| *PLA2G2A* | 1.74 | 0.00034364 |  | *PBX1* | 1.02 | 0.003831342 |
| *GDNF* | 1.74 | 0.00526468 |  | *NUP50-DT* | 1.02 | 0.003113274 |
| *AVPI1* | 1.74 | 0.00078515 |  | *IDH3A* | 1.02 | 0.001261616 |
| *WNT9A* | 1.74 | 3.6896E-07 |  | *NOP2* | 1.02 | 0.002504338 |
| *TARID* | 1.74 | 0.01499061 |  | *MYD88* | 1.02 | 0.000108488 |
| *SGK1* | 1.74 | 9.672E-07 |  | *CPT1A* | 1.02 | 0.036827124 |
| *FMNL1* | 1.73 | 6.213E-06 |  | *PRPF4* | 1.02 | 1.27014E-05 |
| *PRMT9* | 1.73 | 1.37E-04 |  | *ARFGAP1* | 1.02 | 3.94211E-05 |
| *TRABD2B* | 1.73 | 4.69E-05 |  | *ARAP3* | 1.02 | 0.005963062 |
| *SIK3* | 1.73 | 2.3654E-13 |  | *ITPR1* | 1.02 | 0.038814089 |
| *TGM5* | 1.73 | 0.00175709 |  | *CDC37* | 1.02 | 0.001020279 |
| *PDLIM4* | 1.73 | 1.95E-06 |  | *FAM210A* | 1.01 | 7.94352E-05 |
| *SLC2A3* | 1.73 | 9.8164E-05 |  | *GAS1* | 1.01 | 0.049845156 |
| *DDT* | 1.73 | 0.03424145 |  | *SRPRB* | 1.01 | 0.00024622 |
| *HOXA5* | 1.73 | 0.00024432 |  | *AHSA1* | 1.01 | 0.000198551 |
| *GMNN* | 1.73 | 3.18E-06 |  | *CCNJ* | 1.01 | 0.000654924 |
| *CDK2AP2* | 1.72 | 6.78E-09 |  | *LIMS1* | 1.01 | 0.000816988 |
| *GOT1* | 1.72 | 0.00048312 |  | *ZNF222* | 1.01 | 0.003968875 |
| *TRAF4* | 1.72 | 9.95E-08 |  | *ZRANB1* | 1.01 | 0.006507707 |
| *HPDL* | 1.72 | 2.30E-04 |  | *SNHG16* | 1.01 | 7.50462E-05 |
| *YRDC* | 1.72 | 2.5533E-07 |  | *GCAT* | 1.01 | 0.006456904 |
| *BHLHA15* | 1.71 | 0.03740386 |  | *CENPN* | 1.01 | 0.005194355 |
| *NFIL3* | 1.71 | 5.6037E-11 |  | *KCNJ14* | 1.00 | 0.03814517 |
| *TFRC* | 1.71 | 3.61E-10 |  | *QSOX2* | 1.00 | 0.023577329 |
| *MYOM3* | 1.71 | 0.00135863 |  | *RRN3P1* | 1.00 | 0.048641106 |
| *OVOL1* | 1.71 | 3.52E-02 |  | *EIF5A* | 1.00 | 0.001303939 |
| *PCSK9* | 1.70 | 0.00034358 |  | *PLEKHB2* | 1.00 | 0.000847385 |
| *MB21D2* | 1.70 | 5.4079E-08 |  | *TDG* | 1.00 | 0.001635678 |
| *KYNU* | 1.70 | 0.00683201 |  | *PHETA2* | 1.00 | 0.013927318 |
| *SPTBN5* | 1.70 | 1.67E-03 |  |  |  |  |

**Supplementary Table 6.** Downregulated genes by cAMP in DN adipocytes. Log_2_ FC ≤-1, Adjective P value <0.05.

| **Symbol** | **Log2 Fold Change** | **Adj. P Value** |  | **Symbol** | **Log2 Fold Change** | **Adj. P Value** |
| --- | --- | --- | --- | --- | --- | --- |
| *PTX3* | -4.86 | 7.49E-32 |  | *LINGO1* | -1.46 | 0.023726977 |
| *ANKRD1* | -4.27 | 3.84E-29 |  | *AQP1* | -1.46 | 0.03848021 |
| *RPSAP52* | -4.05 | 2.88E-19 |  | *STOML1* | -1.46 | 1.69E-03 |
| *PPL* | -4.02 | 1.83E-42 |  | *TMEM178B* | -1.46 | 0.046643004 |
| *ST8SIA1* | -4.01 | 2.52E-18 |  | *RNF125* | -1.45 | 0.039693856 |
| *SERPINE1* | -3.83 | 6.57E-16 |  | *FMN1* | -1.45 | 0.00780588 |
| *MEOX2* | -3.81 | 1.43E-18 |  | *LINC00957* | -1.44 | 0.021223593 |
| *BMP4* | -3.75 | 1.20E-29 |  | *DACT3* | -1.44 | 2.20E-04 |
| *ADAMTS8* | -3.63 | 6.00E-13 |  | *IL17RE* | -1.44 | 1.23E-03 |
| *KIT* | -3.63 | 7.08E-14 |  | *LVRN* | -1.44 | 0.008785193 |
| *HMGA2-AS1* | -3.54 | 1.03E-10 |  | *OSER1-DT* | -1.44 | 8.68291E-05 |
| *KRTAP1-5* | -3.46 | 1.70E-09 |  | *NR2F2* | -1.44 | 4.61545E-07 |
| *HECW2* | -3.42 | 8.58E-12 |  | *SLC12A8* | -1.43 | 0.006390512 |
| *TENT5B* | -3.35 | 1.04E-12 |  | *TNS3* | -1.43 | 0.010839492 |
| *VGLL3* | -3.34 | 8.74E-17 |  | *TLR3* | -1.43 | 0.002750872 |
| *CCN2* | -3.33 | 2.10E-09 |  | *MYOZ2* | -1.43 | 0.017527399 |
| *DEPTOR* | -3.19 | 4.09E-09 |  | *DCLK2* | -1.43 | 1.56E-05 |
| *CX3CL1* | -3.16 | 4.41E-12 |  | *SLC9A7* | -1.43 | 0.001759896 |
| *ATP8B1* | -3.14 | 2.11E-19 |  | *VMAC* | -1.43 | 1.79E-03 |
| *ANKRD33B* | -3.07 | 1.96E-11 |  | *COBLL1* | -1.43 | 4.33E-03 |
| *SEC14L5* | -3.03 | 4.58E-08 |  | *PIK3R1* | -1.43 | 0.001146763 |
| *MAP3K7CL* | -3.02 | 2.73E-30 |  | *MARCHF4* | -1.42 | 2.19E-02 |
| *CCL2* | -2.95 | 3.72E-08 |  | *MAP1LC3C* | -1.42 | 0.0054357 |
| *KCNS2* | -2.95 | 6.08E-10 |  | *CDC25B* | -1.42 | 1.35E-06 |
| *RGMB-AS1* | -2.92 | 8.90E-09 |  | *USP44* | -1.42 | 0.00213755 |
| *RXFP1* | -2.92 | 2.43E-07 |  | *PNMA8B* | -1.41 | 0.00081286 |
| *KCNJ2* | -2.84 | 2.52E-09 |  | *TNFAIP2* | -1.41 | 0.000377648 |
| *CCN1* | -2.83 | 2.57E-14 |  | *ETS1* | -1.41 | 0.009376052 |
| *ELOVL2* | -2.78 | 9.03E-06 |  | *MYO1D* | -1.41 | 0.000140103 |
| *ITGA3* | -2.78 | 8.04E-11 |  | *CAMK1D* | -1.41 | 2.7681E-06 |
| *FMO2* | -2.76 | 1.31E-05 |  | *DNMT3B* | -1.41 | 0.006857296 |
| *XIRP1* | -2.71 | 1.43E-06 |  | *HEPACAM* | -1.41 | 0.004915536 |
| *P2RX7* | -2.68 | 7.62E-09 |  | *DUSP14* | -1.40 | 1.52E-05 |
| *LMOD1* | -2.68 | 1.56E-10 |  | *LINC00310* | -1.40 | 2.18E-02 |
| *LOXL4* | -2.67 | 7.41E-39 |  | *VEGFC* | -1.40 | 0.000615677 |
| *NR1D1* | -2.66 | 5.29E-19 |  | *RAC2* | -1.40 | 0.014508027 |
| *SCUBE3* | -2.65 | 7.41E-08 |  | *GAS1RR* | -1.40 | 5.68E-03 |
| *PRR15* | -2.64 | 9.38E-08 |  | *KLLN* | -1.40 | 0.015475475 |
| *CDC42EP3* | -2.64 | 1.92E-10 |  | *CMAHP* | -1.40 | 0.009345326 |
| *IFIT1* | -2.64 | 1.43E-08 |  | *NUDT7* | -1.40 | 0.00017059 |
| *CDH5* | -2.60 | 1.70E-05 |  | *GAS6-AS1* | -1.40 | 2.00E-02 |
| *DAPK2* | -2.60 | 8.62E-09 |  | *FYB1* | -1.40 | 3.89E-03 |
| *RTL3* | -2.59 | 7.06E-05 |  | *ZEB1* | -1.39 | 0.010128551 |
| *MAFB* | -2.57 | 3.41E-13 |  | *LFNG* | -1.39 | 1.38E-02 |
| *TRHDE-AS1* | -2.56 | 3.80E-07 |  | *NCAPG* | -1.39 | 2.46E-02 |
| *IRAG1* | -2.54 | 8.64E-12 |  | *AP3M2* | -1.39 | 7.05134E-05 |
| *ACKR4* | -2.54 | 8.02E-09 |  | *RIMS3* | -1.38 | 0.038002944 |
| *IL16* | -2.54 | 2.85E-09 |  | *CEBPA* | -1.38 | 0.036764299 |
| *MYCT1* | -2.53 | 1.59E-06 |  | *CD200* | -1.38 | 1.47E-02 |
| *EVA1A* | -2.53 | 3.12E-11 |  | *TM4SF18* | -1.38 | 0.04438892 |
| *PIK3IP1* | -2.53 | 2.69E-08 |  | *RIPOR3* | -1.38 | 4.64E-03 |
| *INMT* | -2.52 | 2.68E-06 |  | *TRIM47* | -1.38 | 9.2343E-06 |
| *IL34* | -2.49 | 5.27E-08 |  | *CCDC170* | -1.38 | 0.002824174 |
| *RGMB* | -2.49 | 9.49E-07 |  | *BRIP1* | -1.37 | 0.010519324 |
| *HDAC9* | -2.48 | 1.35E-06 |  | *HMMR* | -1.37 | 4.83E-02 |
| *IFIT3* | -2.46 | 1.03E-07 |  | *GEMIN2P2* | -1.36 | 7.51912E-05 |
| *IFIT2* | -2.46 | 1.66E-09 |  | *JAG2* | -1.36 | 0.019568522 |
| *EDN1* | -2.45 | 1.55E-05 |  | *ACVRL1* | -1.36 | 1.5638E-06 |
| *PTGIR* | -2.43 | 3.06E-05 |  | *NEMP1* | -1.36 | 5.18093E-05 |
| *HMGA2* | -2.42 | 5.37E-05 |  | *ANGPTL2* | -1.36 | 0.000543393 |
| *ADRA2A* | -2.42 | 2.15E-11 |  | *LINC00607* | -1.36 | 0.041837241 |
| *MALL* | -2.42 | 8.44E-06 |  | *SREBF1* | -1.36 | 0.00893866 |
| *VCAM1* | -2.41 | 3.33E-03 |  | *LYPLAL1* | -1.36 | 0.003934639 |
| *NAV3* | -2.41 | 2.15E-06 |  | *ZNF438* | -1.35 | 1.00E-03 |
| *RERG* | -2.41 | 4.86E-05 |  | *PRKD1* | -1.35 | 0.000587389 |
| *RGS4* | -2.41 | 9.32E-06 |  | *PLA2G4C* | -1.35 | 0.016755662 |
| *FAXDC2* | -2.39 | 2.44E-08 |  | *ATP1A1-AS1* | -1.35 | 0.010614599 |
| *C10orf55* | -2.37 | 6.21E-06 |  | *DES* | -1.35 | 0.025751887 |
| *ROBO4* | -2.37 | 7.23E-07 |  | *RCAN2* | -1.35 | 9.06848E-05 |
| *RGS3* | -2.37 | 4.71E-16 |  | *PTCHD4* | -1.35 | 0.029774163 |
| *PLAU* | -2.36 | 1.40E-06 |  | *KIAA0754* | -1.35 | 0.020488574 |
| *SHROOM3* | -2.36 | 1.04E-10 |  | *LNX1* | -1.35 | 0.006390512 |
| *CHI3L1* | -2.35 | 1.56E-04 |  | *ST6GAL1* | -1.34 | 0.048259514 |
| *NRXN2* | -2.33 | 9.84E-08 |  | *CCDC191* | -1.34 | 0.002428139 |
| *C1orf198* | -2.33 | 9.07E-14 |  | *RTKN2* | -1.34 | 0.047181092 |
| *AK5* | -2.33 | 1.19E-09 |  | *AKAP3* | -1.34 | 0.013232512 |
| *SLC7A14* | -2.33 | 1.07E-10 |  | *PTPN3* | -1.34 | 0.003507535 |
| *RMI2* | -2.32 | 5.48E-05 |  | *LINC01094* | -1.34 | 0.011943866 |
| *CNTN3* | -2.32 | 5.76E-04 |  | *KCNN4* | -1.34 | 0.046442892 |
| *TRIB2* | -2.32 | 3.12E-06 |  | *NLRC5* | -1.34 | 9.71E-06 |
| *NR0B1* | -2.31 | 2.45E-05 |  | *LOC100507642* | -1.34 | 1.91E-02 |
| *RGS7* | -2.30 | 7.32E-05 |  | *RFX2* | -1.33 | 3.55859E-07 |
| *BBC3* | -2.29 | 2.45E-07 |  | *PDE1C* | -1.33 | 0.009339845 |
| *CYTH3* | -2.29 | 5.21E-14 |  | *CARD10* | -1.33 | 2.75E-03 |
| *DDAH1* | -2.29 | 8.25E-10 |  | *KCNB1* | -1.33 | 0.044627755 |
| *RAB3B* | -2.27 | 9.01E-06 |  | *HIP1R* | -1.33 | 0.001451692 |
| *IFIH1* | -2.27 | 3.64E-11 |  | *LOXL1-AS1* | -1.33 | 0.004207731 |
| *DAB2* | -2.27 | 4.01E-11 |  | *PRDM16* | -1.33 | 0.005426426 |
| *SCNN1A* | -2.26 | 9.49E-04 |  | *AXIN2* | -1.32 | 0.005495737 |
| *SNCAIP* | -2.26 | 3.05E-04 |  | *ROBO3* | -1.32 | 5.35E-03 |
| *CYTOR* | -2.25 | 1.41E-09 |  | *INA* | -1.32 | 0.041486165 |
| *DBP* | -2.25 | 3.18E-05 |  | *XAF1* | -1.32 | 0.049676186 |
| *BMF* | -2.25 | 5.15E-05 |  | *ELOVL6* | -1.32 | 0.003611432 |
| *ZNF367* | -2.24 | 7.16E-07 |  | *CEBPA-DT* | -1.32 | 2.45E-02 |
| *LINC01085* | -2.24 | 1.02E-07 |  | *LOC100288570* | -1.32 | 0.02177474 |
| *CACNB4* | -2.23 | 4.47E-06 |  | *CORO6* | -1.32 | 0.000194011 |
| *CNGA3* | -2.22 | 5.87E-04 |  | *NEK6* | -1.32 | 1.72E-04 |
| *IFFO1* | -2.22 | 8.18E-15 |  | *PLXNC1* | -1.32 | 0.017385325 |
| *ARHGEF3* | -2.22 | 2.58E-10 |  | *MLPH* | -1.32 | 3.67E-02 |
| *THBS1* | -2.21 | 6.89E-05 |  | *FBLIM1* | -1.32 | 0.000138118 |
| *COQ8A* | -2.21 | 8.38E-07 |  | *LPIN1* | -1.32 | 0.000234999 |
| *RUNX1T1* | -2.20 | 3.91E-05 |  | *NCOA3* | -1.31 | 0.010887305 |
| *CST6* | -2.20 | 2.60E-03 |  | *FMO4* | -1.31 | 0.006555077 |
| *CPA4* | -2.20 | 8.68E-08 |  | *EXT1* | -1.31 | 0.000451127 |
| *SLC40A1* | -2.20 | 2.91E-05 |  | *LPP-AS2* | -1.31 | 0.000897818 |
| *STING1* | -2.20 | 4.47E-11 |  | *CENPE* | -1.31 | 0.045351327 |
| *CCDC81* | -2.18 | 7.05E-05 |  | *ZNF519* | -1.31 | 0.002762946 |
| *GIMAP2* | -2.18 | 2.78E-03 |  | *TRIM6* | -1.31 | 0.020081505 |
| *IL17RD* | -2.18 | 3.51E-12 |  | *STMN3* | -1.30 | 0.000767014 |
| *VDR* | -2.18 | 1.31E-08 |  | *LINC00933* | -1.30 | 0.036773545 |
| *KRT18* | -2.17 | 4.31E-06 |  | *MCM5* | -1.30 | 0.000436046 |
| *NHSL2* | -2.17 | 7.32E-05 |  | *FLI1* | -1.30 | 0.004590967 |
| *ANK3* | -2.17 | 6.00E-04 |  | *CD9* | -1.30 | 6.6761E-05 |
| *ARHGAP18* | -2.15 | 7.26E-07 |  | *TNFRSF14* | -1.30 | 1.32E-05 |
| *PSCA* | -2.14 | 7.04E-04 |  | *PIK3CD* | -1.30 | 0.012002902 |
| *SLC6A4* | -2.14 | 1.04E-03 |  | *MYO7B* | -1.30 | 0.002784262 |
| *FAM111B* | -2.13 | 6.02E-03 |  | *OSR1* | -1.30 | 0.016737966 |
| *PHKG1* | -2.13 | 2.44E-04 |  | *ARRB1* | -1.30 | 1.44E-04 |
| *HCAR1* | -2.13 | 3.67E-04 |  | *DOK5* | -1.30 | 8.59527E-05 |
| *USP53* | -2.13 | 1.65E-09 |  | *ETV1* | -1.29 | 0.004315149 |
| *GPAM* | -2.12 | 2.67E-05 |  | *MPP4* | -1.29 | 0.020655481 |
| *CYP7B1* | -2.12 | 1.38E-05 |  | *RHOBTB1* | -1.29 | 0.010118247 |
| *MICAL2* | -2.11 | 7.60E-06 |  | *MTSS1* | -1.29 | 0.000699197 |
| *PLA2G5* | -2.11 | 3.03E-05 |  | *AHNAK2* | -1.29 | 3.29E-02 |
| *NEK2* | -2.11 | 3.91E-03 |  | *RPARP-AS1* | -1.29 | 1.72E-02 |
| *ARNT2* | -2.10 | 2.95E-04 |  | *SH2D5* | -1.29 | 0.034934484 |
| *LAMA3* | -2.09 | 2.69E-03 |  | *ADAMTS12* | -1.29 | 6.66E-06 |
| *MAML3* | -2.09 | 2.60E-06 |  | *BTBD8* | -1.29 | 0.036941499 |
| *GBP3* | -2.09 | 8.25E-05 |  | *ADORA1* | -1.28 | 0.021087839 |
| *CCDC9B* | -2.09 | 1.39E-08 |  | *DDX60L* | -1.28 | 0.021704015 |
| *LIPH* | -2.08 | 2.18E-03 |  | *CELSR2* | -1.28 | 0.005928453 |
| *TRIL* | -2.08 | 8.01E-05 |  | *LRRN4CL* | -1.28 | 0.000507578 |
| *PDE5A* | -2.07 | 7.02E-06 |  | *LRRC75A* | -1.28 | 0.025469898 |
| *NPAS1* | -2.06 | 3.41E-04 |  | *C1orf220* | -1.28 | 0.01352173 |
| *TSPAN5* | -2.06 | 2.85E-11 |  | *MAP2* | -1.28 | 0.0130702 |
| *SMURF2* | -2.06 | 4.28E-08 |  | *PLCB4* | -1.28 | 0.041056124 |
| *RNF144B* | -2.06 | 1.78E-07 |  | *CDCA5* | -1.28 | 0.047094656 |
| *XDH* | -2.05 | 2.24E-04 |  | *TREX1* | -1.28 | 0.00037055 |
| *CLDN11* | -2.05 | 4.73E-10 |  | *TPRG1* | -1.28 | 0.033500439 |
| *TP63* | -2.05 | 5.76E-04 |  | *SBF2-AS1* | -1.27 | 0.001966593 |
| *PLCXD3* | -2.04 | 5.55E-04 |  | *LOH12CR2* | -1.27 | 0.039164129 |
| *EEPD1* | -2.04 | 0.000484007 |  | *TEF* | -1.27 | 2.10E-02 |
| *PTGIS* | -2.03 | 2.03E-07 |  | *PPP1R13B* | -1.27 | 0.000175109 |
| *NEXN* | -2.03 | 5.11E-05 |  | *CNN3* | -1.27 | 7.40905E-06 |
| *ADH1C* | -2.03 | 2.02E-04 |  | *HAVCR2* | -1.27 | 3.62E-03 |
| *RGS7BP* | -2.03 | 1.71E-02 |  | *FLJ32255* | -1.26 | 1.23E-03 |
| *LINC01750* | -2.02 | 4.17E-04 |  | *NUAK2* | -1.26 | 0.005135346 |
| *CHN2* | -2.02 | 3.55E-04 |  | *REXO5* | -1.26 | 0.006230686 |
| *PLCE1-AS1* | -2.02 | 9.17E-03 |  | *SLC29A1* | -1.26 | 0.001947304 |
| *BHLHE41* | -2.02 | 3.24E-07 |  | *CCDC89* | -1.25 | 0.049991993 |
| *S100A3* | -2.02 | 3.13E-05 |  | *GHR* | -1.25 | 4.16E-03 |
| *KCNJ16* | -2.01 | 4.30E-04 |  | *RTN4RL1* | -1.25 | 2.31E-02 |
| *ADGRG4* | -2.00 | 2.01E-03 |  | *FMO5* | -1.25 | 0.018866665 |
| *TNFSF15* | -2.00 | 1.38E-03 |  | *MID1* | -1.24 | 0.007990073 |
| *GAS6-DT* | -1.99 | 3.34E-04 |  | *ZYX* | -1.24 | 7.08E-05 |
| *CCR1* | -1.99 | 0.004019591 |  | *SYTL2* | -1.24 | 0.002821235 |
| *PLK4* | -1.99 | 0.001204689 |  | *MYLIP* | -1.24 | 0.016737966 |
| *SYNE1* | -1.99 | 6.94E-04 |  | *PLEKHA4* | -1.24 | 0.000302389 |
| *SLC1A3* | -1.96 | 1.97E-05 |  | *FGF5* | -1.24 | 0.006107435 |
| *KRT34* | -1.96 | 5.99E-03 |  | *LRIG3* | -1.24 | 0.008846616 |
| *AMOTL2* | -1.96 | 4.91E-07 |  | *TMEM187* | -1.24 | 0.010750798 |
| *NCKAP5* | -1.96 | 7.85E-04 |  | *SMAD6* | -1.24 | 0.000933863 |
| *RASSF9* | -1.96 | 2.30E-04 |  | *LAMA5* | -1.24 | 0.001489824 |
| *KIAA1755* | -1.95 | 0.001051784 |  | *BAHCC1* | -1.23 | 0.013843436 |
| *LRRC2* | -1.94 | 2.37E-04 |  | *PRKAR2A-AS1* | -1.23 | 2.32E-02 |
| *MAN1C1* | -1.93 | 3.85E-05 |  | *PLCE1* | -1.23 | 0.048127796 |
| *RHOJ* | -1.93 | 7.26E-06 |  | *VWCE* | -1.23 | 0.001738568 |
| *MID1IP1* | -1.93 | 5.87051E-05 |  | *LIMS2* | -1.22 | 1.32E-03 |
| *MSTN* | -1.93 | 4.87588E-05 |  | *LOC100287015* | -1.22 | 0.034351646 |
| *ITGB8* | -1.93 | 1.05E-03 |  | *MEGF9* | -1.22 | 0.01194108 |
| *FGD6* | -1.92 | 7.95E-07 |  | *PJVK* | -1.22 | 0.031157279 |
| *OXTR* | -1.92 | 3.71E-06 |  | *TMEM254-AS1* | -1.22 | 0.013536313 |
| *ZNF608* | -1.92 | 1.74E-04 |  | *TRIM22* | -1.22 | 1.28E-03 |
| *TRPA1* | -1.92 | 1.58E-03 |  | *PRSS12* | -1.22 | 0.001182924 |
| *APOBEC3B* | -1.91 | 1.00E-02 |  | *ADAMTSL1* | -1.22 | 1.77E-02 |
| *ARHGEF28* | -1.90 | 3.10E-04 |  | *APOL6* | -1.21 | 0.022283368 |
| *MIR221* | -1.90 | 0.006648544 |  | *ZSWIM4* | -1.21 | 0.014376447 |
| *PKD1L2* | -1.90 | 3.67E-04 |  | *CTHRC1* | -1.21 | 0.009070749 |
| *ART4* | -1.90 | 2.39E-02 |  | *LRRC20* | -1.21 | 3.41E-03 |
| *HERC6* | -1.87 | 8.76E-06 |  | *C6orf132* | -1.21 | 0.022343411 |
| *KCNQ5* | -1.87 | 1.33E-02 |  | *PRR11* | -1.21 | 0.040171054 |
| *KRT19* | -1.86 | 2.74E-02 |  | *SORBS1* | -1.21 | 0.009488813 |
| *ADRA2B* | -1.86 | 1.60E-04 |  | *SIRT4* | -1.20 | 0.025550725 |
| *TMEM37* | -1.86 | 2.44E-02 |  | *FOXM1* | -1.20 | 0.029086373 |
| *MITF* | -1.85 | 3.02E-06 |  | *SOS1* | -1.20 | 7.79E-03 |
| *SVIL* | -1.85 | 4.07E-06 |  | *PKN3* | -1.20 | 0.039194749 |
| *GNA14* | -1.85 | 3.95E-06 |  | *ESR1* | -1.20 | 0.0041932 |
| *SQOR* | -1.85 | 7.53E-14 |  | *PAIP2B* | -1.20 | 0.013833167 |
| *ADH1B* | -1.84 | 2.21E-05 |  | *CDHR3* | -1.20 | 2.23E-02 |
| *UACA* | -1.84 | 0.001204689 |  | *NMNAT2* | -1.20 | 0.000587511 |
| *METTL7B* | -1.84 | 3.09E-04 |  | *PML* | -1.19 | 4.14063E-06 |
| *DTX4* | -1.83 | 2.47E-10 |  | *SAMD4A* | -1.19 | 0.001442661 |
| *CFTR* | -1.83 | 3.18E-02 |  | *DUBR* | -1.19 | 0.013536313 |
| *LINC00968* | -1.82 | 2.52E-11 |  | *FZD6* | -1.19 | 0.00241705 |
| *NEIL3* | -1.82 | 4.59E-03 |  | *ABCG1* | -1.19 | 0.041313457 |
| *KCNE3* | -1.82 | 4.451E-05 |  | *RIN2* | -1.19 | 0.018919023 |
| *TFAP2C* | -1.81 | 3.41E-02 |  | *CBX7* | -1.19 | 0.003623638 |
| *LINC02881* | -1.81 | 2.05E-02 |  | *CCDC171* | -1.19 | 0.047176548 |
| *ANXA3* | -1.81 | 5.54E-03 |  | *CDCA8* | -1.19 | 0.048616187 |
| *CCDC15* | -1.80 | 1.38E-04 |  | *SLC46A3* | -1.19 | 0.007168819 |
| *LOXL1* | -1.80 | 1.97106E-05 |  | *CLIC3* | -1.18 | 0.02638182 |
| *NR1D2* | -1.80 | 6.01E-05 |  | *JAKMIP3* | -1.18 | 0.046666792 |
| *HTR7P1* | -1.80 | 3.20E-08 |  | *CPED1* | -1.18 | 0.002014041 |
| *TSC22D1-AS1* | -1.79 | 0.00125061 |  | *EPB41* | -1.18 | 0.034806508 |
| *RANBP3L* | -1.79 | 3.52E-03 |  | *ATP23* | -1.18 | 0.003476002 |
| *WEE1* | -1.78 | 2.07E-06 |  | *UTRN* | -1.18 | 0.029611806 |
| *PEAR1* | -1.78 | 3.8759E-05 |  | *TRAFD1* | -1.18 | 0.003491545 |
| *DSC3* | -1.78 | 0.011518381 |  | *KIAA1217* | -1.18 | 0.030059303 |
| *PDCD1LG2* | -1.78 | 0.001051784 |  | *DPYD* | -1.18 | 0.004098519 |
| *ACKR2* | -1.78 | 4.57E-03 |  | *MAMDC2* | -1.18 | 0.035544947 |
| *TMEM30B* | -1.77 | 8.67E-05 |  | *NLRP1* | -1.17 | 0.005591856 |
| *SLC16A12* | -1.77 | 2.95E-02 |  | *GPC4* | -1.17 | 0.000334115 |
| *DAPP1* | -1.76 | 3.41E-02 |  | *C11orf68* | -1.17 | 2.23E-07 |
| *LDB2* | -1.76 | 4.45E-05 |  | *ADAMTS5* | -1.17 | 0.021912983 |
| *LINC02458* | -1.76 | 1.63E-03 |  | *FAM117B* | -1.17 | 0.017898664 |
| *TLR4* | -1.75 | 4.24603E-05 |  | *SLC43A1* | -1.17 | 5.02392E-05 |
| *ADH1A* | -1.75 | 6.78E-04 |  | *EMP1* | -1.17 | 0.026524535 |
| *FAM131B* | -1.75 | 3.75E-04 |  | *PSMG3-AS1* | -1.17 | 0.010732882 |
| *RAB11FIP4* | -1.75 | 3.09E-03 |  | *RAP2B* | -1.17 | 0.000558309 |
| *N4BP2L1* | -1.74 | 5.47E-06 |  | *DNAH5* | -1.16 | 0.028251992 |
| *NUF2* | -1.74 | 0.010712201 |  | *LOC100507053* | -1.16 | 0.020053351 |
| *FMN2* | -1.74 | 6.66E-06 |  | *LINC02381* | -1.16 | 1.07609E-05 |
| *AURKB* | -1.74 | 0.026585423 |  | *ZMAT3* | -1.16 | 0.001874149 |
| *ALS2CL* | -1.73 | 6.98E-04 |  | *GLIPR2* | -1.16 | 2.87E-02 |
| *NEK10* | -1.73 | 7.07E-03 |  | *ARHGAP33* | -1.16 | 0.007218222 |
| *KLK10* | -1.72 | 0.048259514 |  | *PLEKHO2* | -1.15 | 0.002882184 |
| *FILIP1L* | -1.72 | 0.000502897 |  | *CNN2* | -1.15 | 0.000138231 |
| *IRF6* | -1.72 | 4.43E-02 |  | *CASP1* | -1.15 | 0.003770851 |
| *BUB1B* | -1.72 | 0.016791965 |  | *CROT* | -1.15 | 3.40E-02 |
| *DEPDC1* | -1.72 | 1.68E-02 |  | *H1-10-AS1* | -1.15 | 0.011367715 |
| *COLEC10* | -1.72 | 1.34E-02 |  | *TRAIP* | -1.15 | 0.04361621 |
| *FRMD6* | -1.71 | 4.82E-05 |  | *BDNF-AS* | -1.14 | 0.04139709 |
| *CALHM3* | -1.71 | 0.029086373 |  | *AR* | -1.14 | 4.18E-03 |
| *TANC1* | -1.71 | 2.19E-06 |  | *RASL11A* | -1.14 | 0.043055188 |
| *RTP4* | -1.70 | 1.13E-02 |  | *SPACA9* | -1.14 | 0.025566955 |
| *MAP3K5* | -1.70 | 8.01E-04 |  | *IL15RA* | -1.14 | 0.000194011 |
| *PCLAF* | -1.70 | 1.90E-02 |  | *TNS2* | -1.14 | 2.36E-03 |
| *XRCC2* | -1.70 | 4.02E-03 |  | *PAQR4* | -1.14 | 0.009873237 |
| *MSC* | -1.70 | 4.33122E-08 |  | *RALGPS2* | -1.14 | 0.02186635 |
| *LOC100129034* | -1.69 | 3.39E-08 |  | *FAM167B* | -1.14 | 0.041019613 |
| *ST3GAL6* | -1.69 | 4.01E-04 |  | *SLC9A3R2* | -1.13 | 0.028950436 |
| *UNC13D* | -1.69 | 3.29E-02 |  | *P2RX6* | -1.13 | 0.049160858 |
| *POLH* | -1.69 | 1.34897E-06 |  | *PPFIBP2* | -1.13 | 0.011626096 |
| *S100A14* | -1.69 | 3.90E-02 |  | *ITPKB* | -1.13 | 7.25E-04 |
| *FGF9* | -1.69 | 3.76E-02 |  | *TXNIP* | -1.13 | 0.005368055 |
| *TCF7* | -1.69 | 1.51E-04 |  | *ANKRD13A* | -1.13 | 2.68253E-06 |
| *S100A2* | -1.68 | 6.51E-03 |  | *TFEB* | -1.13 | 0.028013625 |
| *ARHGAP24* | -1.68 | 1.99773E-06 |  | *TTC30B* | -1.12 | 0.012293982 |
| *ZFHX4-AS1* | -1.68 | 3.48E-03 |  | *LPXN* | -1.12 | 0.041612552 |
| *FLNC* | -1.68 | 8.54911E-07 |  | *MAF* | -1.12 | 0.008762392 |
| *SYNE3* | -1.68 | 0.000679493 |  | *ZEB1-AS1* | -1.12 | 0.008195282 |
| *TTPA* | -1.67 | 7.44E-03 |  | *PLEKHM1* | -1.12 | 0.003245866 |
| *IL1RL1* | -1.67 | 0.002660104 |  | *HPS3* | -1.12 | 0.003369857 |
| *SLC9A9* | -1.67 | 6.00E-11 |  | *EZH1* | -1.11 | 0.001631827 |
| *GSDMC* | -1.67 | 0.006715574 |  | *THRB* | -1.11 | 0.002086398 |
| *PEAK1* | -1.67 | 0.00022436 |  | *ERMAP* | -1.11 | 0.00066108 |
| *ERBB3* | -1.66 | 4.15E-03 |  | *ACSS3* | -1.11 | 0.018070257 |
| *FHOD1* | -1.66 | 7.32E-06 |  | *UST* | -1.11 | 0.006701388 |
| *JCAD* | -1.66 | 6.24E-04 |  | *TP53INP1* | -1.11 | 0.02592986 |
| *TNFAIP8* | -1.66 | 4.03E-05 |  | *TMEM159* | -1.11 | 0.006820504 |
| *CENPU* | -1.66 | 0.001962382 |  | *OSBPL3* | -1.11 | 0.025696236 |
| *SLC45A1* | -1.66 | 2.56E-05 |  | *KIAA1614* | -1.11 | 0.004643636 |
| *RIMS1* | -1.65 | 0.000426008 |  | *NMI* | -1.11 | 0.018428611 |
| *MBP* | -1.65 | 0.000648031 |  | *ARHGEF10* | -1.11 | 0.013827638 |
| *STAT1* | -1.65 | 3.72281E-08 |  | *ABCB9* | -1.11 | 0.044771882 |
| *FAM78A* | -1.65 | 4.11E-04 |  | *C1QTNF6* | -1.10 | 0.0004466 |
| *SFN* | -1.65 | 4.42E-02 |  | *IPO5* | -1.10 | 0.006456904 |
| *EPS8L2* | -1.65 | 1.82E-06 |  | *IRF1-AS1* | -1.10 | 0.009749177 |
| *KLF5* | -1.64 | 5.13E-03 |  | *B3GNT8* | -1.10 | 0.004006088 |
| *CSF1* | -1.64 | 9.65052E-09 |  | *CARHSP1* | -1.10 | 0.005907504 |
| *KCNA1* | -1.64 | 0.001861051 |  | *PASK* | -1.10 | 0.027399712 |
| *AHNAK* | -1.64 | 9.12E-05 |  | *LYRM9* | -1.10 | 0.022321678 |
| *ACSS1* | -1.64 | 3.67E-04 |  | *LOC100507507* | -1.10 | 0.009069152 |
| *ZNF704* | -1.64 | 8.16406E-06 |  | *PBX4* | -1.10 | 4.40E-02 |
| *LINC01119* | -1.64 | 0.001134203 |  | *C21orf58* | -1.10 | 0.021769496 |
| *LINC01465* | -1.64 | 0.004057466 |  | *SLC16A5* | -1.10 | 0.007168819 |
| *GBP4* | -1.64 | 0.012773812 |  | *PARP14* | -1.10 | 0.036675939 |
| *LINC02875* | -1.64 | 2.79E-02 |  | *LOC100507516* | -1.10 | 0.017607439 |
| *INHBB* | -1.63 | 0.007895219 |  | *EFNA5* | -1.09 | 0.001397287 |
| *ANLN* | -1.63 | 1.55E-02 |  | *MANSC1* | -1.09 | 0.008357616 |
| *MSC-AS1* | -1.63 | 6.85769E-09 |  | *PARP9* | -1.09 | 0.009894995 |
| *LLGL2* | -1.63 | 1.45E-02 |  | *ZFP36* | -1.09 | 0.012440745 |
| *RBPJ* | -1.63 | 6.19E-05 |  | *ANKRD35* | -1.09 | 0.02023828 |
| *DDB2* | -1.63 | 8.87726E-06 |  | *PKD2* | -1.09 | 0.023322092 |
| *MGAM* | -1.63 | 2.22E-02 |  | *MYLK* | -1.09 | 0.000513248 |
| *PLEKHG2* | -1.62 | 1.00E-04 |  | *RAB29* | -1.09 | 0.000967277 |
| *FAM13C* | -1.62 | 0.025239406 |  | *SAMD10* | -1.09 | 0.025951551 |
| *PAG1* | -1.62 | 8.02E-09 |  | *PYROXD2* | -1.09 | 0.001141724 |
| *EVI2A* | -1.62 | 0.008070577 |  | *SNTB2* | -1.09 | 0.000323506 |
| *MGARP* | -1.62 | 1.99E-02 |  | *EBF2* | -1.08 | 0.002376276 |
| *ARVCF* | -1.62 | 6.10E-04 |  | *SMAGP* | -1.08 | 0.007644703 |
| *IL18* | -1.62 | 0.033634469 |  | *SH3BP4* | -1.08 | 0.000677029 |
| *CLDN7* | -1.62 | 0.033575877 |  | *CHDH* | -1.08 | 0.036675939 |
| *TNFSF14* | -1.62 | 3.19E-02 |  | *DHX58* | -1.08 | 0.003945097 |
| *PPARGC1B* | -1.61 | 9.60744E-05 |  | *HDAC11* | -1.08 | 0.045066307 |
| *HMGA1* | -1.61 | 1.20E-03 |  | *TMCC3* | -1.08 | 0.010249811 |
| *ERCC6* | -1.60 | 0.003139672 |  | *PLA2G6* | -1.08 | 0.000235382 |
| *CDT1* | -1.60 | 0.007789636 |  | *IKZF2* | -1.08 | 0.044705876 |
| *DIAPH3* | -1.60 | 7.65318E-05 |  | *ACTG1* | -1.08 | 0.002824174 |
| *ADCY8* | -1.60 | 0.047176548 |  | *PTPRN2* | -1.08 | 0.047266688 |
| *LAYN* | -1.60 | 1.58497E-05 |  | *STAT5A* | -1.08 | 0.020655481 |
| *DIRC1* | -1.59 | 0.025640471 |  | *AFF1-AS1* | -1.08 | 0.002586251 |
| *GINS2* | -1.59 | 1.31E-02 |  | *F2R* | -1.08 | 3.36E-02 |
| *CDK6* | -1.59 | 5.06E-03 |  | *WHAMMP2* | -1.07 | 0.040460753 |
| *DDX60* | -1.59 | 1.16E-03 |  | *SMARCA2* | -1.07 | 0.009326774 |
| *CITED2* | -1.59 | 0.000454333 |  | *PXMP4* | -1.07 | 0.004594925 |
| *PAMR1* | -1.59 | 5.27E-03 |  | *SLCO3A1* | -1.07 | 0.001464393 |
| *AMPD3* | -1.59 | 1.23E-06 |  | *FANCE* | -1.07 | 0.038091192 |
| *TK1* | -1.58 | 0.022355539 |  | *APOBEC3G* | -1.07 | 0.038957521 |
| *NTF3* | -1.58 | 0.002997452 |  | *SNX10* | -1.07 | 0.047656155 |
| *TMEM171* | -1.58 | 3.62E-03 |  | *PLS3* | -1.07 | 0.029939521 |
| *TMOD2* | -1.58 | 7.32698E-06 |  | *BCL2L1* | -1.07 | 0.000558309 |
| *PARD6B* | -1.58 | 1.25E-02 |  | *ABCC2* | -1.07 | 0.006857296 |
| *PDGFRA* | -1.58 | 1.40E-02 |  | *FANCD2* | -1.07 | 0.01101335 |
| *IKBKE* | -1.58 | 5.2467E-08 |  | *TMEM17* | -1.07 | 0.028588314 |
| *SSBP2* | -1.57 | 0.000118676 |  | *MCM2* | -1.07 | 0.005463937 |
| *BIRC3* | -1.57 | 0.006441945 |  | *PDK2* | -1.07 | 0.009303474 |
| *MCOLN3* | -1.57 | 0.040703494 |  | *SKP2* | -1.07 | 0.020875873 |
| *GPR176* | -1.57 | 3.96E-07 |  | *PPM1F-AS1* | -1.07 | 0.012267449 |
| *C4orf46* | -1.57 | 7.8708E-08 |  | *FMNL3* | -1.07 | 0.014583627 |
| *MBOAT1* | -1.56 | 0.000426008 |  | *ZNF436* | -1.07 | 0.012989618 |
| *TGFB2* | -1.56 | 5.37E-03 |  | *GASK1B* | -1.07 | 0.008331812 |
| *LINC02709* | -1.56 | 0.008762392 |  | *MAP7D3* | -1.07 | 0.020529897 |
| *CAV1* | -1.56 | 1.59404E-06 |  | *TRIM38* | -1.06 | 0.009330449 |
| *PROM2* | -1.56 | 4.33E-02 |  | *LOC100506639* | -1.06 | 0.026077007 |
| *CALCRL* | -1.56 | 0.016293237 |  | *NBPF3* | -1.06 | 0.030107131 |
| *NKX3-2* | -1.55 | 0.022333382 |  | *XPC* | -1.06 | 0.004309944 |
| *SLC2A12* | -1.55 | 1.64E-03 |  | *BOC* | -1.06 | 0.043329725 |
| *LINC01415* | -1.55 | 0.014376447 |  | *CD27-AS1* | -1.06 | 0.001953893 |
| *DAPK1* | -1.55 | 0.002835552 |  | *GNAI1* | -1.06 | 0.007569437 |
| *RAB3IL1* | -1.55 | 3.46512E-10 |  | *CARF* | -1.06 | 0.019563799 |
| *IRF1* | -1.55 | 2.67853E-05 |  | *CRIM1* | -1.06 | 0.008152088 |
| *NR3C2* | -1.55 | 1.55E-03 |  | *LINC00886* | -1.06 | 0.04594287 |
| *ROR1* | -1.55 | 0.002749552 |  | *RAB40B* | -1.06 | 0.010346993 |
| *PPM1H* | -1.55 | 4.74E-03 |  | *SEMA4F* | -1.06 | 0.016813435 |
| *MAP1A* | -1.54 | 6.21E-06 |  | *PARD3B* | -1.05 | 0.028213669 |
| *ABTB1* | -1.54 | 1.6192E-05 |  | *LINC00857* | -1.05 | 0.040460753 |
| *PSTPIP2* | -1.54 | 8.45E-06 |  | *RBM43* | -1.05 | 0.032783198 |
| *CCBE1* | -1.54 | 4.8161E-05 |  | *C1RL* | -1.05 | 0.00066639 |
| *CEP55* | -1.54 | 1.23E-02 |  | *SCAI* | -1.05 | 0.012116508 |
| *SVIL2P* | -1.54 | 1.22E-02 |  | *SOD2* | -1.05 | 0.021859908 |
| *TNFSF10* | -1.54 | 1.03E-02 |  | *SUN1* | -1.05 | 0.006881218 |
| *CCDC136* | -1.54 | 7.56E-04 |  | *PDLIM2* | -1.05 | 0.022283368 |
| *RRAS2* | -1.54 | 9.01E-06 |  | *APPL2* | -1.05 | 0.011093102 |
| *TJP2* | -1.54 | 1.03E-04 |  | *MAP4K3-DT* | -1.05 | 0.047579565 |
| *FGD4* | -1.53 | 0.001231082 |  | *KLHL3* | -1.05 | 0.046405601 |
| *HSD52* | -1.53 | 0.036924005 |  | *IGF2BP2* | -1.05 | 0.009799088 |
| *SEPSECS-AS1* | -1.53 | 1.43E-02 |  | *PCMTD2* | -1.05 | 0.028875441 |
| *VWA5A* | -1.52 | 7.56429E-05 |  | *IER5* | -1.04 | 1.27014E-05 |
| *ARID5B* | -1.52 | 0.005199369 |  | *ADGRL2* | -1.04 | 0.019561059 |
| *POLQ* | -1.52 | 0.019568522 |  | *FANCI* | -1.04 | 0.013610788 |
| *HNMT* | -1.52 | 9.03E-06 |  | *RAB9B* | -1.04 | 0.007581866 |
| *FRY* | -1.52 | 4.33E-03 |  | *SEMA4B* | -1.04 | 0.007791607 |
| *GPRC5B* | -1.52 | 1.16E-05 |  | *MELK* | -1.04 | 0.036000239 |
| *NATD1* | -1.52 | 0.002552296 |  | *ZHX3* | -1.04 | 0.005054568 |
| *GTSE1-DT* | -1.51 | 4.77E-02 |  | *SH3RF3-AS1* | -1.04 | 0.026943459 |
| *KLHL31* | -1.51 | 4.47E-02 |  | *SELENOP* | -1.04 | 0.032905791 |
| *CALHM5* | -1.51 | 1.71E-03 |  | *MARCKS* | -1.04 | 0.01430582 |
| *TBC1D2* | -1.51 | 7.07584E-07 |  | *MR1* | -1.03 | 0.004288608 |
| *PGBD3* | -1.51 | 0.000149652 |  | *SH3RF3* | -1.03 | 0.001788221 |
| *GIPC2* | -1.51 | 1.37E-02 |  | *GDPD5* | -1.03 | 0.025010994 |
| *AKR1B10* | -1.51 | 0.005217679 |  | *MINDY4* | -1.03 | 0.019176294 |
| *OCLN* | -1.51 | 1.75E-02 |  | *ORMDL3* | -1.03 | 0.014033266 |
| *TRAK1* | -1.50 | 2.93723E-06 |  | *ZNF217* | -1.03 | 0.04863123 |
| *OSR2* | -1.50 | 0.001937354 |  | *ANKEF1* | -1.03 | 0.007510285 |
| *GBP2* | -1.50 | 6.37E-06 |  | *PDGFRL* | -1.03 | 0.018803853 |
| *WNK4* | -1.50 | 3.14E-03 |  | *RECK* | -1.03 | 0.026416636 |
| *SCN8A* | -1.50 | 6.37E-06 |  | *USP49* | -1.03 | 0.029635091 |
| *VSIG10* | -1.49 | 1.08E-05 |  | *VCL* | -1.02 | 0.006242547 |
| *SLCO4C1* | -1.49 | 0.022007691 |  | *EHBP1* | -1.02 | 0.027892891 |
| *MYO1B* | -1.49 | 0.000539576 |  | *MAP3K3* | -1.02 | 0.002749196 |
| *SPEG* | -1.49 | 0.000933806 |  | *HIC1* | -1.02 | 0.049986886 |
| *AFAP1L2* | -1.49 | 0.023577329 |  | *RNF213* | -1.02 | 0.014704601 |
| *SEMA7A* | -1.49 | 2.21572E-07 |  | *RALGAPA2* | -1.02 | 0.023180036 |
| *RNF150* | -1.49 | 0.002998044 |  | *MROH6* | -1.02 | 0.016264111 |
| *GDF5* | -1.49 | 3.38E-02 |  | *TBCK* | -1.02 | 0.011526079 |
| *DTL* | -1.48 | 1.67E-02 |  | *FAM110A* | -1.02 | 0.020655481 |
| *LIFR-AS1* | -1.48 | 2.64E-03 |  | *PLXNB2* | -1.02 | 0.003092458 |
| *SLIT2* | -1.48 | 6.86E-03 |  | *BRCA1* | -1.02 | 0.006881218 |
| *FLT3LG* | -1.48 | 4.63E-04 |  | *FAM117A* | -1.01 | 0.00436925 |
| *THRSP* | -1.48 | 0.019294563 |  | *RAB3D* | -1.01 | 0.029456245 |
| *DICER1-AS1* | -1.48 | 4.48E-02 |  | *FRMD4A* | -1.01 | 0.001641937 |
| *GPR137C* | -1.48 | 0.01035086 |  | *TCEA3* | -1.01 | 0.028931282 |
| *RDH10* | -1.48 | 0.000254062 |  | *EPHA4* | -1.01 | 0.02306293 |
| *CRIM1-DT* | -1.48 | 0.028041707 |  | *GBP1* | -1.01 | 0.006991197 |
| *TXLNB* | -1.48 | 0.034866907 |  | *CCDC121* | -1.00 | 0.021116613 |
| *DPYSL2* | -1.47 | 4.30E-04 |  | *UBALD2* | -1.00 | 0.002802672 |
| *TNFRSF11B* | -1.47 | 1.15E-02 |  | *TRIM66* | -1.00 | 0.043217177 |
| *SPATA9* | -1.47 | 0.004466986 |  | *KCND3* | -1.00 | 0.009710696 |
| *EPG5* | -1.47 | 3.88E-05 |  | *OMA1* | -1.00 | 0.007912497 |
| *FBXO48* | -1.47 | 0.000400049 |  | *CAVIN1* | -1.00 | 0.001236264 |
| *YPEL3* | -1.47 | 4.95054E-06 |  | *EFNB1* | -1.00 | 0.000983144 |
| *TDO2* | -1.47 | 0.034721921 |  | *CYP2U1* | -1.00 | 0.017839215 |
| *PRR16* | -1.47 | 1.76E-02 |  | *SLC27A1* | -1.00 | 0.008897831 |
| *CHST7* | -1.46 | 5.55E-05 |  | *DAGLA* | -1.00 | 0.003831342 |
